# Testing and controlling for horizontal pleiotropy with the probabilistic Mendelian randomization in transcriptome-wide association studies

**DOI:** 10.1101/691014

**Authors:** Zhongshang Yuan, Huanhuan Zhu, Ping Zeng, Sheng Yang, Shiquan Sun, Can Yang, Jin Liu, Xiang Zhou

## Abstract

Integrating association results from both genome-wide association studies (GWASs) and expression quantitative trait locus (eQTL) mapping studies has the potential to shed light on the molecular mechanisms underlying disease etiology. Several statistical methods have been recently developed to integrate GWASs with eQTL studies in the form of transcriptome-wide association studies (TWASs). These existing methods can all be viewed as a form of two sample Mendelian randomization (MR) analysis, which has been widely applied in various GWASs for inferring the causal relationship among complex traits. Unfortunately, most existing TWAS and MR methods make an unrealistic modeling assumption and assume that instrumental variables do not exhibit horizontal pleiotropic effects. However, horizontal pleiotropic effects have been recently discovered to be wide spread across complex traits, and, as we will show here, are also wide spread across gene expression traits. Therefore, not allowing for horizontal pleiotropic effects can be overly restrictive, and, as we will be show here, can lead to a substantial inflation of test statistics and subsequently false discoveries in TWAS applications. Here, we present a probabilistic MR method, which we refer to as PMR-Egger, for testing and controlling for horizontal pleiotropic effects in TWAS applications. PMR-Egger relies on an MR likelihood framework that unifies many existing TWAS and MR methods, accommodates multiple correlated instruments, tests the causal effect of gene on trait in the presence of horizontal pleiotropy, and, with a newly developed parameter expansion version of the expectation maximization algorithm, is scalable to hundreds of thousands of individuals. With extensive simulations, we show that PMR-Egger provides calibrated type I error control for causal effect testing in the presence of horizontal pleiotropic effects, is reasonably robust for various types of horizontal pleiotropic effect mis-specifications, is more powerful than existing MR approaches, and, as a by-product, can directly test for horizontal pleiotropy. We illustrate the benefits of PMR-Egger in applications to 39 diseases and complex traits obtained from three GWASs including the UK Biobank. In these applications, we show how PMR-Egger can lead to new biological discoveries through integrative analysis.

## Introduction

Genome-wide association studies (GWASs) have identified many SNPs associated with common diseases or disease related traits. Parallel expression quantitative trait loci (eQTL) mapping studies have also identified many cis-acting SNPs associated with the expression level of nearby genes. Integrating the existing association results from both GWASs and eQTL mapping studies has the potential to shed light on the molecular mechanisms underlying disease etiology. Several statistical methods have been recently proposed to integrate GWASs with eQTL mapping studies. For example prediXcan^1^ proposes to perform a weighted SNP set test in GWAS by inferring SNP weights from eQTL studies. TWAS^2^ proposes to infer the association between gene expression and disease trait by leveraging the shared common set of cis-SNPs. SMR^3^ or GSMR^4^ directly tests the causal association between gene expression and disease trait under a Mendelian randomization (MR) framework through selecting a single instrument or multiple independent instruments. While each of these integrative methods was originally proposed to solve a different problem, all of them can be viewed as a two-sample MR method with different modeling assumptions. Because of their relationship to MR, these methods effectively attempt to identify genes causally associated with diseases or complex traits in the context of transcriptome-wide association studies (TWAS).

MR analysis is a form of instrumental variable analysis that was originally developed in the field of causal inference^5^. MR aims to determine the causal relationship between an exposure variable (e.g. gene expression) and an outcome variable (e.g. complex trait) in observational studies. MR treats SNPs as instrumental variables for the exposure variable of interest and uses these SNP instruments to estimate and test the causal effect of the exposure variable on the outcome variable. MR methods have been widely applied to investigate the causal relationship among various complex traits^6–9^, and, through a two-sample design, can be easily adapted to settings where the exposure and outcome are measured on two different sets of individuals^10, 11^. However, MR analysis for TWAS is not straightforward and requires the development of new methods that can accommodate two important features of TWAS analysis.

First, both GWASs and eQTL mapping studies collect SNPs that are in high linkage disequilibrium (LD) with each other. Traditional MR methods, such as the random effects version or the fixed effect version of the inverse variance weighted regression^12^, MR-Egger^13^, median-based regression^14^, SMR^3^, or GSMR^4^, can only make use of a single SNP instrument or multiple independent SNP instruments. Handling only independent SNPs is restrictive, as most exposure variables/molecular traits are polygenic/omni-genic and are influenced by multiple SNPs that are in potential LD with each other. As a result, incorporating multiple correlated SNPs can often help explain a greater proportion of variance in the exposure variable than using independent SNPs, and thus can help increase power and improve estimation accuracy of MR analysis^5, 15–17^. Due to the benefits of using multiple correlated instruments, most TWAS methods (e.g. PrediXcan^1^, TWAS^2^, CoMM^18^, DPR^19^, TIGAR^20^) rely on polygenic modeling priors to incorporate all cis-SNPs that are in high LD for TWAS applications. (Certainly, while the prior used in PrediXcan is polygenic, the parameter estimates obtained from PrediXcan is sparse as it uses posterior mode instead of posterior mean.) By incorporating all cis-SNPs, as we will show below, these methods can lead to substantial power improvement over standard MR approaches that use only a few independent SNPs. Unfortunately, many TWAS methods rely on a two-stage MR inference procedure: they estimate SNP effect sizes in the exposure study and plug in these estimates to the outcome study for causal effect inference. The two-stage inference procedure in MR fails to account for the uncertainty in parameter estimates in the exposure study and can often lead to biased causal effect estimates and power loss, especially in the presence of weak instruments^5, 16^. Indeed, similar to what have been observed in the MR filed, our previous study also suggests that the likelihood based inference can substantially improve power for TWAS^18^. Therefore, it is important to incorporate multiple correlated instruments in a likelihood inference framework for MR analysis in TWAS.

Second, perhaps more importantly, SNP instruments often exhibit pervasive horizontal pleiotropic effects^21^. Horizontal pleiotropy occurs when a genetic variant affects the outcome variable through pathways other than or in addition to the exposure variable^22^. Horizontal pleiotropy is in contrast to the vertical pleiotropy, which characterizes instrument effects on the outcome variable through the path of the exposure. Horizontal pleiotropy is widely distributed across the genome, affects a wide spectrum of complex traits, and can be driven by LD and extreme polygenicity of traits^21, 23^. Despite its wide prevalence, however, only a limited number of MR methods have been developed to test and control for horizontal pleiotropy; even fewer are applicable for TWAS applications. For example, some existing methods (e.g. MR-PRESSO^21^) test for horizontal pleiotropic effects without directly controlling for them. Some methods (e.g. CaMMEL^24^) control for horizontal pleiotropic effects without directly testing them^25, 26^. Some methods (e.g. Egger regression^13, 27^, GLIDE^28^, GSMR^4^, MR-median method^14^, profile score approach^29^, MRMix^30^ and Bayesian MR^31, 32^) test and control for horizontal pleiotropic effects, but can only accommodate independent instruments. As far as we are aware, there is only one two-sample MR method currently developed for testing and controlling for pleiotropic effects in the presence of correlated instruments: LDA MR-Egger^33^. Unfortunately, as we will show below, LDA MR-Egger cannot handle realistic LD pattern among cis-SNPs for TWAS applications.

Here, we develop a generative two-sample MR method in a likelihood framework, which we refer to as the probabilistic two-sample Mendelian randomization (PMR), to perform MR analysis using multiple correlated instruments for TWAS applications. We illustrate how the PMR framework can facilitate the understanding of many existing MR approaches as well as many existing integrative analysis approaches. Within the PMR framework, we focus on a particular horizontal pleiotropy effect modeling assumption based on the burden test assumption commonly used for rare variant test. This particular horizontal pleiotropy effect, as we will show later, effectively generalizes the Egger regression assumption commonly used for MR analysis to correlated instruments. Our method allows us to test the causal effect in the presence of horizontal pleiotropy, and, with a parameter expansion version of the expectation maximization algorithm (PX-EM), is scalable to hundreds of thousands of individuals. We refer to our method as PMR-Egger. With simulations, we show that PMR-Egger provides calibrated type I error for causal effect testing in the presence of horizontal pleiotropic effects, is more powerful than existing MR approaches, and, as a by-product, can directly test for horizontal pleiotropy. We apply our method to perform TWAS for 39 diseases and complex traits obtained from three GWASs with sample size ranging from 4,686 to 337,198.

## Methods

### PMR-Egger Overview

We consider a probabilistic Mendelian randomization framework for performing two-sample Mendelian randomization analysis with correlated SNP instruments. Two-sample Mendelian randomization analysis aims to estimate and test for the causal effect of an exposure on an outcome in the setting where the exposure and outcome variables are measured in two separate studies with no sample overlap. In the TWAS applications we consider here, the exposure variable is gene expression level that is measured in a gene expression study, while the outcome variable is a quantitative trait or a dichotomous disease status that is measured in a GWAS. Often times, the gene expression study and GWAS are performed on two separate samples. While we mostly focus on TWAS applications in the present study, we note that the two-sample Mendelian randomization is also commonly performed in settings where both the exposure and outcome variables are complex traits that are measured in two separate GWASs. An illustrative diagram of MR analysis is displayed in Supplementary Fig. 1.

We denote ***x*** as an *n*_1_-vector of exposure variable (i.e. gene expression measurements) that is measured on *n*_1_ individuals in the gene expression study and denote **Z**_x_ as an *n*_1_ by *p* matrix of genotypes for *p* instruments (i.e. cis-SNPs) in the same study. Note that, unlike standard MR methods that select independent instruments, we follow existing TWAS approaches and use all cis-SNPs that are in LD as instruments. We denote **y** as an *n*_2_-vector of outcome variable (i.e. trait) that is measured on *n*_2_ individuals in the GWAS and denote **Z**_*y*_ as an *n*_2_ by *p* matrix of genotypes for the same *p* instruments there. We consider three linear regressions to model the two studies separately

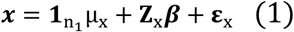

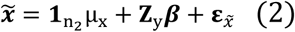

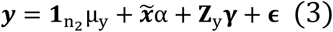

where the equation (1) is for the gene expression data and the equations (2)-(3) are for the GWAS data. Here, μ_x_ and μ_y_ are the intercepts; 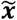 is an unobserved *n*_2_-vector of exposure variable on the *n*_2_ individuals in the GWAS; ***β*** is a *p*-vector of instrumental effect sizes on the exposure variable; α is a scalar that represents the causal effect of the exposure variable on the outcome variable; **γ** is a *p*-vector of horizontal pleiotropic effect sizes of *p* instruments on the outcome variable; ε_x_ is an *n*_1_-vector of residual error with each element independently and identically distributed from a normal distribution 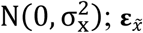 is an *n*_2_-vector of residual error with each element independently and identically distributed from the same normal distribution 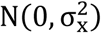; and ∈ is an *n*_2_-vector of residual error with each element independently and identically distributed from a normal distribution 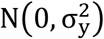. We note that while the above three equations are specified based on two separate studies, they are joined together with the common parameter ***β*** and the unobserved gene expression measurements 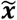. Equations (2)-(3) can also be combined into

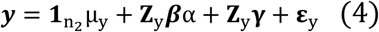

where 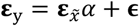.

Our key parameter of interest in the above joint model is the causal effect α. The causal interpretation of α requires two assumptions of MR analysis to hold: (i) instruments are associated with the exposure; (ii) instruments are not associated with any other confounders that may be associated with both exposure and outcome. Note that our model no longer requires the general exclusion restriction condition of traditional MR (i.e. instruments only influence the outcome through the path of exposure), as we make explicite modeling assumptions on the horizontal pleiotropy effects **γ**. Certainly, PMR-Egger still need to satisfy the InSIDE assumption that the instrument-exposure effects and instrument-outcome effects are independent of each other, which is sometimes refered to as the weak exclusion restriction condition^13^. In our model, we derive the causal interpretation and identification of α under the decision-theoretic framework of causal inference^31, 34–36^ (details in Supplementary Note). Because the causal effect interpretation of α depends on MR assumptions as well as other explicit modeling assumptions, many of which are not easily testable in practice, MR analysis in observational studies likely provides weaker causality evidence than randomized clinical trials. Therefore, while we follow standard MR analysis and use the term “causal effect” through the text, we only intend to use this term to emphasize the fact that α estimate from an MR analysis is more trustworthy than the effect size estimate in a standard linear regression of ***y*** on 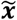.

Because *p* is often larger than *n*_1_, we will need to make additional modeling assumptions on ***β*** to make the model identifiable. In addition, the two instrumental effect terms defined in equation (4), the vertical pleiotropic effect **Z**_y_***β***α and the horizontal pleiotropic effect **Z**_y_**γ**, are also not identifiable from each other, unless we make additional modeling assumptions on **γ**. Here, we follow standard polygenic model and assume that all elements in ***β*** are non-zero and that each follows a normal distribution 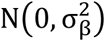. In addition, we follow the burden test assumption commonly used for rare variant test and assume that equal horizontal pleiotropic effects across SNPs **γ**_*j*_ = *γ* for *j* = 1, … *p*. With the burden test assumption on the horizontal pleiotropic effects *γ*, our model becomes a generalization of the commonly used MR-Egger regression model. In the special case where instruments are independent and treated as fixed effects and where a two-stage estimation procedure is used for inference, our model reduces to MR-Egger. However, our method can handle general cases where MR-Egger does not apply to. In particular, unlike MR-Egger, our method can handle multiple correlated instruments and perform inference in a likelihood framework.

In the above model, we are interested in estimating the causal effect *α* and testing the null hypothesis H_0_: α = 0 in the presence of horizontal pleiotropy effects **γ**. In addition, we are interested in estimating the horizontal pleiotropic effect size *γ* and testing the null hypothesis H_0_: γ = 0. We accomplish both tasks through the maximum likelihood inference framework. In particular, we develop an expectation maximization (EM) algorithm for parameter inference by maximizing the joint likelihood defined based on equations (1) and (4) (details in the Supplementary Note). The EM algorithm allows us to obtain the maximum likelihood of the joint model, together with maximum likelihood estimates for both *α* and *γ*. In addition, we apply the EM algorithm to two reduced models, one without *α* and the other without *γ*, to obtain the corresponding maximum likelihoods. Afterwards, we perform likelihood ratio tests for either H_0_: α = 0 or H_0_: γ = 0, by contrasting the maximum likelihood obtained from the joint model to that obtained from each of the two reduced models, respectively. We refer to the above inference procedure as probabilistic, as we place estimation and testing into a maximum likelihood framework. Our inference procedure is in contrast to the commonly used two-stage estimation procedure (as used in, for example, Egger regression^13, 27^, PrediXcan^1^ and TWAS^2^), which estimates 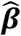 from equation (1) first and then plug in the estimates into equation (4) for inference. The previous two-stage estimation procedure fails to properly account for the estimation uncertainty in 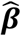 and is known to lose power compared to a formal likelihood inference procedure^5, 16, 18^.

We refer to our model and algorithm together as the two-sample probabilistic Mendelian randomization with Egger regression (PMR-Egger). As explained above, we use “probabilistic” to refer to both the data generative model and the maximum likelihood inference procedure. We use “Egger” to refer to the horizontal pleiotropic assumption on **γ** that effectively generalizes the Egger-regression assumption to correlated instruments. We also note that the joint generative Mendelian randomization model defined in equations (1) and (4) is a useful conceptual framework that unifies many existing MR methods. In particular, almost all existing MR methods are built upon the joint model, but with different modeling assumptions on ***β*** and **γ**, and with different inference procedures (Table 1). Compared with these existing MR approaches, PMR-Egger is capable of modeling multiple correlated instruments, effectively controls for horizontal pleiotropy, and places inference into a likelihood framework.

**Table 1.**
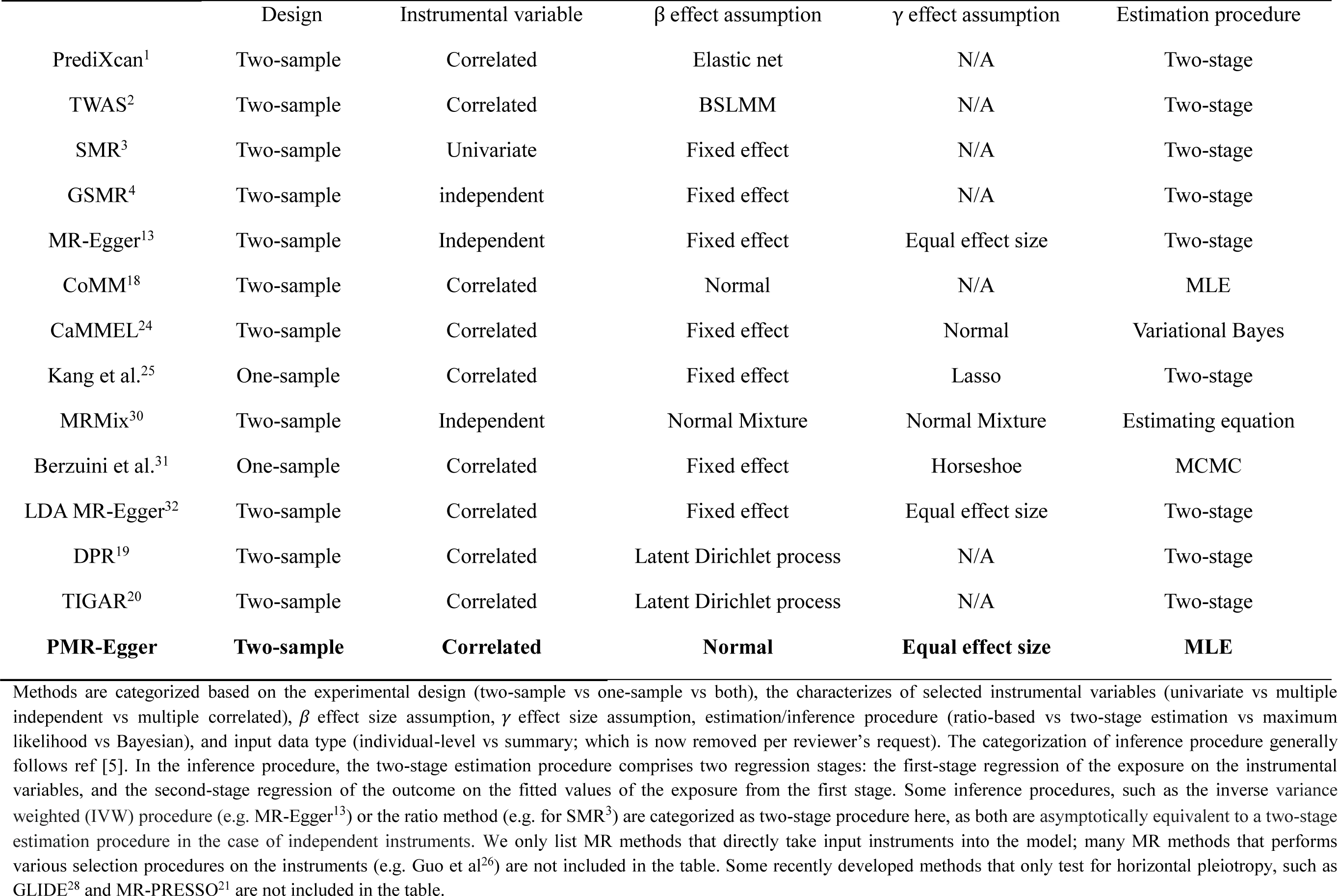
Summary of some existing MR methods

| | Design | Instrumental variable | $\beta$ effect assumption | $\gamma$ effect assumption | Estimation procedure |
| --- | --- | --- | --- | --- | --- |
| PrediXcan<sup>1</sup> | Two-sample | Correlated | Elastic net | N/A | Two-stage |
| TWAS<sup>2</sup> | Two-sample | Correlated | BSLMM | N/A | Two-stage |
| SMR<sup>3</sup> | Two-sample | Univariate | Fixed effect | N/A | Two-stage |
| GSMR<sup>4</sup> | Two-sample | independent | Fixed effect | N/A | Two-stage |
| MR-Egger<sup>13</sup> | Two-sample | Independent | Fixed effect | Equal effect size | Two-stage |
| CoMM<sup>18</sup> | Two-sample | Correlated | Normal | N/A | MLE |
| CaMMEL<sup>24</sup> | Two-sample | Correlated | Fixed effect | Normal | Variational Bayes |
| Kang et al.<sup>25</sup> | One-sample | Correlated | Fixed effect | Lasso | Two-stage |
| MRMix<sup>30</sup> | Two-sample | Independent | Normal Mixture | Normal Mixture | Estimating equation |
| Berzuini et al.<sup>31</sup> | One-sample | Correlated | Fixed effect | Horseshoe | MCMC |
| LDA MR-Egger<sup>32</sup> | Two-sample | Correlated | Fixed effect | Equal effect size | Two-stage |
| DPR<sup>19</sup> | Two-sample | Correlated | Latent Dirichlet process | N/A | Two-stage |
| TIGAR<sup>20</sup> | Two-sample | Correlated | Latent Dirichlet process | N/A | Two-stage |
| **PMR-Egger** | **Two-sample** | **Correlated** | **Normal** | **Equal effect size** | **MLE** |

### Simulations

We performed simulations to assess the performance of PMR-Egger and compare it with existing approaches. To do so, we first obtained 556 cis-SNPs for the gene *BACE1* on chromosome 11 from the GEUVADIS data^37^ (data processing details in the next section) and simulated gene expression values. We used the gene *BACE1* because the number of cis-SNPs in this gene represents the median of all genes. With the scaled genotype data ***Z***_*x*_, we simulated SNP effect sizes ***β*** from a normal distribution *N*(0, *PVE*_zx_/556), where the scalar *PVE*_zx_ represents the proportion of gene expression variance explained by genetic effects. We summed the genetic effects across all cis-SNPs as ***Z***_*x*_***β***. In addition, we simulated residual errors ε_x_ from a normal distribution *N*(0, 1 − *PVE*_zx_). We then summed the genetic effects and residual errors to yield the simulated gene expression level.

Next, we obtained genotypes for the same 556 SNPs from 2,000 randomly selected control individuals in the Kaiser Permanente/UCSF Genetic Epidemiology Research Study on Adult Health and Aging (GERA)^38, 39^ and simulated a quantitative trait. Here, we directly used ***β*** from the gene expression data, which, when paired with the causal effect *α*, yielded the vertical pleiotropic effects α***β***. We set 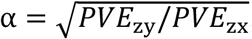, and we simulated residual errors ε_y_ from a normal distribution *N*(0, 1 − *PVE*_zy_). Here, the scalar parameter *PVE*_zy_ represents the proportion of phenotypic variance explained by vertical pleiotropic effects in the absence of horizontal pleiotropic effects. Afterwards, we simulated horizontal pleiotropic effects **γ** for these SNPs (more details below). We summed the horizontal pleiotropic effects, vertical pleiotropic effects and residual errors to yield the simulated trait.

In the simulations, we first examined a baseline simulation setting where we set *PVE*_zx_ = 10%, *PVE*_zy_ = 0, with all γ_j_ = 0. On top of the baseline setting, we varied one parameter at a time to examine the influence of various parameters. For *PVE*_zx_, we set it to be either 1%, 5% or 10%, close to the median gene expression heritability estimates across genes^40, 41^. For ***β***, we examined alternative SNP effect size distributions that deviate from the polygenic assumption in the baseline setting. Specifically, we randomly selected either 1 SNP, 1%, 10% or 100% of the SNPs to have non-zero effect, while simulated their effects from a normal distribution to explain a fixed *PVE*_zx_. For *PVE*_zy_, we varied its value to be either 0% (for null simulations), 0.2%, 0.4% or 0.6% (for power simulations). For the horizontal pleiotropy effects **γ**, we randomly assigned a fixed proportion of γ_j_ to be non-zero (proportion equals 10%, 30%, 50%, or 100%). Afterwards, we set the absolute value of non-zero γ_j_ to be the same value of γ. As a sensitivity analysis, we also randomly assigned some of their signs to be positive and some of their signs to be negative, with the ratio of positive effects to negative effects being either 1:9, 3:7, or 5:5. Here, we set γ to be 1 × 10^−4^, 5 × 10^−4^, 1 × 10^−3^ or 2 × 10^−3^, which corresponds to the 50%, 70%, 90%, 95% quantiles of horizontal pleiotropic effect estimates across all genes and all traits in the WTCCC data (more details below), respectively. For null simulations and type I error control examination, we performed 10,000 simulation replicates for each simulation scenario described above. For power calculation, for each scenario, we performed 1,000 alternative simulations together with 9,000 null simulations and calculated power based on false discovery rate (FDR).

While we applied PMR-Egger to analyze individual-level data from all simulations, we also applied PMR-Egger to analyze summary statistics in a subset of simulations to validate the implementation of the summary statistics based PMR-Egger algorithm. These results are presented in the Discussion section. Here, we considered the simulation settings with a fixed sample size (*n*_1_ = 465, *n*_2_ = 2,000), different causal effect sizes (*PVE*_*zy*_ = 0 or 0.6%) and different pleiotropy effect sizes (*γ* = 0 or 0.0005). In the analysis, we calculated the LD matrix in the eQTL data using the observed individual-level genotypes in the eQTL study. We calculated the LD matrix in the GWAS data from a reference panel. The reference panel is constructed in three different ways, by using individual-level genotypes from either all individuals in the GWAS (n=2,000), 10% of randomly selected individuals from the GWAS (n=200), or the individuals with European ancestry from the 1,000 Genomes project (n=503).

Besides the single gene-based simulations, we also conducted cross-gene simulations. Specifically, we randomly selected 10,000 genes from GEUVADIS. We extracted cis-SNPs for these 10,000 genes, obtaining a median of 576 cis-SNPs per gene (min=11; max=7,409). For each gene in turn, we used its cis-SNPs to simulate its gene expression level as described above. Afterwards, we applied different methods to analyze simulated data. The cross-gene based simulations reflect the varying LD pattern and the varying number of cis-SNPs across genes that we observe in real data, and thus are likely to be realistic than the single gene-based simulations. We performed cross-gene simulations under all simulation settings described above, including settings with varying gene expression heritability, varying genetic architectures underlying gene expression, as well as varying causal and horizontal pleiotropy effects.

### Real Data Applications

We applied our method to perform TWAS by integrating gene expression data with several GWASs. Specifically, we obtained GEUVADIS data^37^ as the gene expression data and examined 39 phenotypes from three GWASs. The three GWASs include the Wellcome trust case control study (WTCCC)^42^, the Kaiser Permanente/UCSF Genetic Epidemiology Research Study on Adult Health and Aging (GERA)^38, 39^, and the UK Biobank^43^.

The GEUVADIS data^37^ contains gene expression measurements for 465 individuals collected from five different populations that include CEPH (CEU), Finns (FIN), British (GBR), Toscani (TSI) and Yoruba (YRI). In the expression data, we only focused on protein coding genes and lincRNAs that are annotated in GENCODE (release 12)^44, 45^. Among these genes, we removed lowly expressed genes that have zero counts in at least half of the individuals to obtain a final set of 15,810 genes. We performed PEER normalization to remove confounding effects and unwanted variations following previous studies^19, 46^. Afterwards, following^19^, to remove remaining population stratification, we quantile normalized the gene expression measurements across individuals in each population to a standard normal distribution, and then further quantile normalized the gene expression measurements to a standard normal distribution across individuals from all five populations. Besides expression data, all individuals in GEUVADIS also have their genotypes sequenced in the 1,000 Genomes Project. We obtained genotype data from the 1,000 Genomes Project phase 3. We filtered out SNPs that have a Hardy-Weinberg equilibrium (HWE) p-value < 10^−4^, a genotype call rate <95%, or a minor allele frequency (MAF) <0.01. We retained a total of 7,072,917 SNPs for analysis.

The WTCCC data consists of about 14,000 cases from seven common diseases and 2,938 shared controls^42^. The diseases include type 1 diabetes (T1D; *n*=1,963), Crohn’s disease (CD; *n*=1,748), rheumatoid arthritis (RA; *n*=1,861), bipolar disorder (BD; *n*=1,868), type 2 diabetes (T2D; *n*=1,924), coronary artery disease (CAD; *n*=1,926), and hypertension (HT; *n*=1,952). We obtained quality controlled genotypes from WTCCC and initially imputed missing genotypes using BIMBAM^47^ to arrive at a total of 458,868 SNPs shared across all individuals. Afterwards, we further imputed SNPs using the 1,000 Genomes as the reference panel using SHAPEIT and IMPUTE2^48^. We filtered out SNPs that have an HWE p-value < 10^−4^, a genotype call rate <95%, or an MAF<0.01 to obtain a total of 2,793,818 imputed SNPs. For each trait in turn, we first regressed the phenotype on the top 10 genotype principal components (PCs) and obtained phenotype residuals. We then scaled the phenotype residuals to have a mean of zero and standard deviation of one and used these phenotype residuals for TWAS analysis. In addition to the main analysis that uses phenotype residuals, we also performed parallel analysis with PMR-Egger where we used the original phenotype as the outcome variable and the top 10 genotype PCs as covariates.

The GERA study consists of 61,953 individuals and 675,367genotyped SNPs. We filtered out SNPs that had a genotype calling rate below 0.95, MAF<0.01, or HWE p value<10^−4^ to yield a total of 487,609 SNPs. We phased genotypes using SHAPEIT^49^ and imputed SNPs based on the Haplotype Reference Consortium (HRC version r1.1) reference panel^50^ on the Michigan Imputation Server using Minimac3^51^. Afterwards, we further filtered out SNPs that have a HWE p-value < 10^−4^, a genotype call rate <95%, an MAF<0.01, or an imputation score<0.30 to arrive at a total of 8,385,867 SNPs that are shared across 61,953 individuals. We examined 22 diseases in GERA that include Asthma (number of cases n=10,101), Allergic Rhinitis (n=15,193), Cardiovascular Disease (CARD, n=16,431), Cancers (n=18,714), Depressive Disorder (n=7,900), Dermatophytosis (n=8,443), Type 2 Diabetes (T2D, n=7,638), Dyslipidemia (n=33,071), Hypertension (HT, n=31,044), Hemorrhoids (n=9,922), Abdominal Hernia (n=6,876), Insomnia (n=4,357), Iron Deficiency (n=2,706), Irritable Bowel Syndrome (n=3,367), Macular Degeneration (n=4,031), Osteoarthritis (n=22,062), Osteoporosis (n=5,909), Peripheral Vascular Disease (PVD, n=4,718), Peptic Ulcer (n=1,007), Psychiatric disorders (n=9408), Stress Disorders (n=4,706), and Varicose Veins (n=2,714). For each trait in turn, we first regressed the phenotype on the top 10 genotype principal components (PCs) and obtained phenotype residuals. We then scaled the phenotype residuals to have a mean of zero and standard deviation of one and used these phenotype residuals for TWAS analysis. In addition to the main analysis that uses phenotype residuals, we also performed parallel analysis with PMR-Egger where we used the original phenotype as the outcome and the top 10 genotype PCs as covariates.

The UK Biobank data consists of 487,409 individuals and 92,693,895 imputed SNPs^43^. We followed the same sample QC procedure in Neale lab (https://github.com/Nealelab/UK_Biobank_GWAS/tree/master/imputed-v2-gwas) to retain a total of 337,198 individuals of European ancestry. We filtered out SNPs with an HWE p-value < 10^−7^, a genotype call rate <95%, or an MAF<0.001 to obtain a total of 13,876,958 SNPs. We selected 10 UK Biobank quantitative traits that have a phenotyping rate > 80%, a SNP heritability > 0.2 and a low correlation among them following a previous study^52^. The 10 traits include Height (ℎ^2^ = 0.579;), Platelet count (ℎ^2^ = 0.404), Bone mineral density (ℎ^2^ = 0.401), Red blood cell count (ℎ^2^ = 0.324), FEV1-FVC ratio (ℎ^2^ = 0.313), Body mass index (BMI, ℎ^2^ = 0.308), RBC distribution width (ℎ^2^ = 0.288), Eosinophils count (ℎ^2^ = 0.277), Forced vital capacity (ℎ^2^ = 0.277), White blood cell count (ℎ^2^ = 0.272). For each trait in turn, we regressed the resulting standardized phenotypes on sex and top 10 genotype principal components (PCs) to obtain the residuals, standardized the residuals to have a mean of zero and a standard deviation of one, and finally used these scaled residuals to conduct TWAS analysis. We also performed parallel analysis with PMR-Egger by including the top 10 genotype PCs as covariates.

We combined the GEUVADIS data with each of the three GWASs for TWAS analysis. To do so, in the GEUVADIS data, for each gene in turn, we extracted cis-SNPs that are within either 100 kb upstream of the transcription start site (TSS) or 100 kb downstream of the transcription end site (TES). We overlapped these SNPs in GEUVADIS with the SNPs obtained from each of the three GWASs to obtain common sets of SNPs. The median number of the overlapped cis-SNPs between GEUVADIS and WTCCC, GERA or UK Biobank are 200, 556 or 500, respectively. Afterwards, for each pair of gene (from GEUVADIS) and trait (from GWAS) in turn, we examined the causal relationship between gene expression and trait of interest while testing and controlling for potential horizontal pleiotropic effects.

### Compared Methods

For testing the causal effect, we compared the performance of PMR-Egger with five existing methods that include: (1) SMR, which uses a single instrument and does not control for horizontal pleiotropy. For SMR, we first performed a linear regression to choose the top associated cis-SNP to be the instrumental variable. (2) PrediXcan, which uses multiple correlated instruments but does not control for horizontal pleiotropy. For PrediXcan, we used all cis-SNPs for the model and used ElasticNet implemented in the R package glmnet to obtain the coefficient estimates for the cis-SNPs. (3) TWAS, which uses multiple correlated instruments but does not control for horizontal pleiotropy. For TWAS, we used all cis-SNPs for the model and used BSLMM^53^ implemented in the GEMMA software^54^ to obtain coefficient estimates for the cis-SNPs. (4) CoMM, which uses multiple correlated instruments but does not control for horizontal pleiotropy. We used all cis-SNPs for the model and used the R package CoMM for model fitting. (5) LDA MR-Egger, which uses multiple correlated instruments and controls for horizontal pleiotropy. We used all cis-SNPs for the model and contacted the authors of LDA MR-Egger to obtain the method source code. All these methods are suitable for two-sample design and yield *p* values for testing the causal effect *α*. Note that PrediXcan, TWAS and CoMM are not originally described as an MR method but conceptually rely on the same joint MR model based on equations (1) and (4). These three methods differ in their prior assumptions on ***β***: PrediXcan relies on ElasticNet assumption; TWAS relies on BSLMM^53^ assumption; while CoMM relies on the normal prior assumption. In addition, PrediXcan and TWAS rely on a two-stage regression procedure while CoMM is based on maximum likelihood. We were unable to compare our method with either GSRM or the standard Egger regression, as both require multiple independent SNP instruments that are generally not feasible to obtain in TWAS applications.

Again, we used all cis-SNPs for methods that can make use of multiple correlated instruments (i.e. PMR-Egger, TWAS, PrediXcan, CoMM, and LDA MR Egger). We performed a linear regression to select the top associated cis-SNP as the instrumental variable for SMR, as it can only use a single instrument. In all simulations and real data applications, methods that can use either individual-level data or summary statistics (PMR-Egger, PrediXcan and TWAS) are applied using individual-level data as input to ensure their optimal performance. Methods that can only use individual-level data (CoMM) are applied using individual-level data as input. Methods that can only use summary statistics (SMR and LDA MR-Egger) are applied using summary data as input. For PMR-Egger, we used individual-level data for all main analyses and used summary data for a subset of analyses that are described in the Discussion section.

Besides the above methods, we also compared different methods to a recently published fine-mapping TWAS method, FOCUS^55^. In the FOCUS analysis, we followed^55^ and obtained a set of independent non-overlapping genomic regions termed as LD blocks from LDetect^56^. We removed genomic regions that overlap with the MHC region due to the extensive LD structure. Following^55^, we also focus our analysis on a subset of regions that harbor at least one genome-wide-significant SNP (*p* < 5 × 10^−8^; the default threshold used in FOCUS), and for each TWAS/MR method (i.e. PMR-Egger, TWAS, PrediXcan, CoMM, or SMR), also harbor at least one TWAS gene that is declared significant by the given method. We then applied FOCUS to analyze these remaining regions and identify genes that are in the 90% credible set.

For testing horizontal pleiotropic effect, we compared the performance of PMR-Egger with two existing methods that include (1) LDA MR-Egger; and (2) the global test in MR-PRESSO, which is implemented as an R package. Both these methods examine one gene at a time and output a *p* value for testing horizontal pleiotropic effects.

## Results

Our method is described in the Methods (inside the method overview subsection there), with technical details provided in the Supplementary Note. For TWAS applications, our method examines one gene at a time and estimates and tests its causal effect on a trait of interest. Our method models multiple correlated instruments, performs MR inference in a maximum likelihood inference framework, and is capable of testing and controlling for horizontal pleiotropic effects commonly encountered in TWAS. We refer to our method as the probabilistic Mendelian randomization with Egger regression (PMR-Egger), which is implemented as an R package. Our method is computationally efficient and can analyze each gene in minutes in a GWAS with a few hundred thousand individuals (Table 2).

**Table 2.** Mean computational time (in second) of various MR methods

| Trait | #SNP in the<br>exemplary gene | CoMM | PMR-Egger | TWAS | LDA MR-Egger | SMR | PrediXcan | MR-PRESSO |
| --- | --- | --- | --- | --- | --- | --- | --- | --- |
| T1D from<br>WTCCC<br>(n=4901) | 300 | 0.51(0.19) | 0.80(0.57) | 1.97(0.86) | 0.08(0.02) | 0.0003(0.0005) | 26.74(2.81) | 408.27(74.76) |
|  | 500 | 1.21(0.41) | 1.42(0.77) | 3.48(1.16) | 0.14(0.03) | 0.0004(0.0005) | 11.77(0.64) | 829.04(135.79) |
|  | 983 | 5.85(1.50) | 9.79(1.56) | 4.69(1.73) | 0.60(0.09) | 0.0004(0.0005) | 9.96(0.78) | 2023.77(260.43) |
|  | 2106 | 111.00(12.87) | 97.33(7.63) | 5.87(2.26) | 4.18(0.59) | 0.0005(0.0005) | 22.90(2.63) | 4913.22(554.47) |
| Asthma<br>from GERA<br>(n=61,953) | 300 | 1.47(0.29) | 2.06(0.22) | 2.61(1.48) | 0.05(0.02) | 0.0002(0.0004) | 33.39(3.09) | 464.64(62.18) |
|  | 500 | 1.21(0.33) | 4.21(0.81) | 2.54(0.87) | 0.09(0.03) | 0.0002(0.0004) | 11.71(0.70) | 919.66(102.83) |
|  | 1000 | 24.37(5.13) | 21.68(1.66) | 3.07(2.55) | 0.46(0.13) | 0.0002(0.0004) | 14.29(1.30) | 2275.42(263.95) |
|  | 2008 | 59.01(4.98) | 52.52(4.47) | 4.51(1.48) | 2.33(0.71) | 0.0004(0.0005) | 20.18(3.28) | 5213.73(601.46) |
| Platelet<br>Count from<br>UK<br>Biobank<br>(n=337,198) | 300 | 2.56(0.53) | 5.57(4.54) | 5.04(4.19) | 0.09(0.02) | 0.0008(0.0004) | 10.93(1.96) | 471.55(50.44) |
|  | 500 | 6.82(2.75) | 7.61(2.30) | 5.44(4.30) | 0.15(0.02) | 0.0007(0.0005) | 12.17(1.04) | 876.06(92.90) |
|  | 1052 | 24.92(6.28) | 23.59(3.21) | 5.91(4.79) | 0.81(0.09) | 0.0008(0.0004) | 16.05(2.38) | 2133.03(77.56) |
|  | 2605 | 186.14(28.45) | 178.68(16.75) | 5.37(0.73) | 8.11(1.20) | 0.0008(0.0004) | 9.89(1.74) | 6949.72(245.75) |

Computation is carried out on a single thread of a Xeon Gold 6138 CPU. The computation time is averaged across 20 replicates, with values inside parentheses denoting the standard deviation. #SNP denotes the number of cis-SNPs for four exemplary genes in each study. The computational time for MR-PRESSO is based on 10,000 permutations.

### Simulations: Testing and estimating the causal effect

We performed simulations to examine the effectiveness of our method and compared it with existing MR approaches. Simulation details are provided in the Methods. Briefly, we simulated gene expression values based on genotypes from 456 individuals in GEUVADIS and simulated phenotypes based on genotypes from 2,000 randomly selected individuals in GERA. In the simulations, we varied the genetic architecture underlying gene expression from sparse (one SNP or 1% of SNPs are causal) to polygenic (10% or 100% of SNPs are causal). We varied the proportion of SNPs exhibiting horizontal pleiotropic effects in a wide range (from 0%, 10%, 30%, 50% to 100%). We examined directional pleiotropy setting (the ratio of SNPs with negative vs positive horizontal pleiotropic effects is 0:10), approximately directional pleiotropy setting (1:9 or 3:7) and balanced pleiotropy settings (5:5). We varied the magnitude of horizontal pleiotropic effects *γ* to be either 1×10^−4^, 5×10^−4^, 1×10^−3^, or 2×10^−3^, which corresponds to the 50%, 70%, 90%, 95% percentiles of the horizontal pleiotropic effect estimate in real data. We also varied the magnitude of causal effect *α* to be either 0, 0.14, 0.2 or 0.245, which corresponds to a proportion of phenotypic variance explained by vertical pleiotropic effects (*PVE*_zy_) as 0, 0.2%, 0.4% and 0.6% respectively.

Our first set of simulations is focused on causal effect testing. Here, we compared PMR-Egger with five different methods that include SMR, PrediXcan, TWAS, CoMM, and LDA MR-Egger. We first examined type I error control of different methods under the null (*α* = 0). In the absence of horizontal pleiotropic effects, PMR-Egger, together with PrediXcan, TWAS, and CoMM, all provides calibrated type I error (Fig. 1a). Consistent with previous observations^57^, we found that SMR produces overly-conservative/deflated p-values. The deflation of SMR p-values is presumably because SMR requires the selected instrument being a true causal SNP with a large effect size, which is not always guaranteed in practice. In addition, we found that LDA MR-Egger produces inflated p-values, presumably because LDA MR-Egger makes a fixed effect assumption on **β**. The fixed effect assumption on ***β*** is not expected to work well in TWAS settings where the number of SNPs are on the same order of the sample size in the gene expression study and where the cis-SNPs are all highly correlated with each other due to LD. Such fixed effect assumption on ***β***, when paired with the two-stage inference procedure that ignores the estimation uncertainty in the first stage, makes LDA MR-Egger sensitive to the collinearity induced by SNP correlations caused by LD. Indeed, we found that the *p*-values from LDA MR-Egger are well calibrated when we followed the exact same simulation setting used in^33^, where SNP genotypes were simulated based on an autoregressive covariance matrix with a moderate correlation parameter. However, when such correlation parameter was set to be realistically high (>0.9) or if we used SNPs from real data to carry out the same set of simulations, then we observed p-value inflation from LDA MR-Egger (Supplementary Fig. 2).

**Fig. 1.**
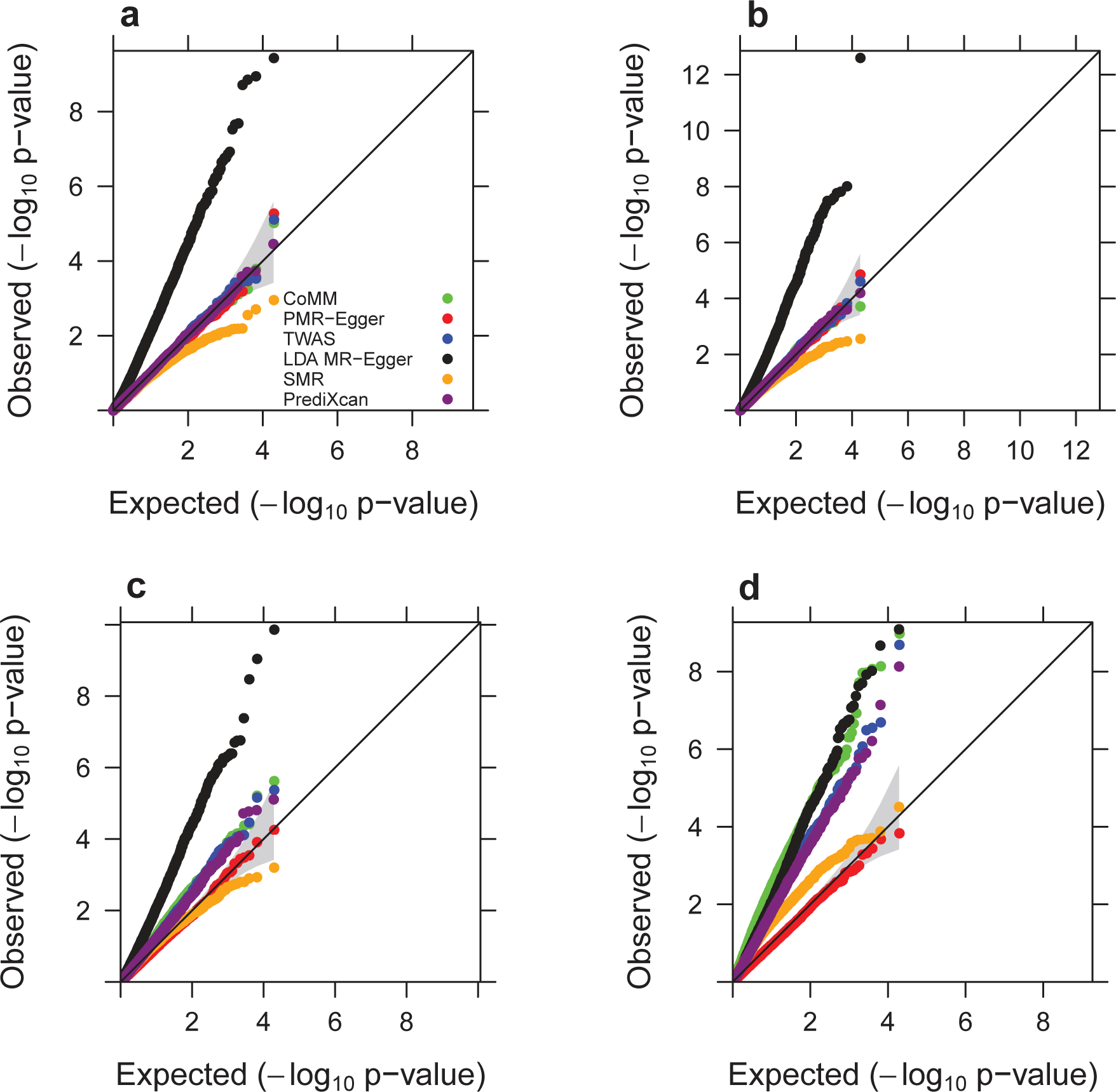
Quantile-quantile plot of −log10 p-values from different methods for testing the causal effect either in the absence or in the presence of horizontal pleiotropic effect under null simulations. Compared methods include CoMM (green), PMR-Egger (red), TWAS (blue), LDA MR-Egger (black), SMR (orange), and PrediXcan (purple). Null simulations are performed under different horizontal pleiotropic effect sizes: (a) *γ* =0; (b) *γ* =0.0001; (c) *γ* =0.0005; (d) *γ* =0.001. Only p-values from PMR-Egger adhere to the expected diagonal line across a range of horizontal pleiotropic effect sizes.

In the presence of horizontal pleiotropic effects, PMR-Egger becomes the only method that produces calibrated (or slightly conservative) p-values (Fig. 1b, c, d). In contrast, the p-values from all other methods become inflated, and more so with increasingly large horizontal pleiotropic effect. For example, when *γ* is 5×10^−4^, the genomic control factors from PMR-Egger, SMR, PrediXcan, TWAS, CoMM, and LDA MR-Egger are 0.93, 1.30, 1.33, 1.33, 1.49 and 2.61 respectively. When *γ* is increased to 1×10^−3^, the genomic control factors from PMR-Egger, SMR, PrediXcan, TWAS, CoMM, and LDA MR-Egger become 0.93, 2.39, 2.27, 2.46, 4.03 and 2.57 respectively.

The null p-value distributions from different methods remain largely similar regardless of the genetic architecture underlying gene expression being sparse or polygenic (Supplementary Fig. 3). Note that, the p-values from SMR become less deflated when there is a sparse set of SNPs affecting gene expression; however, such deflation is not completely abolished even when one SNP has non-zero effect on gene expression, presumably because we cannot always identify the true non-zero effect SNP through eQTL mapping and may supply a tagged SNP for SMR analysis. In addition, the p-value distribution pattern for different methods under the null does not change much with reduced the gene expression heritability value *PVE*_zx_. When *PVE*_zx_ is either 5% or 1%, PMR-Egger still produces well-calibrated *p* values (Supplementary Fig. 4).

We note that, like the standard MR-Egger regression, our PMR-Egger also makes a relatively strong assumption on the horizontal pleiotropic effect and assumes that all SNPs have the same horizontal pleiotropic effect. To examine the robustness of such assumption, besides the above settings where either 0% or 100% SNPs have horizontal pleiotropic effects, we varied the proportion of horizontal pleiotropic SNPs to be either 10%, 30%, 50%. We found that the p-values from PMR-Egger remain calibrated regardless of the sparsity of the horizontal pleiotropic SNPs (Supplementary Fig. 5). In addition, besides the above directional pleiotropy settings where the ratio of SNPs with negative vs positive effects is set to be 0:10, we also examined two approximately directional pleiotropy settings (1:9 or 3:7) and one balanced setting (5:5). We found that the p-values from PMR-Egger remains calibrated in either the approximately directional pleiotropy settings or the balanced setting when horizontal pleiotropic effect is small or moderate (*γ* = 1×10^−4^, 5×10^−4^, or 1×10^−3^; Supplementary Fig. 6a, b, c). However, when horizontal pleiotropic effect is large (*γ* =2×10^−3^), as one would expect, the p-values from PMR-Egger becomes inflated, with genomic control factor being 1.08, 1.31 and 1.37, for settings where the ratio is 1:9, 3:7 and 5:5, respectively (Supplementary Fig. 6d). Finally, we repeated all the above analyses with cross-gene based simulations, which provide consistent results on the type I error control of different methods for testing the causal effects (Supplementary Fig. 7-12).

Next, we examined the power of different methods to identify the causal effect for a range of possible causal effect sizes *α*. Because the same p-value from different methods may correspond to different type I errors, we computed power based on FDR of 0.1 instead of a nominal p-value threshold to allow for fair comparison across methods. In the absence of horizontal pleiotropic effects or in the presence of small horizontal pleiotropic effects, PMR-Egger, TWAS and CoMM have similarly power, all outperforming the other three methods, highlighting the importance of making polygenic assumptions on ***β*** and modeling all cis-SNPs together (Fig. 2a, b). The power of PMR-Egger is slightly lower than the other two, presumably because PMR-Egger uses extra parameters to model horizontal pleiotropy, which leads to a loss of degrees of freedom and subsequent loss of power in the absence of horizontal pleiotropy. The power of all methods increases with *α*, though their relative performance rank does not change. In the presence of horizontal pleiotropy, the power of all methods reduces (Fig. 2c, d). However, the power reduction from PMR-Egger is substantially smaller than all other methods. For example, when *PVE*_zy_ = 0.006 and *γ* = 0.0005, PMR-Egger reaches a power of 41%; the power of SMR, PrediXcan, TWAS, CoMM, and LDA MR-Egger are 7%, 24%, 31%, 33% and 1%, respectively. When *PVE*_zy_ = 0.006 but *γ* = 0.001, the power of PMR-Egger remains similar and is 40%; the power of SMR, PrediXcan, TWAS, CoMM, and LDAMR-Egger reduces to 3%, 13%, 16%, 16% and 0.9%, respectively. Besides the horizontal pleiotropic effects **γ**, we examined how power is influenced by the genetic architecture underlying gene expression, ***β*** (Supplementary Fig. 13). We found that the power of different methods in the setting where 10% of SNPs have non-zero effects on gene expression are similar to the baseline setting where all SNPs have non-zero effects, both in the absence (Supplementary Fig. 13e vs Fig. 2a) or in the presence of horizontal pleiotropic effects (Supplementary Fig. 13F vs Fig. 2d). However, the relative performance of different methods changes when there is only one SNP or 1% SNPs having non-zero effect on gene expression. Specifically, in the absence of horizontal pleiotropic effects, the power of both PrediXcan and SMR become slightly higher than PMR-Egger, TWAS and CoMM, all of which have substantially higher power than LDA MR-Egger (Supplementary Fig. 13a, c). The higher power of PrediXcan and SMR in the sparse setting presumably is because the ElasticNet estimation procedure employed in PrediXcan favors sparse eQTLs while SMR explicitly makes a single eQTL assumption. In the presence of horizontal pleiotropic effects, however, PMR-Egger remains the most powerful, even in the setting where only one SNP has non-zero effect on gene expression (Supplementary Fig. 13b, d). We also found that PMR-Egger produces accurate estimate of the causal effect *α*, both under the null and under various alternatives, in the presence or absence of horizontal pleiotropic effects (Supplementary Fig. 14). The causal effect estimates remain reasonably unbiased in the two approximately directional pleiotropy settings and one balanced setting (Supplementary Fig. 15a, c, e). Finally, we repeated all the above analyses in cross-gene based simulations, which provide consistent results on the power of different methods for detecting causal effects (Supplementary Fig. 16-17).

**Fig. 2.**
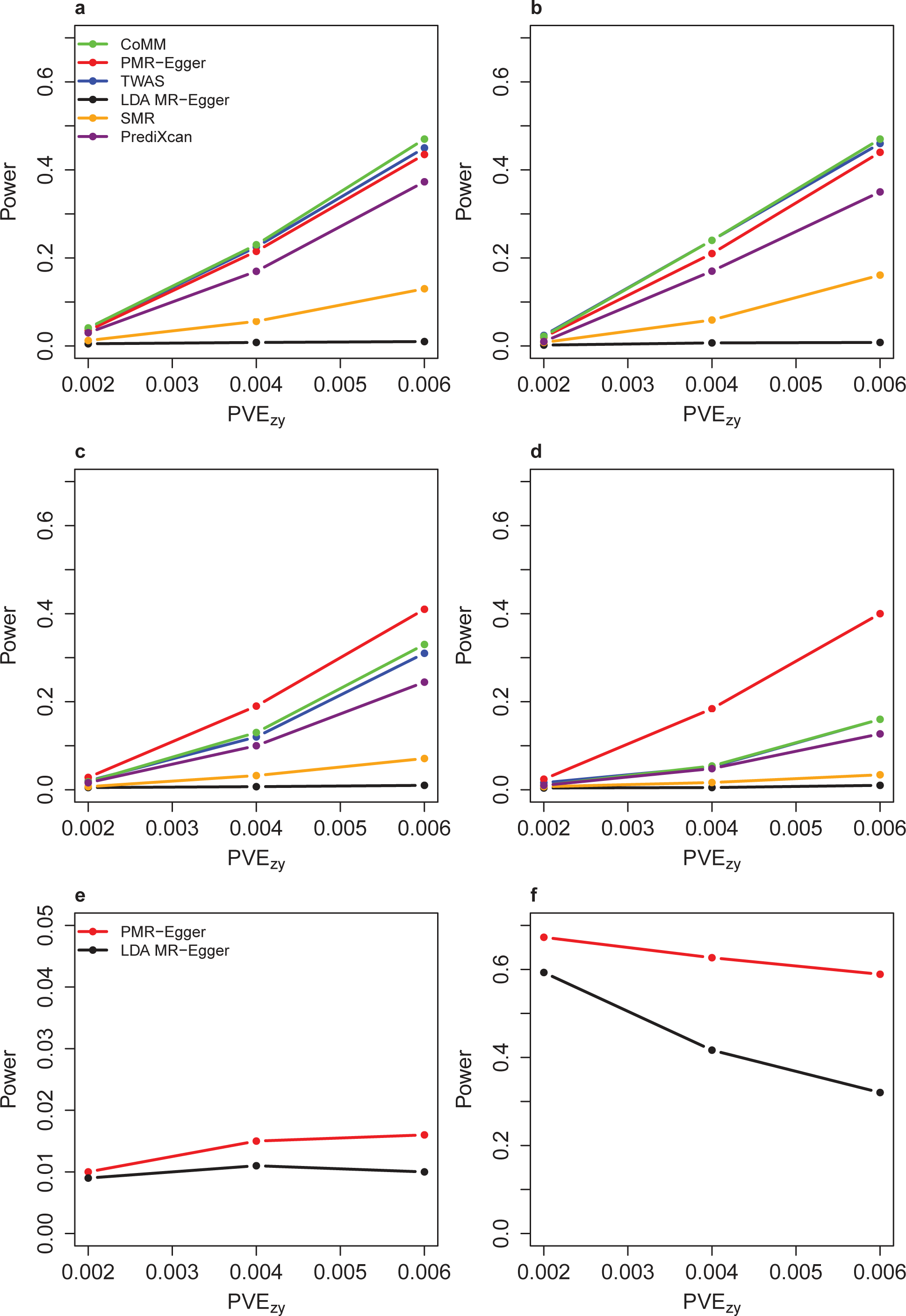
Power of different methods under various simulation scenarios. Power (y-axis) at a false discovery rate of 0.1 to detect the causal effect (a-d) or the horizontal pleiotropic effect (e-f) is plotted against different causal effect size characterized by PVE_zy (x-axis). Compared methods include CoMM (green), PMR-Egger (red), TWAS (blue), LDA MR-Egger (black), SMR (orange), and PrediXcan (purple). Simulations are performed under different horizontal pleiotropic effect sizes: (a) γ=0; (b) γ=0.0001; (c, e) γ=0.0005; (d, f) γ=0.001.

**Fig. 3.**
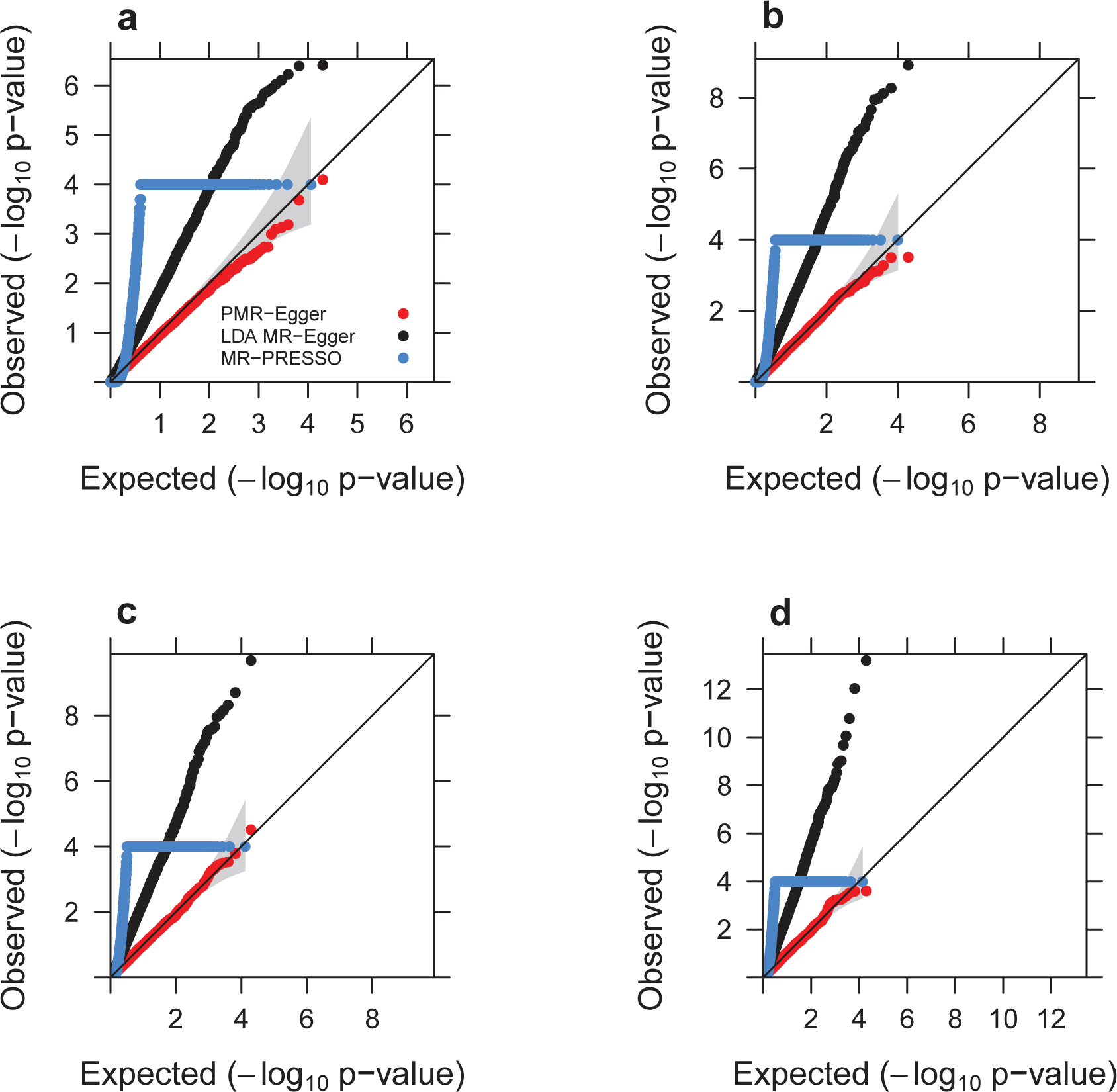
Quantile-quantile plot of −log10 p-values from different methods for testing the horizontal pleiotropic effect either in the absence or in the presence of causal effect under null simulations. Compared methods include PMR-Egger (red), LDA MR-Egger (black), and MR-PRESSO (dodger blue). Null simulations are performed under different causal effect sizes characterized by PVE_zy_: **(a)** PVE_zy_=0; **(b)** PVE_zy_=0.2%; **(c)** PVE_zy_=0.4%; and **(d)** PVE_zy_=0.6%. Only p-values from PMR-Egger adhere to the expected diagonal line across a range of horizontal pleiotropic effect sizes. Due to heavy computational burden, we are only able to run 10,000 permutations for MR-PRESSO. Therefore, the minimal p-value from MR-PRESSO is 10^−4^.

**Fig. 4.**
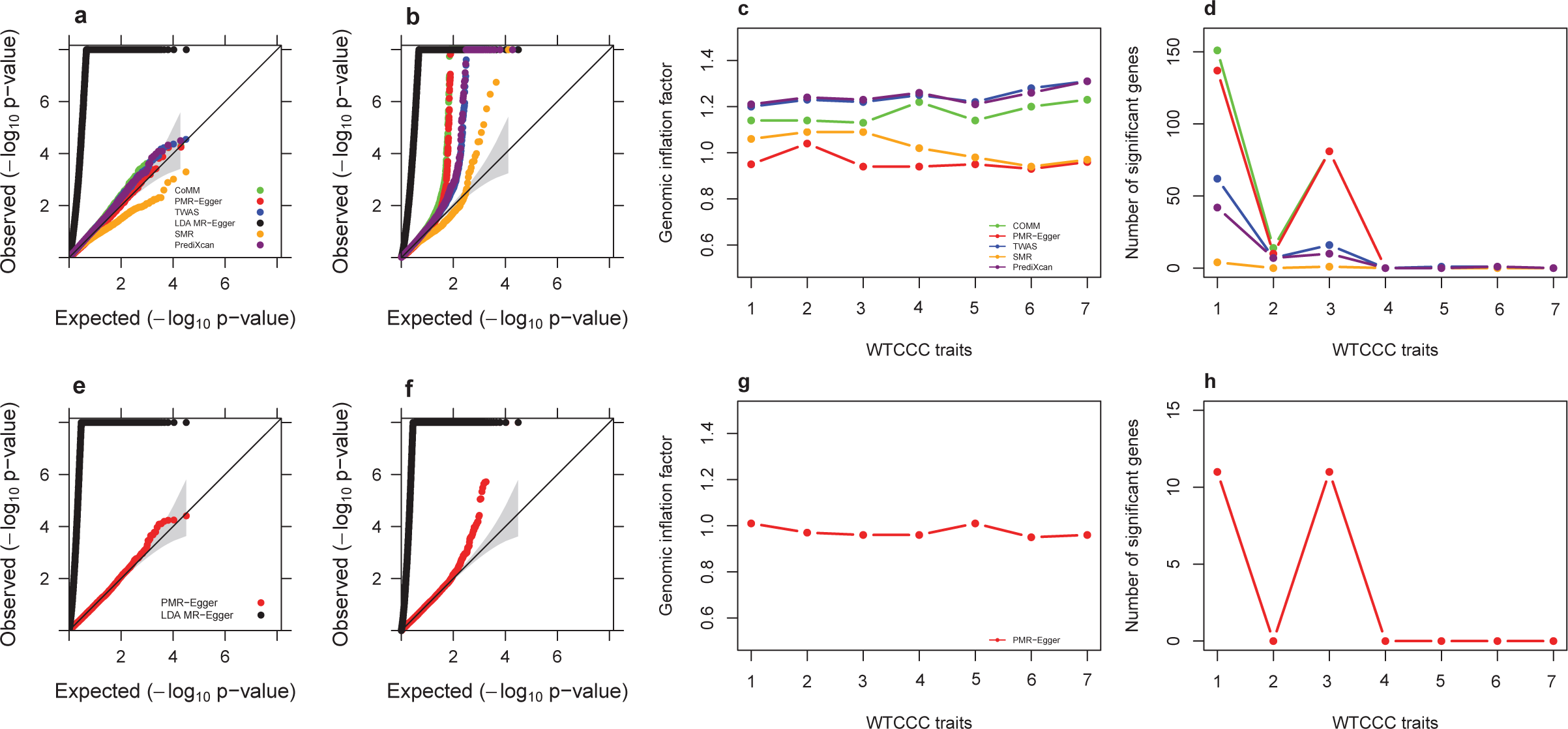
TWAS analysis results by different methods for traits in the WTCCC data. Compared methods include CoMM (green), PMR-Egger (red), TWAS (blue), LDA MR-Egger (black), SMR (orange), and PrediXcan (purple). **(a)** Quantile-quantile plot of −log10 p-values from different methods for testing the causal effect for an exemplary trait BD. **(b)** Quantile-quantile plot of −log10 p-values from different methods for testing the causal effect for another exemplary trait T1D. **(c)** Genomic inflation factor for testing the causal effect for each of the 7 traits by different methods. **(d)** Number of causal genes identified for each of the 7 traits by different methods. **(e)** Quantile-quantile plot of −log10 p-values from different methods for testing the horizontal pleiotropic effect for an exemplary trait BD. **(f)** Quantile-quantile plot of −log10 p-values from different methods for testing the horizontal pleiotropic effect for another exemplary trait T1D. **(g)** Genomic inflation factor for testing the horizontal pleiotropic effect for each of the 7 traits by different methods. **(h)** Number of genes identified to have significant horizontal pleiotropic effect for each of the 7 traits by different methods. For c, d, g, h, the number on the x-axis represents seven traits in order: T1D, CD, RA, BD, T2D, CAD, HT.

**Fig. 5.**
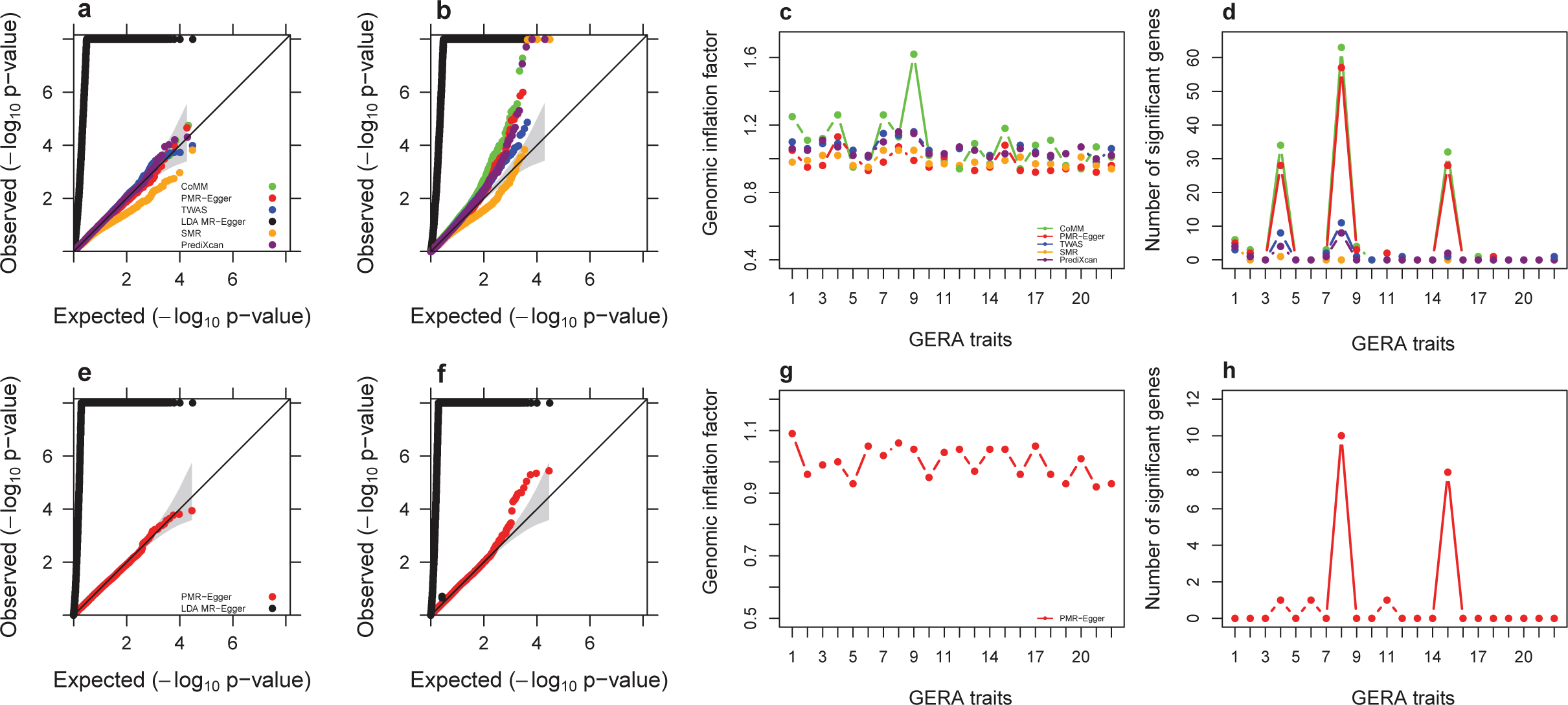
TWAS analysis results by different methods for traits in the GERA data. Compared methods include CoMM (green), PMR-Egger (red), TWAS (blue), LDA MR-Egger (black), SMR (orange), and PrediXcan (purple). **(a)** Quantile-quantile plot of −log10 p-values from different methods for testing the causal effect for an exemplary trait Irritable Bowel Syndrome. **(b)** Quantile-quantile plot of −log10 p-values from different methods for testing the causal effect for another exemplary trait Asthma. **(c)** Genomic inflation factor for testing the causal effect for each of the 22 traits by different methods. (d) Number of causal genes identified for each of the 22 traits by different methods. **(e)** Quantile-quantile plot of −log10 p-values from different methods for testing the horizontal pleiotropic effect for an exemplary trait Irritable Bowel Syndrome. **(f)** Quantile-quantile plot of −log10 p-values from different methods for testing the horizontal pleiotropic effect for another exemplary trait Asthma. **(g)** Genomic inflation factor for testing the horizontal pleiotropic effect for each of the 22 traits by different methods. **(h)** Number of genes identified to have significant horizontal pleiotropic effect for each of the 22 traits by different methods. For c, d, g, h, the number on the x-axis represents 22 traits in order: Asthma, Allergic Rhinitis, CARD, Cancers, Depressive Disorder, Dermatophytosis, T2D, Dyslipidemia, HT, Hemorrhoids, Abdominal Hernia, Insomnia, Iron Deficiency, Irritable Bowel Syndrome, Macular Degeneration, Osteoarthritis, Osteoporosis, PVD, Peptic Ulcer, Psychiatric disorders, Stress Disorders, Varicose Veins.

**Fig. 6.**
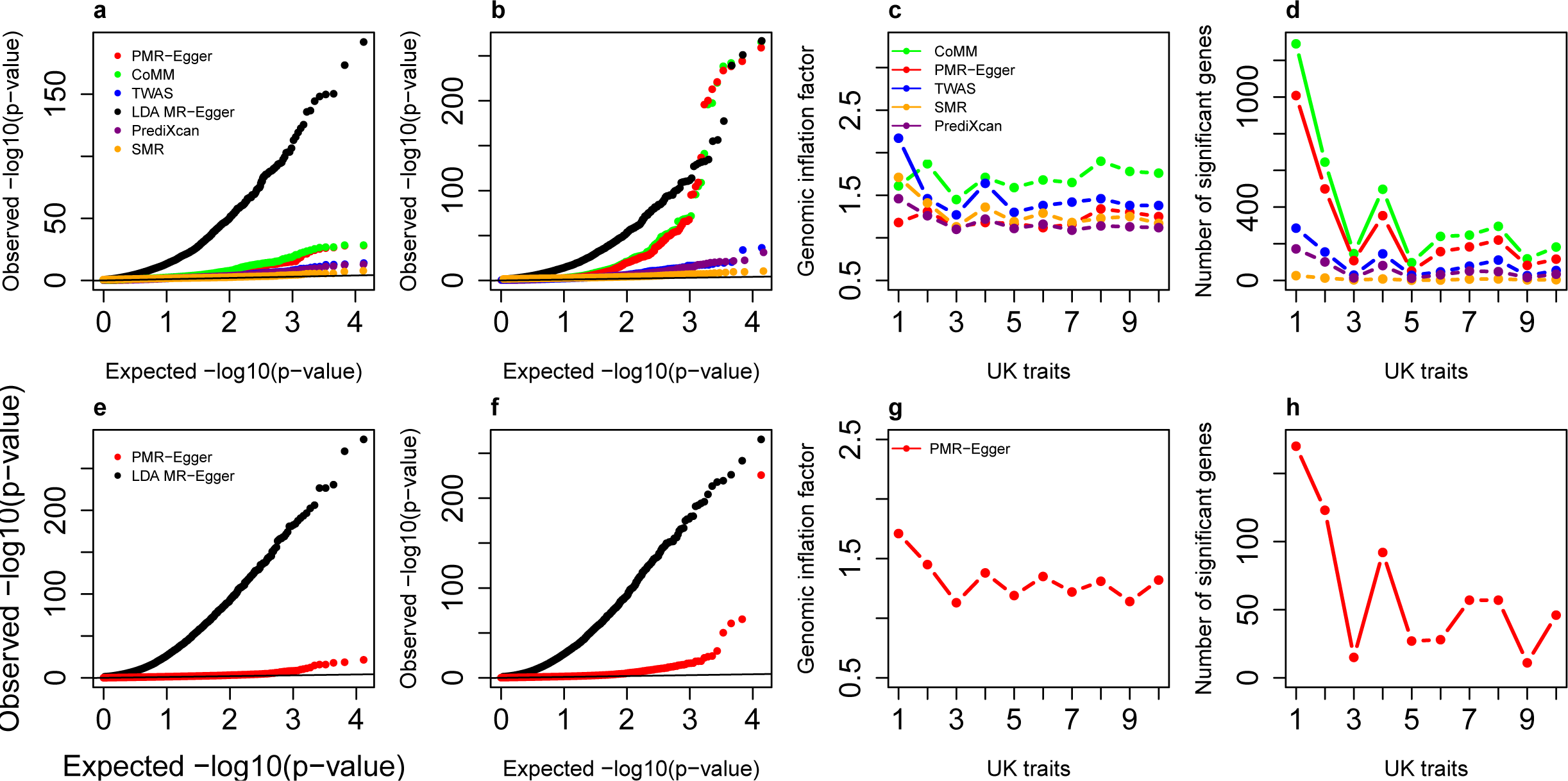
TWAS analysis results by different methods for traits in the UK Biobank data. Compared methods include CoMM (green), PMR-Egger (red), TWAS (blue), LDA MR-Egger (black), SMR (orange), and PrediXcan (purple). **(a)** Quantile-quantile plot of −log10 p-values from different methods for testing the causal effect for an exemplary trait BMI. **(b)** Quantile-quantile plot of −log10 p-values from different methods for testing the causal effect for another exemplary trait Platelet Count. **(c)** Genomic inflation factor for testing the causal effect for each of the 10 traits by different methods. **(d)** Number of causal genes identified for each of the 10 traits by different methods. **(e)** Quantile-quantile plot of −log10 p-values from different methods for testing the horizontal pleiotropic effect for an exemplary trait BMI. **(f)** Quantile-quantile plot of −log10 p-values from different methods for testing the horizontal pleiotropic effect for another exemplary trait Platelet Count. **(g)** Genomic inflation factor for testing the horizontal pleiotropic effect for each of the 10 traits by different methods. **(h)** Number of genes identified to have significant horizontal pleiotropic effect for each of the 10 traits by different methods. For c, d, g, h, the number on the x-axis represents 10 traits in order: Height, Platelet count, Bone mineral density, Red blood cell count, FEV1-FVC ratio, BMI, RDW, Eosinophils count, Forced vital capacity, White blood cell count.

### Simulations: Testing and estimating horizontal pleiotropic effect

Our second set of simulations is focused on horizontal pleiotropic effect testing. Here, we compared PMR-Egger with two different methods: LDA MR-Egger and MR-PRESSO. All three methods examine one gene at a time and test whether cis-SNPs within the gene exhibit non-zero horizontal pleiotropic effects. Note that, unlike PMR-Egger and LDA MR-Egger, MR-PRESSO requires independent instruments and uses permutation to obtain the empirical p-values. Due to the heavy computational burden resulting from permutations, we restricted the number of permutations in MR-PRESSO to 10,000 (the lowest possible p value from MR-PRESSO is thus 10^−4^) and were only able to apply MR-PRESSO to a subset of simulation scenarios.

We first examined type I error control of different methods under the null, where there is no horizontal pleiotropic effect. We found that the p-values from PMR-Egger provide calibrated type I error control under a range of causal effect sizes *α* (Fig. 3). However, p-values from both LDA MR-Egger and MR-PRESSO are inflated, and more so with increasingly large causal effect *α*. For example, when PVE_zy_ = 0, the genomic control factor from PMR-Egger and LDA MR-Egger are 0.96 and 2.31, respectively. When PVE_zy_ is increased to 0.6%, the genomic control factor from PMR-Egger remains 0.96, while the genomic control factor from LDA MR-Egger becomes 3.04. (We are unable to accurately compute the genomic control factor for MR-PRESSO because its minimal p-value is 10^−4^.) The overly inflated p-values from LDA MR-Egger is presumably due to its fixed effect modeling assumption on ***β*** and the subsequent failure to control for realistic LD patterns. The inflation of MR-PRESSO p values is presumably because MR-PRESSO can only handle independent instruments and thus does not fare well in TWAS settings. Inflation of p-value on testing horizontal pleiotropy would incorrectly identify genes with no pleiotropic effects, thus likely reducing the power to detect true causal effect *α*. Importantly, the p-values from PMR-Egger remain calibrated regardless of the genetic architecture underlying gene expression (Supplementary Fig. 18). Finally, we repeated all the above analyses in cross-gene based simulations, which provide consistent results on the type I error control of different methods for testing pleiotropic effects (Supplementary Fig. 19-20).

Next, we examined the power of different methods in detecting non-zero horizontal pleiotropic effect. Again, we computed power based on an FDR of 0.1 instead of the nominal p-value to allow for fair comparison across methods. We dropped MR-PRESSO for comparison here due to its heavy computational burden. We found that PMR-Egger outperforms LDA MR-Egger in a range of possible horizontal pleiotropic effect sizes, and that the power of both methods increases with increasing horizontal pleiotropy (Fig. 2e, f). For example, when PVE_zy_ = 0.6% and *γ* = 0.0005, PMR-Egger achieves a power of 1.6% while LDA MR-Egger achieves a power of 1% (note that the power is relatively small due to the small sample size used in the simulations). When PVE_zy_ = 0.6% but *γ* = 0.001, the power of PMR-Egger increases to 58.9% while the power of LDA MR-Egger increases to 32%. In addition, the power to detect horizontal pleiotropic effects is not influenced by the sparsity level of the genetic architecture underlying gene expression (Supplementary Fig. 21). The power to detect horizontal pleiotropic effects does, however, depend on the sparsity level of **γ** (Supplementary Fig. 22a). Specifically, power of both PMR-Egger and LDA MR-Egger reduces with increasing sparsity of **γ**, though the power of PMR-Egger remains higher than LDA MR-Egger across a range of sparsity values. Similarly, the power to detect pleiotropic effects also suffers in the absence of directional pleiotropic effect (Supplementary Fig. 22b). In addition, PMR-Egger can estimate the horizontal pleiotropic effect size accurately in the presence of directional pleiotropic effect (Supplementary Fig. 23). However, in the absence of directional pleiotropic effect, as one would expect, the estimates of pleiotropic effects become under-ward biased, more so in the balanced setting than in the approximately directional pleiotropy settings (Supplementary Fig. 15b, d, f). Finally, we repeated all the above analyses in cross-gene based simulations, which provide consistent results on the power of different methods for detecting pleiotropic effects (Supplementary Fig. 24-25).

### Real data applications

We performed TWAS to detect genes causally associated with any of the 39 phenotypes collected from three GWASs (details in Methods). The examined gene expression data is obtained from the GEUVADIS study and contains 15,810 genes. The examined phenotypes include 7 common diseases from WTCCC, 22 diseases from GERA, and 10 quantitative traits from UK Biobank. The GWAS sample size ranges from 4,686 (for Crohn’s disease in WTCCC) to 337,198 (for UK Biobank). We applied PMR-Egger together with five other approaches (SMR, PrediXcan, TWAS, CoMM, and LDA MR-Egger) to examine pairs of gene and phenotype one at a time. In the analysis, we regressed phenotypes on the top 10 genotyping PCs to obtain the phenotype residuals, which we used further to conduct TWAS analysis for all compared methods. The p-values for testing the causal effect of each gene on the phenotype are shown for WTCCC traits (Fig. 4a, b and Supplementary Fig. 26), GERA traits (Fig. 5a, b and Supplementary Fig. 27), and UK Biobank traits (Fig. 6a, b and Supplementary Fig. 28); with genomic control factors listed in Supplementary Table 1 and visualized in Fig. 4c, Fig. 5c and Fig. 6c. Besides these main analyses, we also performed parallel analysis for PMR-Egger where we used the original phenotype as the outcome and included the top 10 genotype PCs as covariates (Supplementary Figures 29-31). The results from these parallel analyses are largely consistent with the main results. Therefore, we will mainly report the main results in the following text. For illustration purpose, we display qq-plots for two selected traits in each data, one with a relatively low number of gene associations and the other with a relatively high number of gene associations, in Fig. 4a, b, Fig. 5a, b and Fig. 6a, b, respectively. Among the selected six traits, the one with zero number of associated genes (BD in WTCCC; Fig. 4a) and the one with one associated gene (Irritable Bowel Syndrome in GERA; Fig. 5a), represent approximately null traits with no apparently associated genes. For the six selected traits, consistent with simulations, we found that the p-values from PMR-Egger are well calibrated, more so than the other methods. In contrast, the p-values from CoMM, TWAS, PrediXcan and LDA MR-Egger are inflated and deviated upward from the diagonal line, while the p-values from SMR are overly conservative and lie below the diagonal line. The results observed in these exemplary traits generalize to all other examined traits. For example, the genomic control factor from PMR-Egger is the lowest among all methods in 25 out of the 39 traits, and ranges from 0.93 to 1.04 in WTCCC (Fig. 4c), from 0.92 to 1.13 in GERA (Fig. 5c), and from 1.12 to 1.34 in UK Biobank (Fig. 6c). (Note that the higher genomic control factor in the large UK Biobank as compared to WTCCC and GERA is expected under polygenic architecture^58^ and reflects at least in part the higher power in the UK Biobank as compared to GERA and WTCCC.) In contrast, the genomic control factors from CoMM, TWAS, PrediXcan are often higher than that from PMR-Egger for most traits examined. For example, the genomic control factor from CoMM is often the highest among all other methods (except for LDA MR-Egger) in 22 out of the 39 traits, and ranges from 1.13 to 1.23 in WTCCC, 0.94 to 1.62 in GERA, and 1.45 to 1.90 in UK Biobank. The genomic control factor from TWAS is the highest among all other methods (except for LDA MR-Egger) in 14 out of the 39 traits, ranges from 1.20 to 1.31 in WTCCC, 0.98 to 1.15 in GERA, and 1.30 to 2.17 in UK Biobank. The genomic control factor from PrediXcan is the highest among all other methods (except for LDA MR-Egger) in 5 out of the 39 traits, and ranges from 1.21 to 1.31 in WTCCC, 1.00 to 1.16 in GERA, and 1.09 to 1.46 in UK Biobank. In addition, consistent with simulations, we observed a substantial inflation of LDA MR-Egger p-values: its genomic control factor ranges from 17.60 to 18.56 in WTCCC, 32.13 to 34.74 in GERA, and 10.48 to 16.65 in UK Biobank. Also consistent with simulations, the p-value from SMR often lies underneath the expected null, even though its genomic control factors are often well behaved (Fig. 4a, b, Fig. 5a, b, Fig. 6a, b and Supplementary Figs. 26-28).

We examined the number of associated genes detected by different methods based on a Bonferroni corrected genome-wide threshold (Fig. 4d, Fig. 5d and Fig. 6d; Supplementary Table 2). We note that the number of detected genes based on this p-value threshold may artificially favors those methods that have inflated type I error control. For this analysis, we excluded LDA MR-Egger for comparison, as its p-values are overly inflated. Comparing across the remaining methods, we found that SMR can barely detect any genes significantly associated with traits across all three data sets, much less so than that detected by the other four methods. The much lower number of genes detected by SMR than the other four methods are consistent with the relatively low power of SMR observed in simulations. For the other four methods, we found that the number of gene-trait pairs detected by CoMM and PMR-Egger is higher than that detected by TWAS and PrediXcan in all three GWASs (Fig. 4d, Fig. 5d and Fig. 6d; Supplementary Table 2). The higher number of discoveries by both CoMM and PMR-Egger in the three GWASs is consistent with our simulations as well as previous observations that likelihood-based inference often achieves higher power than two-stage inference for MR analysis. However, we do notice that PMR-Egger detects slightly lower number of gene-trait pairs than CoMM based on the same genome-wide p-value threshold, consistent with the inflated genomic inflation factors observed for CoMM. Indeed, we found that the estimated 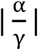 for the common set of genes detected by both CoMM and PMR-Egger is higher than the set of genes only detected by CoMM across traits (Supplementary Fig. 32a, b). Therefore, the genes detected by CoMM but not PMR-Egger tend to have large |γ| and small |α|, likely reflecting false associations due to horizontal pleiotropic confounding.

Overall, by controlling for horizontal pleiotropic effects, PMR-Egger detected many likely causal genes that the other methods failed to detect. For example, the *LNK*/*SH2B3* gene (111,743,752-111,989,427 on chr 12) is only identified by PMR-Egger to be associated with platelet count in the UK Biobank (PMR-Egger *p* = 1.17 × 10^−221^; CoMM *p*=0.98; TWAS *p* = 8.6 × 10^−5^; PrediXcan *p*=0.68; SMR *p*=0.024). The association between *LNK* and plate count is consistent with results from recent large-scale GWASs^59–61^. *LNK*/*SH2B3* encodes the lymphocyte adaptor protein (LNK) that is primarily expressed in hematopoietic and endothelial cells^62^. In hematopoietic cells, LNK functions as a negative regulator of cell proliferation as well as the thrombopoietin-mediated cytokine signaling pathway, which is a key signaling pathway that promotes megakaryocytes to form platelets^62, 63^. Indeed, platelets are overproduced and accumulated in *LNK* knockdown cells as well as *Lnk* knockout mouse^64–66^, supporting a causal role of *LNK* in platelets production. As the second example, the *NOD2* gene (50,627,514-50,866,988 on chr 16) is identified by PMR-Egger to be associated with Crohn’s disease (CD; *p* = 6.1 × 10^−19^), and, with a slightly less significance, also by CoMM (*p* = 7.8 × 10^−15^). The association between *NOD2* and CD was not identified by the other methods (TWAS *p*=0.005; PrediXcan *p*=0.92; SMR *p*=0.15). *NOD2* encodes a cytosolic pattern recognition receptor that acts both as a cytoplasmic sensor of microbial products and as an important mediator of innate immunity and inflammatory response^67^ The *NOD2* gene is a well-known susceptible gene for CD and is perhaps one of the first genes ever implied for CD^68^. Multiple SNPs in *NOD2* have been found to be associated with CD in both early linkage studies^69–71^ and many recent GWASs^72, 73^. *NOD2* variants associated with CD often reside in the ligand recognition domain of NOD2 and can lead to aberrant bacterial handling and antigen presentation^74^. Indeed, *NOD2*-deficient mice displays dysregulated bacterial community in the ileum and *NOD2*-deficient ileal epithelia exhibit impaired ability of inducing immune responses for bacteria elimination^75^. It is thus hypothesized that mis-regulation of *NOD2* can causally lead to altered interactions between ileal microbiota and mucosal immunity, resulting in increased disease susceptibility to CD^75^. As a third example, the *TFRC* gene (195,654,054-195,909,060 on chr 3) is identified by PMR-Egger to be associated with red blood cell distribution width (RDW) in the UK Biobank (*p* = 3.3 × 10^−17^). Such association is not identified by the other methods (CoMM *p*=0.95; TWAS *p*=0.76; PrediXcan *p*=0.97; SMR *p*=0.38). *TFRC* encodes the classical transferrin receptor that is involved in cellular iron uptake^76, 77^. Multiple SNPs in *TFRC* have been established to be associated with various erythrocyte phenotypes in GWASs^78, 79^. These associated erythrocyte phenotypes include the mean corpuscular hemoglobin (MCH) and mean corpuscular volume (MCV, the average volume of red blood cells) which is directly related to RDW^77, 78^. The variants in *TFRC* likely lead to decreased iron availability for red cell precursors, as has been observed in mice deficient in *TFRC*, thus resulting in a compensatory increase of red blood cell size as measured by RDW^80^. The regional association plots for all these three genes are presented in the Supplementary Fig.33-35.

We compared the results from different MR methods with a recently published TWAS fine-mapping method, FOCUS^55^. The analysis details are provided in the Materials and Methods section. Briefly, we follow ^55^ and focused on independent and non-overlapping genomic regions that harbor at least one genome-wide-significant SNP and at least one TWAS gene that is significant by the MR methods. The number of genes and regions analyzed by FOCUS for each of the three data sets are shown in Supplementary Table 3, which also contains the number of associated genes detected by FOCUS in the credible set. Due to the small number of associated genes detected in WTCCC, we focus our main comparison in GERA and UK Biobank. In these real data applications, we found that the results from PMR-Egger is largely consistent with that of FOCUS, more so than the other methods (Supplementary Fig. 41). Specifically, the average PMR-Egger −log10(p-value) for genes in the FOCUS 90% credible set is 22.43 in GERA and 10.67 in UK Biobank. The average −log10(p-value) of PMR-Egger is higher than CoMM (13.83 and 10.43), TWAS (5.71 and 7.55), PrediXcan (4.66 and 7.06) and SMR (NA for GERA, as no gene in the credible set is detected by SMR; 1.78 for UKbiobank). In addition, the difference of the average PMR-Egger −log10(p-value) between genes in the FOCUS credible set and genes outside is large (16.61 in GERA and 7.43 in UK Biobank). The −log10(p-value) difference is again larger than CoMM (8.41 and 6.02), TWAS (4.52 and 5.35), PrediXcan (3.50 and 4.74) and SMR (NA and 0.28). Similarly, the proportion of significant genes detected by PMR-Egger in the FOCUS credible set is 78% in GERA and 60% in UK Biobank. The proportion of significant genes by PMR-Egger is higher than CoMM (75% and 53%), TWAS (50% and 47%), PrediXcan (50% and 48%) and SMR (NA and 8%). In addition, the difference in the proportion of significant genes detected by PMR-Egger between genes in the FOCUS credible set and genes outside is high (53% in GERA and 41% in UK Biobank). This proportion difference by PMR-Egger is again higher than CoMM (51% and 39%), TWAS (46% and 36%), PrediXcan (50% and 35%) and SMR (NA and 1%). The consistency between PMR-Egger and FOCUS validates the high power of PMR-Egger.

Next, we shift our focus to testing horizontal pleiotropic effects. The p-values for testing the causal effect of gene on phenotype are shown for WTCCC traits (Fig. 4e, f and Supplementary Fig. 26), GERA traits (Fig. 5e, f and Supplementary Fig. 36), and UK Biobank traits (Fig. 6e, f and Supplementary Fig. 37); with genomic control factors visualized in Fig. 4g, Fig. 5g and Fig. 6g. We also display qq-plots for the previously selected exemplary traits in Fig. 4e, f, Fig. 5e, f, and Fig. 6e, F. Overall, consistent with simulations, the p-values from PMR-Egger are well behaved while the p-value from LDA MR-Egger display substantial inflation. For example, the genomic control factor from PMR-Egger ranges from 0.93 to 1.01 in WTCCC (Fig. 4g), from 0.92 to 1.09 in GERA (Fig. 5g), and from 1.13 to 1.71 in UK Biobank (Fig. 6g). In contrast, the genomic control factor from LDA MR-Egger ranges from 34.00 to 36.00 in WTCCC, from 69.82 to 72.19 in GERA and from 17.75 to 29.85 in UK Biobank (Supplementary Table 1). With the same Bonferroni adjusted genome-wide p-value threshold, PMR-Egger detected 33 gene-trait pairs in WTCCC in which the cis-SNPs exhibit significant horizontal pleiotropy, 37 gene-trait pairs in GERA, and 626 gene-trait pairs in the UK Biobank.

Horizontal pleiotropic effect tests can help us explain some of the discrepancy in terms of the causal associations detected by PMR-Egger and the other methods. For example, for the trait of red blood cell count in UK Biobank, the *MAPT* gene on chromosome 17 shows a significant pleiotropy effect (*p* = 2.35 × 10^−9^) but displays no significant causal effect (*p*=0.98) by PMR-Egger. In contrast, *MAPT* is detected to be significantly associated with red blood cell count by PrediXcan (*p* = 8.11 × 10^−10^), and, to a much lesser extent, by TWAS (*p* = 1.72 × 10^−3^). However, no previous evidence suggests that *MAPT* is associated with red blood cell count. Indeed, we found that the genomic location of *MAPT* (43,871,748-44,205,700) is close to and partially overlapped with *KANSL1* (44,007,282-44,402,733), which has been previously identified to be associated with red blood cell traits^81, 82^. The association between *KANSL1* and red blood cell count is also detected by PMR-Egger (*p* = 1.02 × 10^−7^), by CoMM (*p* = 2.72 × 10^−8^), and, to a much lesser extent, by TWAS (*p* = 1.66 × 10^−3^) in the present study. By controlling for the expression level of the *KANSL1* gene in the PrediXcan framework, the association between the predicted *MAPT* expression level and red blood cell count is no longer significant (*p* = 0.10). Therefore, the causal association between *MAPT* and red blood cell count detected by PrediXcan likely reflects either the true horizontal pleiotropic effect of *MAPT* cis-SNPs on red blood cell count through *KANSL1* or their tagging effects of the neighboring eQTLs of *KANSL1*. As another example, for height in the UK Biobank, the pseudogene *RP11-9E13.2* (70,137,755-70,340,521) on chromosome 10 has a significant pleiotropy effect (*p* = 1.08 × 10^−13^) but displays no significant causal effect (*p*=0.93) by PMR-Egger. In contrast, *RP11-9E13.2* is detected to be significantly associated with height by PrediXcan (*p* = 4.34 × 10^−10^), and, to a lesser extent, by TWAS (*p* = 9.05 × 10^−6^). The pseudogene *RP11-9E13.2* is in the neighborhood of *MYPN* (69,765,912-70,071,774), which has been previously identified to be associated with height^83^. The association between *MYPN* and height is also detected by PMR-Egger (*p* = 1.82 × 10^−7^), CoMM (*p* = 2.13 × 10^−14^), and to a lesser extent, PrediXcan (*p* = 3.94 × 10^−4^) and TWAS (*p* = 1.55 × 10^−3^), in the present study. By controlling for the predicted expression level of *MYPN* gene in the PrediXcan framework, the association between the predicted *RP11-9E13.2* expression level and height is no longer significant at the genome-wide threshold (*p* = 3.37 × 10^−4^). Therefore, the causal association between the pseudogene *RP11-9E13.2* and height as detected by PrediXcan and TWAS likely reflects either the horizontal pleiotropic effect of *RP11-9E13.2* cis-SNPs on height through *MYPN* or their tagging effects of the neighboring eQTLs of *MYPN*. The results suggest the practical importance of testing and controlling for pleiotropic effects in TWAS applications. Certainly, we acknowledge that, both these examples are focused on the special case where the false gene association with the trait disappears when conditional on a neighboring gene. We did not provide examples where the apparently false gene association with the trait may be explained by horizontal pleiotropic effects acted upon/through a gene far away, as it is often challenging to convincingly identify trans eQTL effects. In the special case we focused on, while it is possible that SNPs display true horizontal pleiotropic effects through the neighboring gene, it is equally likely that SNPs used in the model are simply tagging nearby eQTLs of the neighboring causal gene^55, 84^ and thus display apparent “horizontal pleiotropic effects” through the neighboring gene, as also mentioned above. Subsequently, the horizontal pleiotropic effect term in PMR-Egger may represent the apparent “horizontal pleiotropic effects” through SNP tagging to the nearby eQTLs of the causal gene, rather than the truly horizontal pleiotropic effect acted through other molecular pathways. Regardless of the interpretation of the horizontal pleiotropic effect term, we found it reassuring that by modeling the horizontal pleiotropic effect term in PMR-Egger can reduce false discoveries in the case of SNP tagging.

We note that an important feature of PMR-Egger is its ability to test both causal effect and horizontal pleiotropy effect simultaneously. We contrast the p-values obtained from these two different tests across genes for those traits in which at least one gene is detected as significant from either of the two tests (Supplementary Figs. 38-40). We found that different traits exhibit different gene association patterns. For example, some traits may only contain genes with a significant causal effect but without a significant horizontal pleiotropic effect (e.g. CD and CAD in WTCCC; Allergic Rhinitis, Irritable Bowel Syndrome and Psychiatric disorders in GERA). Some traits may only contain genes with a significant horizontal pleiotropic effect but without a significant causal effect (e.g. Dermatophytosis in GERA). Some traits may contain genes with a significant causal effect as well as genes with a significant horizontal pleiotropic effect, but with the two sets of genes being non-overlapped (e.g. Asthma, Dyslipidemia, HT, Abdominal Hernia and Macular Degeneration in GERA; Fored Vitral Capacity in UK Biobank). While the majority of traits contain genes with both a significant causal effect and a significant horizontal pleiotropic effect. The top gene which is most significant for both causal effect test and pleiotropy test is highlighted in the plots. Being capable of testing both causal effect and horizontal pleiotropy effect facilitates our understanding of the gene association pattern with various different complex traits.

## Discussion

We have presented a data generative model and a likelihood framework for MR analysis that unifies many existing transcriptome wide association analysis methods and many existing MR methods. Under the framework, we have presented PMR-Egger, a new method that conducts MR analysis using multiple correlated instruments while properly controlling for horizontal pleiotropic effects. By properly controlling for horizontal pleiotropic effects and making inference under a likelihood framework, PMR-Egger yields calibrated p-values across a wide range of scenarios and improves power of MR analysis over existing approaches. We have illustrated the benefits of PMR-Egger through extensive simulations and multiple real data applications of TWAS.

One important modeling assumption we made in PMR-Egger is that the horizontal pleiotropic effects of all SNPs equal to each other. The equal effect size assumption directly follows the commonly used Egger regression modeling assumption for MR analysis and is analogous to the burden effect size assumption commonly used for rare variant tests. Consistent with existing literature on applications of the Egger regression and burden test, we also found that equal effect size assumption employed in PMR-Egger works reasonably robust for causal effect estimation and testing with respect to a range of model mis-specifications and appears to be effective in several real data applications examined here. However, we do acknowledge that our equal effect size assumption in PMR-Egger can be overly restrictive in many settings. For example, as described in the Results, in the absence of direction pleiotropy, the pleiotropic effect estimate becomes down-ward biased and the pleiotropic effect test loses power. We have attempted to alleviate this restrictive modeling assumption by imposing an alternative modeling assumption on the horizontal effect sizes based on variance component assumption. In particular, we have attempted to assume that the horizontal pleiotropic effect of each SNP follows a normal distribution with mean zero and a certain variance component parameter, i.e. analogous to the SKAT test assumption^85^. Such variance component assumption is a more flexible modeling assumption than the equal effect size assumption, potentially alleviating much of the concern with respect to the sensitivity and robustness of equal effect size assumption. Unfortunately, under the variance component assumption, inference for the resulting PMR model becomes overly complicated. In particular, due to the estimation uncertainty in the hyper-parameter estimates, the p-values from the PMR variance component model becomes severely deflated even under simple null simulations (Supplementary Fig. S42). Such deflation of p-values has been previously observed in variance component tests for microbiome applications^86^. Only few methods exist to address such p-value in-calibration issue resulting from hyper-parameter estimation uncertainty^87^, and it is not straightforward to adapt any of these methods to our PMR variance component model. Besides the equal effect size modeling restriction, we also note that neither PMR-Egger nor the PMR variance component model is capable of accounting for correlation between horizontal pleiotropic effects *γ* and the SNP effects on gene expression *β*. Therefore, while we view PMR-Egger as in important first step towards effective control of horizontal pleiotropic effects in TWAS applications, we emphasize that imposing more realistic modeling assumptions on the horizontal pleiotropic effects in the PMR framework will likely yield more fruitful results in the future.

We have primarily focused on modeling continuous traits with PMR-Egger. For case control studies, we have followed previous approaches and directly treated binary phenotypes as continuous outcomes^19, 53, 88, 89^, which appears to work well in both WTCCC and GERA data applications we examined. Treating binary phenotypes as continuous outcomes can be justified by recognizing the linear model as a first order Taylor approximation to a generalized linear model^53^. However, it would be desirable to extend PMR-Egger to accommodate case control data or other discrete data types in a principled way, by, for example, extending PMR-Egger into the generalized linear model framework. In particular, we could use a probit or a logistic link to extend PMR-Egger to directly model case control data. Extending PMR-Egger to model discrete data types using the generalized linear model framework would likely lead to wider applications of PMR-Egger and is thus an important avenue for future research.

We have primarily focused on modeling individual-level data with PMR-Egger. However, like many other linear model-based methods in statistical genetics, PMR-Egger can also be easily extended to make use of summary statistics. The summary statistics version of PMR-Egger is described in detail in the Supplementary Text. Briefly, the summary statistics version of PMR-Egger requires marginal SNP effect size estimates and their standard errors, both on the gene expression and on the trait of interest. In addition, it requires a SNP by SNP correlation matrix that can be constructed based on a reference panel. We validated the implementation of the summary statistics-based approach of PMR-Egger in simulations (details in Materials and Methods). In the comparison, we constructed the SNP by SNP correlation matrix from three different reference panels, by using either all individuals from the GWAS data, 10% randomly selected individuals from the GWAS data, or individuals of European ancestry from the 1,000 Genomes project. We applied the summary statistics-based approach of PMR-Egger to each reference panel and compared results with the individual level data-based approach of PMR-Egger that was applied to the complete data. The p values from both approaches for testing causal effects as well as for testing pleiotropy effects are largely consistent with each other, demonstrating the effectiveness of summary statistics-based approach of PMR-Egger (Supplementary Fig. 43). The summary statistics-based approach of PMR-Egger is implemented in the same software package. Being able to make use of summary statistics extends the applicability of PMR-Egger to data sets where individual-level genotype or phenotype are not available.

Finally, in addition to what we have already mentioned in the Materials and Methods, we emphasize here again, that, while we have followed the previous MR literature and use “causal effect” through the text, the effect is causal only when certain MR modeling assumptions hold. These MR assumptions are often not straightforward to prove. For example, without measuring all potential confounders, it is not straightforward to argue that the SNP instruments are not associated with any other confounders that may be associated with both exposure and outcome. Therefore, we caution against the over-interpretation of causal inference in observation studies such as TWAS applications. However, we do believe MR is an important step that allows us to move beyond standard linear regressions and is an important analysis that can provide potentially more trustworthy evidence with regard to causality compared to simpler approaches.

## Code availability

Our method is implemented in the R package PMR, freely available at http://www.xzlab.org/software.html and https://cran.r-roject.org/web/packages/PPMR/index.html. The code to reproduce all the analyses are available on GitHub (https://github.com/yuanzhongshang/PMRreproduce).

## Data availability

No data were generated in the present study. The GEUVADIS gene expression data is publicly available at http://www.geuvadis.org. The WTCCC genotype and phenotype data is publicly available at https://www.wtccc.org.uk. The GERA genotype and phenotype data is available in dbGaP (https://www.ncbi.nlm.nih.gov/gap) with accession number phs000788. The UK Biobank data is from UK Biobank resource under Application Number 30686.

## Acknowledgements

This study was supported by the National Institutes of Health (NIH) grants R01HG009124 and R01HL142023, National Science Foundation (NSF) grant DMS1712933, and the National Natural Science Foundation of China (81872712 and 81673272). We thank the Wellcome Trust Centre for Human Genetics for making the heterogenous stock mouse data available online. This study makes use of data generated by the Wellcome Trust Case Control Consortium (WTCCC). A full list of the investigators who contributed to the generation of the data is available from http://www.wtccc.org.uk/. Funding for the WTCCC project was provided by the Wellcome Trust under award 076113 and 085475. The GERA Data (dbGaP accession number phs000788) came from a grant, the Resource for Genetic Epidemiology Research in Adult Health and Aging (RC2 AG033067; Schaefer and Risch, PIs) awarded to the Kaiser Permanente Research Program on Genes, Environment, and Health (RPGEH) and the UCSF Institute for Human Genetics. The RPGEH was supported by grants from the Robert Wood Johnson Foundation, the Wayne and Gladys Valley Foundation, the Ellison Medical Foundation, Kaiser Permanente Northern California, and the Kaiser Permanente National and Northern California Community Benefit Programs. The RPGEH and the Resource for Genetic Epidemiology Research in Adult Health and Aging are described in the following publication, Schaefer C, et al., The Kaiser Permanente Research Program on Genes, Environment and Health: Development of a Research Resource in a Multi-Ethnic Health Plan with Electronic Medical Records, in preparation, 2013. This study has been conducted using UK Biobank resource under Application Number 30686. UK Biobank was established by the Wellcome Trust medical charity, Medical Research Council, Department of Health, Scottish Government and the Northwest Regional Development Agency. It has also had funding from the Welsh Assembly Government, British Heart Foundation and Diabetes UK.

## Author contributions

XZ conceived the idea and provided funding support. XZ and ZY developed the methods. ZY developed the software tool with assistance from JL and CY. ZY performed simulations and real data analysis with assistance from HZ, PZ, SY, and SS. XZ and ZY wrote the manuscript with input from all other authors. All authors reviewed and approved the final manuscript.

