## Supplemental files for "Testing and controlling for horizontal pleiotropy with the probabilistic Mendelian randomization in transcriptome-wide association studies"

### 1. Model and likelihood

The proposed model is

$$\boldsymbol{x} = \mu_x + \mathbf{Z}_x\boldsymbol{\beta} + \boldsymbol{\varepsilon}_x \qquad (1)$$

$$\boldsymbol{y} = \mu_y + \mathbf{Z}_y\boldsymbol{\beta}\alpha + \mathbf{Z}_y\boldsymbol{\gamma} + \boldsymbol{\varepsilon}_y \quad (2)$$

Where the equation (1) is for the gene expression data and the equation (2) is for the GWAS data. Here, $\mu_x$ and $\mu_y$ are the intercepts for the two models, respectively; $\boldsymbol{\beta}$ is a $p$-vector of instrumental effect sizes on the explanatory variable; $\alpha$ is a scalar that represents the causal effect of the explanatory variable on the outcome variable; $\boldsymbol{\gamma}$ is a $p$-vector of horizontal pleiotropic effect sizes of $p$ instruments on the outcome variable; $\boldsymbol{\varepsilon}_x$ is an $n_1$-vector of residual error with each element independently and identically distributed from a normal distribution $N(0, \sigma_x^2)$;and $\boldsymbol{\varepsilon}_y$ is an $n_2$-vector of residual error with each element independently and identically distributed from a normal distribution $N(0, \sigma_y^2)$. We note that while the above two equations are specified based on two separate studies, they are joined together with the common parameter $\boldsymbol{\beta}$. We assume $\boldsymbol{\beta} \sim N(0, \sigma_z^2 I_p)$.

Given $\boldsymbol{\beta}$, we have $\boldsymbol{x} \sim N(\mu_x + \mathrm{Z}_x\boldsymbol{\beta}, \sigma_x^2 I_{n_1}), \boldsymbol{y} \sim N\left(\mu_y + \mathbf{Z}_y\boldsymbol{\beta}\alpha + \mathbf{Z}_y\boldsymbol{\gamma}, \sigma_y^2 I_{n_2}\right)$

Given $\mathbf{Z}_x, \boldsymbol{Z}_y$, the observed data are $(\boldsymbol{x}, \mathbf{y})$ and the observed likelihood function is

$$f(\boldsymbol{x}, \mathbf{y}) = \int f(\boldsymbol{x}, \boldsymbol{y}, \boldsymbol{\beta})d\boldsymbol{\beta} = \int f(\boldsymbol{x}, \mathbf{y}|\boldsymbol{\beta})f(\boldsymbol{\beta})d\boldsymbol{\beta} = \int f(\mathbf{y}|\boldsymbol{\beta})f(\boldsymbol{x}|\boldsymbol{\beta})f(\boldsymbol{\beta})d\boldsymbol{\beta}$$

the last equality is due to that given $\boldsymbol{\beta}$, $\boldsymbol{y}$ and $\boldsymbol{x}$ are independent.

While $f(\boldsymbol{y}|\boldsymbol{\beta})f(\boldsymbol{x}|\boldsymbol{\beta})f(\boldsymbol{\beta}) = (2\pi)^{-\frac{n_2}{2}}(\sigma_y^2)^{-\frac{n_2}{2}}\exp\left(-\frac{(\boldsymbol{y}-\mu_y-\alpha\mathbf{Z}_y\boldsymbol{\beta}-\mathrm{Z}_y\boldsymbol{\gamma})^T(\boldsymbol{y}-\mu_y-\alpha\mathbf{Z}_y\boldsymbol{\beta}-\mathbf{Z}_y\boldsymbol{\gamma})}{2\sigma_y^2}\right)\cdot$

$$(2\pi)^{-\frac{n_1}{2}}(\sigma_x^2)^{-\frac{n_1}{2}}\exp\left(-\frac{(\boldsymbol{x}-\mu_x-\mathbf{Z}_x\boldsymbol{\beta})^T(\boldsymbol{x}-\mu_x-\mathbf{Z}_x\boldsymbol{\beta})}{2\sigma_x^2}\right)(2\pi)^{-\frac{p}{2}}(\sigma_z^2)^{-\frac{p}{2}}exp\left(\frac{\boldsymbol{\beta}^T\boldsymbol{\beta}}{2\sigma_z^2}\right)$$

$$= C \cdot exp\left(-\frac{(\boldsymbol{y}-\mu_y-\mathbf{Z}_y\boldsymbol{\gamma})^T(\boldsymbol{y}-\mu_y-\mathbf{Z}_y\boldsymbol{\gamma})}{2\sigma_y^2}\right)exp\left(-\frac{(\boldsymbol{x}-\mu_x)^T(\boldsymbol{x}-\mu_x)}{2\sigma_x^2}\right)$$

$$\cdot\, exp\left(-\frac{\boldsymbol{\beta}^T K\boldsymbol{\beta} - 2\left(\frac{(\boldsymbol{y}-\mu_y-\mathbf{Z}_y\boldsymbol{\gamma})^T\alpha\mathbf{Z}_y}{\sigma_y^2} + \frac{(\boldsymbol{x}-\mu_x)^T\mathbf{Z}_x}{\sigma_x^2}\right)\boldsymbol{\beta}}{2}\right)$$

Where $C = (2\pi)^{-\frac{n_2}{2}}(\sigma_y^2)^{-\frac{n_2}{2}}(2\pi)^{-\frac{n_1}{2}}(\sigma_x^2)^{-\frac{n_1}{2}}(2\pi)^{-\frac{p}{2}}(\sigma_z^2)^{-\frac{p}{2}}$,

$$K_{p\times p} = \frac{\alpha^2\mathbf{Z}_y^T\mathbf{Z}_y}{\sigma_y^2} + \frac{\mathbf{Z}_x^T\mathbf{Z}_x}{\sigma_x^2} + \frac{1}{\sigma_z^2}I_p$$

Let $S = K^{-1}$

The above formula is equal to

$$C \cdot exp\left(-\frac{(\boldsymbol{y}-\mu_y-\mathbf{Z}_y\boldsymbol{\gamma})^T(\boldsymbol{y}-\mu_y-\mathbf{Z}_y\boldsymbol{\gamma})}{2\sigma_y^2}\right).exp\left(-\frac{(\boldsymbol{x}-\mu_x)^T(\boldsymbol{x}-\mu_x)}{2\sigma_x^2}\right)$$

$$\cdot\, exp\left(-\frac{\boldsymbol{\beta}S^{-1}\boldsymbol{\beta}-2Q^TS^{-1}\boldsymbol{\beta}}{2}\right)$$

where

$$Q_{1\times p}^T=\left(\frac{(\boldsymbol{y}-\mu_y-\mathbf{Z}_y\boldsymbol{\gamma})^T\alpha\mathbf{Z}_y}{\sigma_y^2}+\frac{(\boldsymbol{x}-\mu_x)^T\mathbf{Z}_x}{\sigma_x^2}\right)\cdot S$$

Note that the last term is a Gaussian kernel

Thus

$$f(\boldsymbol{x},\boldsymbol{y})=\int f(\boldsymbol{y}|\boldsymbol{\beta})f(\boldsymbol{x}|\boldsymbol{\beta})f(\boldsymbol{\beta})d\boldsymbol{\beta}$$

$$= C\cdot exp\left(-\frac{(\boldsymbol{y}-\mu_y-\mathbf{Z}_y\boldsymbol{\gamma})^T(\boldsymbol{y}-\mu_y-\mathbf{Z}_y\boldsymbol{\gamma})}{2\sigma_y^2}\right).exp\left(-\frac{(\boldsymbol{x}-\mu_x)^T(\boldsymbol{x}-\mu_x)}{2\sigma_x^2}\right)\cdot \exp\left(\frac{Q^TKQ}{2}\right)$$

$$\cdot\,(2\pi)^{\frac{p}{2}}\cdot|S|^{\frac{1}{2}}$$

$$=(2\pi)^{-\frac{n_2}{2}}(\sigma_y^2)^{-\frac{n_2}{2}}(2\pi)^{-\frac{n_1}{2}}(\sigma_x^2)^{-\frac{n_1}{2}}(\sigma_z^2)^{-\frac{p}{2}}|K|^{-\frac{1}{2}}exp\left(-\frac{(\boldsymbol{y}-\mu_y-\mathbf{Z}_y\boldsymbol{\gamma})^T(\boldsymbol{y}-\mu_y-\mathbf{Z}_y\boldsymbol{\gamma})}{2\sigma_y^2}\right).exp\left(-\frac{(\boldsymbol{x}-\mu_x)^T(\boldsymbol{x}-\mu_x)}{2\sigma_x^2}\right.$$

$$\exp\left(\frac{\left(\frac{(\boldsymbol{y}-\mu_y-\mathbf{Z}_y\boldsymbol{\gamma})^T\alpha\mathbf{Z}_y}{\sigma_y^2}+\frac{(\boldsymbol{x}-\mu_x)^T\mathbf{Z}_x}{\sigma_x^2}\right)K^{-1}\left(\frac{(\boldsymbol{y}-\mu_y-\mathbf{Z}_y\boldsymbol{\gamma})^T\alpha\mathbf{Z}_y}{\sigma_y^2}+\frac{(\boldsymbol{x}-\mu_x)^T\mathbf{Z}_x}{\sigma_x^2}\right)^T}{2}\right)$$

Let $\theta=\left(\alpha,\gamma,\sigma_y^2,\sigma_x^2,\sigma_z^2,\mu_x,\mu_y\right)$ indicate all model parameters.

The hypothesis test for $\alpha$ is $H_0:\alpha=0\ v.s.H_1:\alpha\neq 0$

The likelihood ratio test (LRT) is given by

$$\Lambda_\alpha=2\left\{logf\left(x,y\middle|\mathrm{Z}_x,Z_y,\hat{\theta}\right)-logf\left(x,y\middle|\mathrm{Z}_x,Z_y,\hat{\theta}_{\alpha=0}\right)\right\}$$

Where $\widehat{\theta}$ is the parameter estimator, and $\hat{\theta}_{\alpha=0}$ is the estimator under $\alpha=0$. Similarly, the hypothesis test for $\gamma$ is $H_0:\gamma=0\ \ v.s.\ H_1:\gamma\neq 0$

The LRT is given by

$$\Lambda_\gamma=2\left\{logf\left(x,y\middle|\mathrm{Z}_x,Z_y,\hat{\theta}\right)-logf\left(x,y\middle|\mathrm{Z}_x,Z_y,\hat{\theta}_{\gamma=0}\right)\right\}$$

Where $\hat{\theta}_{\gamma=0}$ is the parameter estimator under $\gamma=0$.

### 2. Estimation procedure

We develop an expectation-maximization (EM) algorithm for inference, where we treat the SNP effect sizes $\beta$ as missing data. Traditional EM algorithm converges very slowly while Newton's method may be unstable and sensitive to initial values. Therefore, we use a parameter-expanded version of EM, i.e. PX-EM[1], for estimation. PX-EM improves the convergence rate of traditional EM algorithm while is simple to implement and enjoys the stability of traditional EM. To do so, we consider the parameter expanded version of our model as follows

$$\boldsymbol{x} = \mu_x + \lambda \mathbf{Z}_x \boldsymbol{\beta} + \boldsymbol{\varepsilon}_x \qquad (3)$$

$$\boldsymbol{y} = \mu_y + \mathbf{Z}_y \boldsymbol{\beta} \alpha + \mathbf{Z}_y \boldsymbol{\gamma} + \boldsymbol{\varepsilon}_y \quad (4)$$

Where $\lambda$ is the expanded parameter. Let $\boldsymbol{\theta} = \left(\lambda, \alpha, \gamma, \sigma_y^2, \sigma_x^2, \sigma_z^2, \mu_x, \mu_y\right)$ denote all parameters for parameter expanded model. When $\lambda = 1$, the expanded model is equal to the model for the observated data. The reduction function can be defined as $R\left(\lambda, \alpha, \gamma, \sigma_y^2, \sigma_x^2, \sigma_z^2, \mu_x, \mu_y\right) = \left({}^{\alpha}/_{\lambda}, \gamma, \sigma_y^2, \sigma_x^2, \lambda^2\sigma_z^2, \mu_x, \mu_y\right)$.

From the derivation similar as above, it is easy to obtain that, given $\boldsymbol{x}$, $\boldsymbol{y}$, $\mathbf{Z}_x$, $\mathbf{Z}_y$ and $\boldsymbol{\theta}$, the distribution of the latent variable $\boldsymbol{\beta}$ is a normal distribution $\mathrm{N}(\beta|\mu_\beta, \Sigma_\beta)$, where

$$\Sigma_\beta^{-1} = \left(\frac{\alpha^2 \boldsymbol{Z}_y^T \boldsymbol{Z}_y}{\sigma_y^2} + \frac{\lambda^2 \boldsymbol{Z}_x^T \boldsymbol{Z}_x}{\sigma_x^2} + \frac{1}{\sigma_z^2}\boldsymbol{I}_p\right)$$

$$\mu_\beta = \left(\frac{\alpha^2 \boldsymbol{Z}_y^T \boldsymbol{Z}_y}{\sigma_y^2} + \frac{\lambda^2 \boldsymbol{Z}_x^T \boldsymbol{Z}_x}{\sigma_x^2} + \frac{1}{\sigma_z^2}\boldsymbol{I}_p\right)^{-1} \left(\frac{\lambda}{\sigma_x^2}\boldsymbol{Z}_x^T(\boldsymbol{x} - \mu_x) + \frac{\alpha}{\sigma_y^2}\boldsymbol{Z}_y^T\left(\boldsymbol{y} - \mu_y - \mathbf{Z}_y\boldsymbol{\gamma}\right)\right)$$

The complete likelihood for the parameter expanded model can be calculated as

$$f(\boldsymbol{y}|\boldsymbol{\beta})f(\boldsymbol{x}|\boldsymbol{\beta})f(\boldsymbol{\beta}) = (2\pi)^{-\frac{n_2}{2}}(\sigma_y^2)^{-\frac{n_2}{2}}\exp\left(-\frac{(\boldsymbol{y}-\mu_y-\alpha\mathbf{Z}_y\boldsymbol{\beta}-\mathbf{Z}_y\boldsymbol{\gamma})^T(\boldsymbol{y}-\mu_y-\alpha\mathbf{Z}_y\boldsymbol{\beta}-\mathbf{Z}_y\boldsymbol{\gamma})}{2\sigma_y^2}\right)\cdot$$

$$(2\pi)^{-\frac{n_1}{2}}(\sigma_x^2)^{-\frac{n_1}{2}}\exp\left(-\frac{\left(\boldsymbol{x} - \mu_x - \lambda\mathbf{Z}_x\boldsymbol{\beta}\right)^T\left(\boldsymbol{x} - \mu_x - \lambda\mathbf{Z}_x\boldsymbol{\beta}\right)}{2\sigma_x^2}\right)(2\pi)^{-\frac{p}{2}}(\sigma_z^2)^{-\frac{p}{2}}exp\left(\frac{\boldsymbol{\beta}^T\boldsymbol{\beta}}{2\sigma_z^2}\right)$$

**In the E-step**, we derive $\boldsymbol{Q}$ function by taking expectation of the complete-data log-likelihood with respect to the distribution $\mathrm{N}(\beta|\mu_\beta, \Sigma_\beta)$. Remember that $E(\beta^T A\beta) = \mu_\beta^T A\mu_\beta + Tr(A\Sigma_\beta)$ for any symmetric matrix *A*, where $\mathrm{Tr}(M)$ denotes the trace of matrix *M*. We can get

$$E\left((\boldsymbol{x} - \mu_x - \lambda\mathbf{Z}_x\boldsymbol{\beta})^T(\boldsymbol{x} - \mu_x - \lambda\mathbf{Z}_x\boldsymbol{\beta})\right)$$

$$= \left(\boldsymbol{x} - \mu_x - \lambda\mathbf{Z}_x\mu_\beta\right)^T\left(\boldsymbol{x} - \mu_x - \lambda\mathbf{Z}_x\mu_\beta\right) + \lambda^2 Tr\left(\boldsymbol{Z}_x^T\boldsymbol{Z}_x\Sigma_\beta\right)$$

$$E\left(\left(\boldsymbol{y} - \mu_y - \mathbf{Z}_y\boldsymbol{\beta}\alpha - \mathbf{Z}_y\boldsymbol{\gamma}\right)^T\left(\boldsymbol{y} - \mu_y - \mathbf{Z}_y\boldsymbol{\beta}\alpha - \mathbf{Z}_y\boldsymbol{\gamma}\right)\right)$$

$$= \left(\boldsymbol{y} - \mu_y - \mathbf{Z}_y\boldsymbol{\gamma} - \alpha\mathbf{Z}_y\mu_\beta\right)^T\left(\boldsymbol{y} - \mu_y - \mathbf{Z}_y\boldsymbol{\gamma} - \alpha\mathbf{Z}_y\mu_\beta\right) + \alpha^2 Tr\left(\boldsymbol{Z}_y^T\boldsymbol{Z}_y\Sigma_\beta\right)$$

$$E(\beta^T\beta) = \mu_\beta^T\mu_\beta + Tr(\Sigma_\beta)$$

Given the current value $\boldsymbol{\theta}_{old}$ and the observed data, the $\boldsymbol{Q}$ function is

$$\boldsymbol{Q}(\boldsymbol{\theta}|\boldsymbol{\theta}_{old}) = -\frac{n_1}{2}log\,\sigma_{\mathrm{x}}^2 - \frac{n_2}{2}log\sigma_{\mathrm{y}}^2 - \frac{p}{2}log\sigma_{\mathrm{z}}^2 - \frac{1}{2\sigma_{\mathrm{z}}^2}\mu_\beta^T\mu_\beta$$

$$-\frac{1}{2\sigma_{\mathrm{x}}^2}\left[\left(\boldsymbol{x} - \mu_x - \lambda\mathbf{Z}_{\mathrm{x}}\mu_\beta\right)^T\left(\boldsymbol{x} - \mu_x - \lambda\mathbf{Z}_{\mathrm{x}}\mu_\beta\right)\right]$$

$$- -\frac{1}{2\sigma_{\mathrm{y}}^2}\left[\left(\boldsymbol{y} - \mu_{\mathrm{y}} - \mathbf{Z}_{\mathrm{y}}\boldsymbol{\gamma} - \alpha\mathbf{Z}_{\mathrm{y}}\mu_\beta\right)^T\left(\boldsymbol{y} - \mu_{\mathrm{y}} - \mathbf{Z}_{\mathrm{y}}\boldsymbol{\gamma} - \alpha\mathbf{Z}_{\mathrm{y}}\mu_\beta\right)\right]$$

$$- Tr\left[\left(\frac{\lambda^2}{2\sigma_{\mathrm{x}}^2}\boldsymbol{Z}_x^T\boldsymbol{Z}_x + \frac{\alpha^2}{2\sigma_{\mathrm{y}}^2}\boldsymbol{Z}_y^T\boldsymbol{Z}_y + \frac{1}{2\sigma_{\mathrm{z}}^2}\boldsymbol{I}_p\right)\Sigma_\beta\right]$$

**In the M-step**, by setting the derivative of $\boldsymbol{Q}$ function to zero, we obtain the new updates for all parameters. Where

$$\lambda = \frac{(\boldsymbol{x} - \mu_x)^T\mathbf{Z}_{\mathrm{x}}\mu_\beta}{\mu_\beta^T\boldsymbol{Z}_x^T\boldsymbol{Z}_x\mu_\beta + Tr\left(\boldsymbol{Z}_x^T\boldsymbol{Z}_x\Sigma_\beta\right)}$$

$$\gamma = \frac{\mathbf{1}_p^T\boldsymbol{Z}_y^T\left(\boldsymbol{y} - \mu_{\mathrm{y}} - \alpha\mathbf{Z}_{\mathrm{y}}\mu_\beta\right)}{\mathbf{1}_p^T\boldsymbol{Z}_y^T\boldsymbol{Z}_y\mathbf{1}_p^T}$$

$$\alpha = \frac{\left(\boldsymbol{y} - \mu_{\mathrm{y}} - \mathbf{Z}_{\mathrm{y}}\boldsymbol{\gamma}\right)^T\boldsymbol{Z}_y\mu_\beta}{\mu_\beta^T\boldsymbol{Z}_y^T\boldsymbol{Z}_y\mu_\beta + Tr\left(\boldsymbol{Z}_y^T\boldsymbol{Z}_y\Sigma_\beta\right)}$$

$$\sigma_x^2 = \frac{1}{n_1}\left[\left(\boldsymbol{x} - \mu_x - \lambda\mathbf{Z}_{\mathrm{x}}\mu_\beta\right)^T\left(\boldsymbol{x} - \mu_x - \lambda\mathbf{Z}_{\mathrm{x}}\mu_\beta\right) + Tr\left(\lambda^2\boldsymbol{Z}_x^T\boldsymbol{Z}_x\Sigma_\beta\right)\right]$$

$$\sigma_y^2 = \frac{1}{n_2}\left[\left(\boldsymbol{y} - \mu_{\mathrm{y}} - \mathbf{Z}_{\mathrm{y}}\boldsymbol{\gamma} - \alpha\mathbf{Z}_{\mathrm{y}}\mu_\beta\right)^T\left(\boldsymbol{y} - \mu_{\mathrm{y}} - \mathbf{Z}_{\mathrm{y}}\boldsymbol{\gamma} - \alpha\mathbf{Z}_{\mathrm{y}}\mu_\beta\right) + Tr\left(\alpha^2\boldsymbol{Z}_y^T\boldsymbol{Z}_y\Sigma_\beta\right)\right]$$

$$\sigma_{\mathrm{z}}^2 = \frac{1}{p}\left[\mu_\beta^T\mu_\beta + Tr\left(\Sigma_\beta\right)\right]$$

$$\mu_x = \frac{1}{n_1}\left(\boldsymbol{x} - \lambda\mathbf{Z}_{\mathrm{x}}\mu_\beta\right)^T\mathbf{1}_{n_1}$$

$$\mu_y = \frac{1}{n_2}\left(\boldsymbol{y} - \mathbf{Z}_{\mathrm{y}}\boldsymbol{\gamma} - \alpha\mathbf{Z}_{\mathrm{y}}\mu_\beta\right)^T\mathbf{1}_{n_2}$$

Where $\mathbf{1}_d$ denote the length *d* vector with all elements to be 1.

**In the Reduction step**, we re-set the estimation of parameters using the reduction function $R$, and re-assigned $\lambda = 1$.

Finally, for TWAS applications, we note that some genes have close to zero heritability. For genes whose expression levels are not affected by cis-SNP genotypes, not much information is available for the estimation of $\boldsymbol{\beta}$, which subsequently leads to an extremely large standard error of $\hat{\alpha}$. Therefore, PMR-Egger can return the p value equal or close to 1 for these genes.

### 3. Causal effect identification

The causal interpretation of the parameter $\alpha$ and its identification can be derived under the framework of decision-theoretic causal inference[2-5]. We define the causal effect of gene expression

$X$ on the phenotype Y as the difference between the expected values of Y under an intervention that imposes on $X$ a reference value $x_0$ and another intervention that imposes another value $x$. Let the symbol $F_X$ label the regime under which the value of $X$ is generated, with $F_X = x$ indicating that $X$ is fixed to value $x$ by an intervention, and $F_X = \emptyset$ denoting the observational regime under which the data have actually been generated. Then the average causal effect (ACE) of $X$ on the continuous phenotype Y is defined by

$$\mathrm{ACE} = \mathrm{E}(Y|F_X = x) - \mathrm{E}(Y|F_X = x_0)$$

Let the notation $A \perp\!\!\!\perp \mathrm{B}|\, C$ indicates that $A$ is independent of $B$ given $C$.

Our proposed MR model based on the observational data obtained under $F_X = \emptyset$ has been presented in Figure S1. Note that we directly model the horizontal pleiotropic effects through the $Z \rightarrow \mathrm{y}$ arrow. Therefore, our model does not require the Exclusion Restriction condition of traditional MR. However, the other two assumptions in tradtional MR must be satisfied; that is, $Z$ is associated with $X$, and $U \perp\!\!\!\perp Z$ **(1)**.

One must note that requirements relating only to the observational regime can never be sufficient to estimate the causal effect of $X$ on y, which is defined in terms of interventional regimes. Instead, we need to make additional assumptions that relate the observational regime $F_X = \emptyset$ to the interventional regimes $F_X = x$. Under the assumption that the unobserved $U$ is a sufficient covariate for the effect of $X$ on y, we can do this by elaborating Figure S1 to explicitly include the nonstochastic regime indicator $F_X$ for $X$, as in the following Figure. For $F_X = \emptyset$, this recovers the assumptions embedded in Figure S1, but in addition it relates the observational structure to what would happen under an intervention to set $X$.

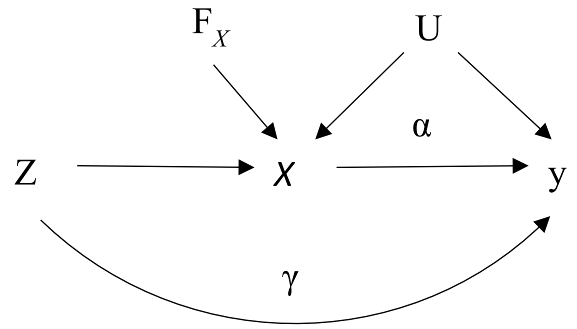


Figure. The causal diagram of PMR-Egger with the nonstochastic regime indicator $F_X$

It is illustrated that an intervention on $X$ will not affect $Z$ or U, that is $F_X \perp\!\!\!\perp (\mathrm{U}, Z)$ **(2)**. And conditional on $Z$ and U, the distribution of y given $X$ does not depend on whether the value of $X$ has been generated by passive observation or intervention; that is $\mathrm{y} \perp\!\!\!\perp F_X|(X, Z, \mathrm{U})$ **()).** Furthermore, the formula **(1)** can be extended to $\mathrm{U} \perp\!\!\!\perp Z|\, F_X$ **(4)**.

We can describe the dependence of y on $(X, \mathrm{U})$ (the same in all regimes by (3)) by a linear model: $\mathrm{E}(\mathrm{y}|X, Z, \mathrm{U}) = \mathrm{W} + \alpha X + \gamma Z$ **(5)**, where W is some function of U.

Because (5) holds in the interventional regime $F_X = x$, we deduce

$$\mathrm{E}(\mathrm{y}|F_X = x) = \mathrm{W}_0 + \alpha x + \mathrm{Z}_0$$

where $\mathrm{W}_0 := \mathrm{E}(\mathrm{W}|F_X = x)$ and $\mathrm{Z}_0 := \gamma \mathrm{E}(Z|F_X = x)$ is a constant independent of $x$ following (2). Thus α can be interpreted causally, as it describs how the mean of y responds to manipulation of *X*. Next we show how to estimate α.

Again by (3), the formula (5) is also $\mathrm{E}(\mathrm{y}|X, Z, \mathrm{U}, \mathrm{F}_X = \emptyset)$. Then

$$\mathrm{E}(\mathrm{y}|Z, \mathrm{F}_X = \emptyset) = \mathrm{E}(\mathrm{W}|Z, \mathrm{F}_\mathrm{X} = \emptyset) + \alpha\mathrm{E}(X|Z, \mathrm{F}_\mathrm{X} = \emptyset) + \gamma Z$$

By (4), the first term on the right side is constant, thus

$\mathrm{E}(\mathrm{y}|Z, \mathrm{F}_X = \emptyset) = \mathrm{constant} + \alpha\mathrm{E}(X|Z, \mathrm{F}_\mathrm{X} = \emptyset) + \gamma Z$ **(6)**

Equation (6) relates two functions of $Z$, each of which can be identified from observational data. Consequently, we can estimate the causal parameter $\alpha$ from such data.

### 4. PMR-Egger model for summary statistics

We denote the LD structure of the cis-SNPs for one specific gene as $\Sigma_1$ in gene expression data, and $\Sigma_2$ in GWAS data; both are $\mathrm{p} \times \mathrm{p}$ symmetric positive definite matrices. The marginal estimates of $\mathbf{Z}_\mathrm{x}$ on $\boldsymbol{x}$ are $\hat{\beta}_x^*$ (eQTL effects), the marginal estimates of $\mathbf{Z}_y$ on $\boldsymbol{y}$ are $\hat{\beta}_y^*$ (GWAS effects). While the corresponding conditional estimates are $\hat{\beta}_x = \Sigma_1^{-1}\hat{\beta}_x^*$, $\hat{\beta}_y = \Sigma_2^{-1}\hat{\beta}_y^*$.

The corresponding model for summary statistics are

$$\begin{cases} \hat{\beta}_x = \boldsymbol{\beta} + E_1 & (3) \\ \hat{\beta}_y = \boldsymbol{\gamma} + \alpha\boldsymbol{\beta} + E & (4) \end{cases}$$

where $\boldsymbol{\beta} \sim N(0, \sigma_\mathrm{z}^2 I_p)$,$E_1 \sim N(0, {\Sigma_1}^{-1}\sigma_\mathrm{x}^2)$, $E \sim N\left(0, {\Sigma_2}^{-1}\sigma_\mathrm{y}^2\right)$, $\boldsymbol{\gamma}$ is the pleiotropy effect vector with each element equal to common parameter $\gamma$.

Given $\boldsymbol{\beta}, \hat{\beta}_x$ and $\hat{\beta}_y$ are independent.

The observed likelihood function is

$$f(\hat{\beta}_x, \hat{\beta}_y) = \int f(\hat{\beta}_x, \hat{\beta}_y, \boldsymbol{\beta})d\boldsymbol{\beta} = \int f(\hat{\beta}_x, \hat{\beta}_y|\boldsymbol{\beta})f(\boldsymbol{\beta})d\boldsymbol{\beta} = \int f(\hat{\beta}_y|\boldsymbol{\beta})f(\hat{\beta}_x|\boldsymbol{\beta})f(\boldsymbol{\beta})d\boldsymbol{\beta}$$

$$f(\hat{\beta}_y|\boldsymbol{\beta})f(\hat{\beta}_x|\boldsymbol{\beta})f(\boldsymbol{\beta}) = (2\pi)^{-\frac{p}{2}}|{\Sigma_1}^{-1}\sigma_\mathrm{x}^2|^{-\frac{1}{2}} \exp\left(-\frac{(\hat{\beta}_x - \boldsymbol{\beta})^T \Sigma_1 (\hat{\beta}_x - \boldsymbol{\beta})}{2\sigma_\mathrm{x}^2}\right)(2\pi)^{-\frac{p}{2}}|{\Sigma_2}^{-1}\sigma_\mathrm{y}^2|^{-\frac{1}{2}}$$

$$\exp\left(-\frac{(\hat{\beta}_y - \boldsymbol{\gamma} - \alpha\boldsymbol{\beta})^T \Sigma_2 (\hat{\beta}_y - \mathbf{u} - \alpha\boldsymbol{\beta})}{2\sigma_\mathrm{y}^2}\right)(2\pi)^{-\frac{p}{2}}(\sigma_\mathrm{z}^2)^{-\frac{p}{2}} exp\left(-\frac{\boldsymbol{\beta}^T\boldsymbol{\beta}}{2\sigma_\mathrm{z}^2}\right)$$

$$= C \cdot exp\left(-\frac{(\hat{\beta}_y - \boldsymbol{\gamma})^T \Sigma_2 (\beta_y - \boldsymbol{\gamma})}{2\sigma_\mathrm{y}^2}\right) \cdot exp\left(-\frac{{\hat{\beta}_x}^T \Sigma_1 \hat{\beta}_x}{2\sigma_\mathrm{x}^2}\right)$$

$$\cdot\, exp\left(-\frac{\boldsymbol{\beta}^T K \boldsymbol{\beta} - 2\left(\frac{(\hat{\beta}_y - \boldsymbol{\gamma})^T \Sigma_2 \alpha}{\sigma_\mathrm{y}^2} + \frac{{\hat{\beta}_x}^T \Sigma_1}{\sigma_x^2}\right)\boldsymbol{\beta}}{2}\right)$$

where $C = (2\pi)^{-\frac{p}{2}}|{\Sigma_1}^{-1}\sigma_\mathrm{x}^2|^{-\frac{1}{2}}(2\pi)^{-\frac{p}{2}}|{\Sigma_2}^{-1}\sigma_\mathrm{y}^2|^{-\frac{1}{2}}(2\pi)^{-\frac{p}{2}}(\sigma_\mathrm{z}^2)^{-\frac{p}{2}}$,$\mathrm{K} = \frac{1}{\sigma_\mathrm{z}^2} I_p + \frac{\alpha^2}{\sigma_\mathrm{y}^2}\Sigma_2 + \frac{\Sigma_1}{\sigma_\mathrm{x}^2}$

Let $S = K^{-1}$

The above formula is equal to

$$C \cdot exp\left(-\frac{(\hat{\beta}_y - \boldsymbol{\gamma})^T \Sigma_2 (\beta_y - \boldsymbol{\gamma})}{2\sigma_\mathrm{y}^2}\right) \cdot exp\left(-\frac{{\hat{\beta}_x}^T \Sigma_1 \hat{\beta}_x}{2\sigma_\mathrm{x}^2}\right) exp\left(-\frac{\boldsymbol{\beta}^T S^{-1} \boldsymbol{\beta} - 2Q^T S^{-1} \boldsymbol{\beta}}{2}\right)$$

where $Q_{1\times p}^T = \left(\frac{(\hat{\beta}_y - \boldsymbol{\gamma})^T \Sigma_2 \alpha}{\sigma_\mathrm{y}^2} + \frac{{\hat{\beta}_x}^T \Sigma_1}{\sigma_\mathrm{x}^2}\right) \cdot \mathrm{S}$. Thus

$$f(\hat{\beta}_x, \hat{\beta}_y) = C \cdot exp\left(-\frac{(\hat{\beta}_y - \boldsymbol{\gamma})^T \Sigma_2 (\beta_y - \boldsymbol{\gamma})}{2\sigma_\mathrm{y}^2}\right) \cdot exp\left(-\frac{{\hat{\beta}_x}^T \Sigma_1 \hat{\beta}_x}{2\sigma_\mathrm{x}^2}\right) \cdot exp\left(\frac{Q^T K Q}{2}\right) \cdot (2\pi)^{\frac{p}{2}}|S|^{\frac{1}{2}}$$

$$= (2\pi)^{-\frac{p}{2}}|\Sigma_1{}^{-1}\sigma_x^2|^{-\frac{1}{2}}(2\pi)^{-\frac{p}{2}}|\Sigma_2{}^{-1}\sigma_y^2|^{-\frac{1}{2}}(\sigma_z^2)^{-\frac{p}{2}}|K|^{-\frac{1}{2}} exp\left(-\frac{(\hat{\beta}_y-\boldsymbol{\gamma})^T\Sigma_2(\beta_y-\boldsymbol{\gamma})}{2\sigma_y^2}\right)$$

$$\cdot\, exp\left(-\frac{\hat{\beta}_x{}^T\Sigma_1\hat{\beta}_x}{2\sigma_x^2}\right)\exp\left(\frac{\left(\frac{(\hat{\beta}_y-\boldsymbol{\gamma})^T\Sigma_2\alpha}{\sigma_y^2}+\frac{\hat{\beta}_x{}^T\Sigma_1}{\sigma_x^2}\right)K^{-1}\left(\frac{(\hat{\beta}_y-\boldsymbol{\gamma})^T\Sigma_2\alpha}{\sigma_y^2}+\frac{\hat{\beta}_x{}^T\Sigma_1}{\sigma_x^2}\right)^T}{2}\right)$$

Let $\theta = (\alpha, \gamma, \sigma_y^2, \sigma_x^2, \sigma_z^2)$ indicate all model parameters.

The hypothesis test for $\alpha$ is $H_0\colon \alpha = 0$ $v.s. H_1\colon \alpha \neq 0$.

The likelihood ratio test (LRT) is given by

$$\Lambda_\alpha = 2\left\{logf(\hat{\beta}_x, \hat{\beta}_y|\hat{\theta}) - logf(\hat{\beta}_x, \hat{\beta}_y|\hat{\theta}_{\alpha=0})\right\}$$

where $\widehat{\theta}$ is the parameter estimator, and $\hat{\theta}_{\alpha=0}$ is the estimator under $\alpha = 0$. Similarly, the hypothesis test for $\gamma$ is $H_0\colon \gamma = 0$ $v.s.$ $H_1\colon \gamma \neq 0$.

The LRT is given by

$$\Lambda_\gamma = 2\left\{logf(\hat{\beta}_x, \hat{\beta}_y|\hat{\theta}) - logf(\hat{\beta}_x, \hat{\beta}_y|\hat{\theta}_{\gamma=0})\right\}$$

where $\hat{\theta}_{\gamma=0}$ is the parameter estimator under $\gamma = 0$.

### 5. PX-EM algorithm for summary statistics

The parameter expanded version of our model is

$$\begin{cases} \hat{\beta}_x = \lambda\boldsymbol{\beta} + E_1 & (3) \\ \hat{\beta}_y = \boldsymbol{\gamma} + \alpha\boldsymbol{\beta} + E & (4) \end{cases}$$

where $\lambda$ is the expanded scalar parameter. Let $\boldsymbol{\Omega} = (\lambda, \alpha, \gamma, \sigma_y^2, \sigma_x^2, \sigma_z^2)$ denote all parameters for parameter expanded model. When $\lambda = 1$, the expanded model is equal to the model for the observated data. The reduction function can be defined as $R(\lambda, \alpha, \gamma, \sigma_y^2, \sigma_x^2, \sigma_z^2) = ({}^{\alpha}/_{\lambda}, \gamma, \sigma_y^2, \sigma_x^2, \lambda^2\sigma_z^2)$.

Treating $\boldsymbol{\beta}$ as the missing variable, the complete data likelihood is

$$(2\pi)^{-\frac{p}{2}}|\Sigma_1{}^{-1}\sigma_x^2|^{-\frac{1}{2}}\exp\left(-\frac{(\hat{\beta}_x-\lambda\boldsymbol{\beta})^T\Sigma_1(\hat{\beta}_x-\lambda\boldsymbol{\beta})}{2\sigma_x^2}\right)(2\pi)^{-\frac{p}{2}}|\Sigma_2{}^{-1}\sigma_y^2|^{-\frac{1}{2}}$$

$$\exp\left(-\frac{(\hat{\beta}_y-\boldsymbol{\gamma}-\alpha\boldsymbol{\beta})^T\Sigma_2(\hat{\beta}_y-\boldsymbol{\gamma}-\alpha\boldsymbol{\beta})}{2\sigma_y^2}\right)(2\pi)^{-\frac{p}{2}}(\sigma_z^2)^{-\frac{p}{2}}exp\left(-\frac{\boldsymbol{\beta}^T\boldsymbol{\beta}}{2\sigma_z^2}\right)$$

Given the data $\hat{\beta}_x$ and $\hat{\beta}_y$, and parameters $\boldsymbol{\Omega}$, it is easy to obtain the distribution of $\boldsymbol{\beta}$ is Gaussian with the mean

$$\mu_\beta = \left(\frac{1}{\sigma_z^2}I_p + \frac{\alpha^2}{\sigma_y^2}\Sigma_2 + \frac{\lambda^2\Sigma_1}{\sigma_x^2}\right)^{-1}\cdot\left(\frac{(\hat{\beta}_y-\boldsymbol{\gamma})^T\Sigma_2\alpha}{\sigma_y^2} + \frac{\hat{\beta}_x{}^T\lambda\Sigma_1}{\sigma_x^2}\right)^T$$

the covariance matrix $\Sigma_\beta = \left(\frac{1}{\sigma_z^2}I_p + \frac{\alpha^2}{\sigma_y^2}\Sigma_2 + \frac{\lambda^2\Sigma_1}{\sigma_x^2}\right)^{-1}$

Using formula $E(\beta^T A\beta) = \mu_\beta^T A\mu_\beta + Tr(A\Sigma_\beta)$ for any symmetrix *A*.

The *E*-step can generate the following Q function

$$Q(\theta|\hat{\beta}_x, \hat{\beta}_y) = -\frac{p}{2}log2\pi + \frac{1}{2}log|\Sigma_1| - \frac{p}{2}log\,\sigma_x^2 - \frac{p}{2}log2\pi + \frac{1}{2}log|\Sigma_2| - \frac{p}{2}log\,\sigma_y^2$$

$$-\frac{p}{2}log2\pi - \frac{p}{2}log\,\sigma_z^2 - \frac{1}{2\sigma_z^2}\left[\mu_\beta^T\mu_\beta + tr(\Sigma_\beta)\right]$$

$$-\frac{1}{2\sigma_x^2}\left[\hat{\beta}_x{}^T\Sigma_1\hat{\beta}_x - 2\hat{\beta}_x{}^T\lambda\Sigma_1\mu_\beta + \lambda^2\left(\mu_\beta^T\Sigma_1\mu_\beta + tr(\Sigma_1\Sigma_\beta)\right)\right]$$

$$-\frac{1}{2\sigma_y^2}\Big\{(\hat{\beta}_y - \boldsymbol{\gamma})^T\Sigma_2(\hat{\beta}_y - \boldsymbol{\gamma}) - 2(\hat{\beta}_y - \boldsymbol{\gamma})^T\Sigma_2\alpha\mu_\beta$$

$$+\alpha^2\left[\mu_\beta^T\Sigma_2\mu_\beta + tr(\Sigma_2\Sigma_\beta)\right]\Big\}$$

In the *M*-step, we set the derivative of the Q function to zero and obtain the following updated equations for all parameters.

$$\lambda = \frac{\hat{\beta}_x\Sigma_1\mu_\beta}{\mu_\beta^T\Sigma_1\mu_\beta + tr(\Sigma_1\Sigma_\beta)}$$

$$\alpha = \frac{(\hat{\beta}_y - \boldsymbol{\gamma})^T\Sigma_2\mu_\beta}{\mu_\beta^T\Sigma_2\mu_\beta + tr(\Sigma_2\Sigma_\beta)}$$

$$\sigma_x^2 = \frac{1}{p}\left[\hat{\beta}_x{}^T\Sigma_1\hat{\beta}_x - 2\hat{\beta}_x{}^T\lambda\Sigma_1\mu_\beta + \lambda^2\left(\mu_\beta^T\Sigma_1\mu_\beta + tr(\Sigma_1\Sigma_\beta)\right)\right]$$

$$\sigma_y^2 = \frac{1}{p}\left\{\left(\hat{\beta}_y - \boldsymbol{\gamma}\right)^T\Sigma_2\left(\hat{\beta}_y - \boldsymbol{\gamma}\right) - 2\left(\hat{\beta}_y - \boldsymbol{\gamma}\right)^T\Sigma_2\alpha\mu_\beta + \alpha^2[\mu_\beta^T\Sigma_2\mu_\beta + tr(\Sigma_2\Sigma_\beta)]\right\}$$

$$\sigma_z^2 = \frac{1}{p}\left[\mu_\beta^T\mu_\beta + Tr\left(\Sigma_\beta\right)\right]$$

$$\gamma = \frac{\mathbf{1}_p^T\Sigma_2\left(\hat{\beta}_y - \alpha\mu_\beta\right)}{\mathbf{1}_p^T\Sigma_2\mathbf{1}_p^T}$$

where $\mathbf{1}_p$is a length p vector with elements equal to 1.

**In the Reduction step**, we re-set the estimation of parameters using the reduction function $R$, and re-assigned $\lambda = 1$.

### 6. Sparse vs polygenic modeling assumptions in previous TWAS methods

Currently, almost all previous TWAS methods (TWAS[6], PrediXcan[7], CoMM[8], DPR[9],TIGAR[10] etc.) make a polygenic modeling assumption and assume that cis-SNPs have non-zero polygenic effects on gene expression. Specifically, TWAS makes the BSLMM[11] polygenic modeling assumption: all cis-SNPs have non-zero effects and their effect sizes follow a mixture of two normal distributions. PrediXcan[7] makes the ElasticNet modeling assumption: all cis-SNPs have non-zero effects *a priori* and their effect sizes follows a mixture of Laplace (L1) and normal (L2) distributions. Both TIGAR and DPR makes the Bayesian non-parametric polygenic modeling assumption: all cis-SNPs have non-zero effects and their effect sizes follow a mixture of many normal distributions. CoMM makes the standard polygenic modeling assumption: all cis-SNPs have non-zero effects and their effect sizes follow a normal distribution. Perhaps the only method previously used in TWAS setting that makes a sparse modeling assumption is SMR[12]. Certainly, while the modeling assumption underlying PrediXcan is polygenic, it does use an optimization algorithm that obtains the posterior mode estimates (which is sparse) instead of the posterior mean estimates (which is polygenic). However, it is important to distinguish the modeling assumption from the inference algorithm. Overall, the polygenic modeling assumption made in most existing TWAS methods are consistent with the previously known fact that polygenic models often outperform sparse models in gene expression prediction[4,7] and are also consistent with our current results showing that polygenic models also outperform sparse models for TWAS applications.

### 7. Number of discoveries and the genomic control factor in real data applications

The number of genes identified above a genome-wide threshold is commonly used in the literature as a measure of statistical power in real data applications. The genomic control factor $\lambda$ is commonly used in the literature to measure type I error control for real data applications, as $\lambda$ captures approximately the type I error control at the level of 0.05. Power and $\lambda$ are two different statistical terms that are not necessarily correlated with each other. A similar $\lambda$ among different methods would suggest similar type I error control at the nominal level of 0.05, while a higher number of genes identified by one method over another would suggests its higher power. Certainly, for a data with a relatively large sample size (e.g. UK Biobank) and a trait with a highly polygenic genetic architecture, then $\lambda$ may also be influenced by power of the method in addition to its type I error control. In addition, the number of significant genes may not be a perfect measure of power in certain cases and can be influenced by $\lambda$, as a method that fails to control for type I error could yield inflated p-values, leading to a high number of false discoveries.

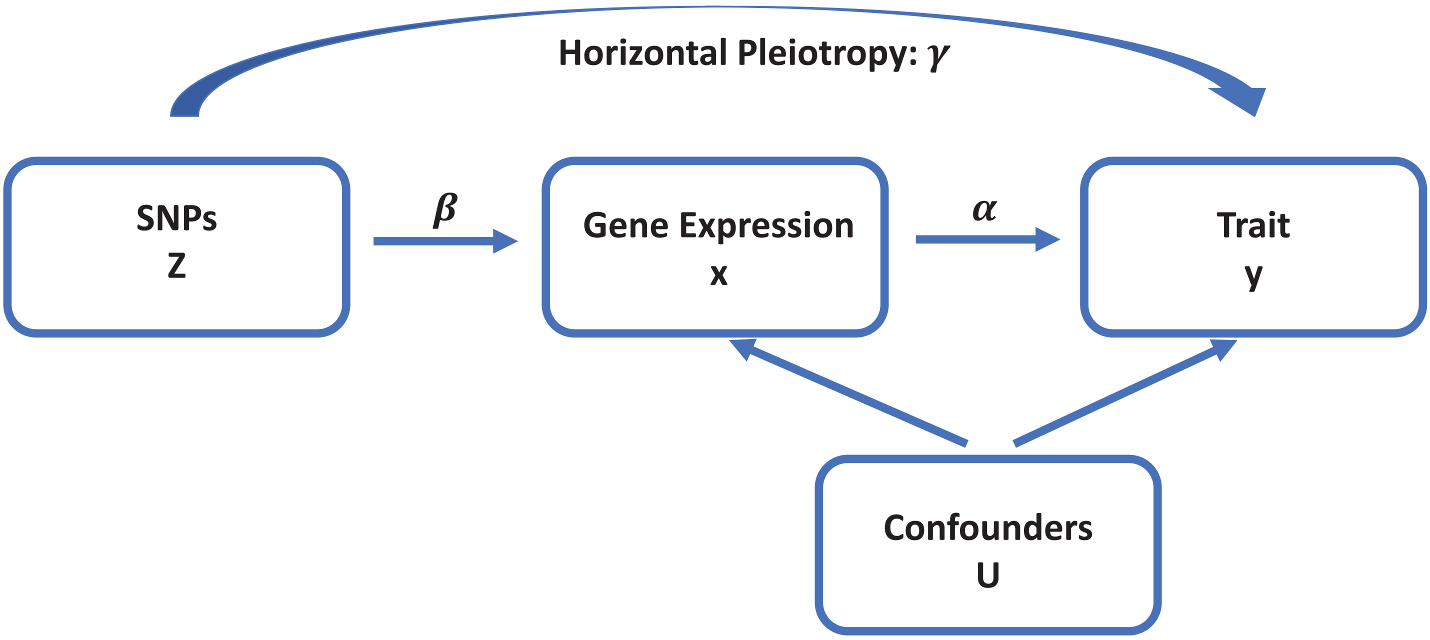


**Fig.1** An illustrative diagram for Mendelian randomization analysis. Mendelian randomization analysis in the TWAS setting attempts to estimate the causal effect of gene expression (x) on the trait of interest (y) in the presence of confounding factors (U) by using cis-SNPs (Z) as instrumental variables. An important requirement of Mendelian randomization analysis is to model and control for horizontal pleiotropic effects.

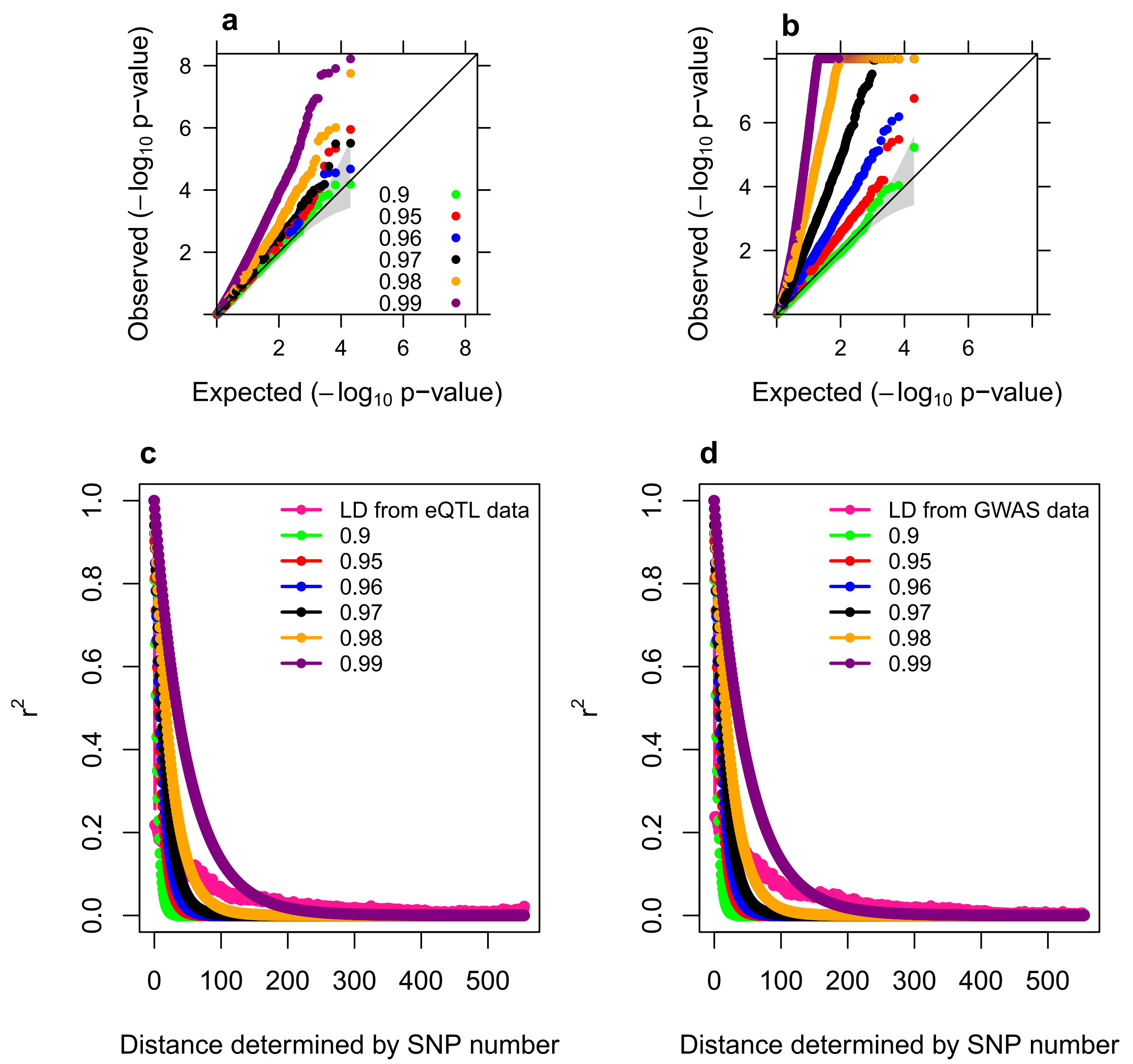


**Fig. 2** Quantile-quantile plot of -log10 p-values from LDA MR-Egger under null simulations. LDA MR-Egger tests for either causal effect (a) or horizontal pleiotropic effect (b). We follow the same simulation design in the original LDA MR-Egger paper to perform simulations. The simulation design assumes that the SNP covariance matrix is an AR(1) covariance structure, where we set the autocorrelation parameter to be either 0.9 (green), 0.95 (red), 0.96 (blue), 0.97 (black), 0.98 (orange), and 0.99 (purple). The inflation of LDA MR-Egger becomes apparent when the autocorrelation becomes greater than 0.9, and such inflation increases with increasing autocorrelation values. The LD decay pattern under these covariance structures are plotted together with the realistic LD pattern of the BACE1 gene (pink). The LD pattern of the BACE1 gene is estimated either in the eQTL data (c) or in the GWAS data (d).

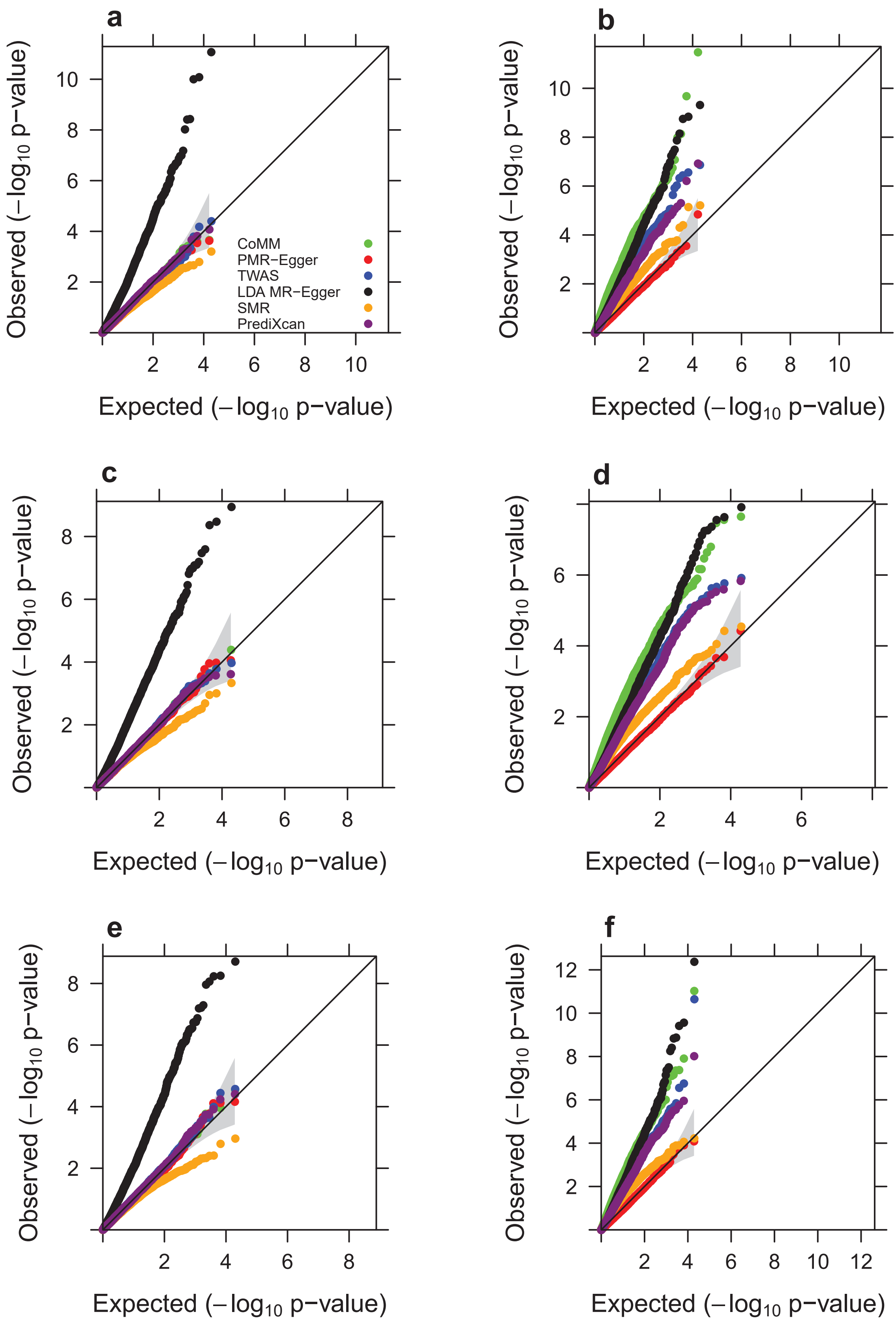


**Fig. 3** Quantile-quantile plot of -log10 p-values from different methods for testing the causal effect under the null simulations, in various sparse settings where only a small proportion of SNPs are associated with the gene expression level. Compared methods include CoMM (green), PMR-Egger (red), TWAS (blue), LDA MR-Egger (black), SMR (orange), and PrediXcan (purple). Simulations are performed either in the absence ($\gamma$=0; a, c, e) or in the presence of horizontal pleiotropic effect ($\gamma$=0.001; b, d, f). Either one SNP (a, b), 1% of SNPs (c, d), or 10% SNPs (e, f) have non-zero effects on gene expression. Only p-values from PMR-Egger adhere to the expected diagonal line across a range of settings.

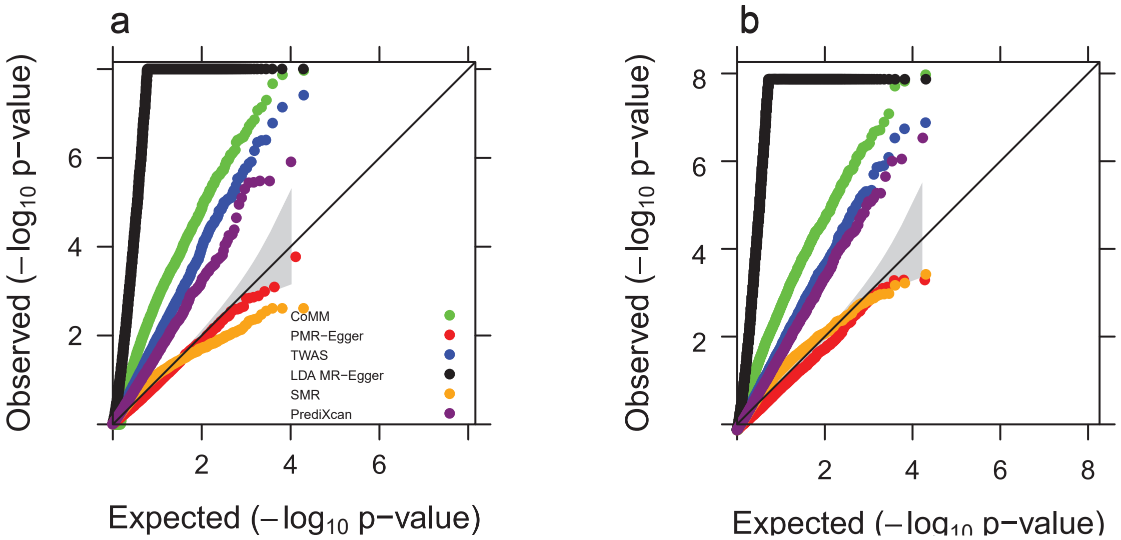


**Fig. 4** Quantile-quantile plot of -log10 p-values from different methods for testing the causal effect under the null, across different gene expression heritability values. Compared methods include CoMM (green), PMR-Egger (red), TWAS (blue), LDA MR-Egger (black), SMR (orange), and PrediXcan (purple). Simulations are performed or in the presence of horizontal pleiotropic effect ($\gamma$=0.001) with gene expression heritability being either (a) $PVE_{zx}$=1% or (b) $PVE_{zx}$=5%. Only p-values from PMR-Egger adhere to the expected diagonal line across a range of settings.

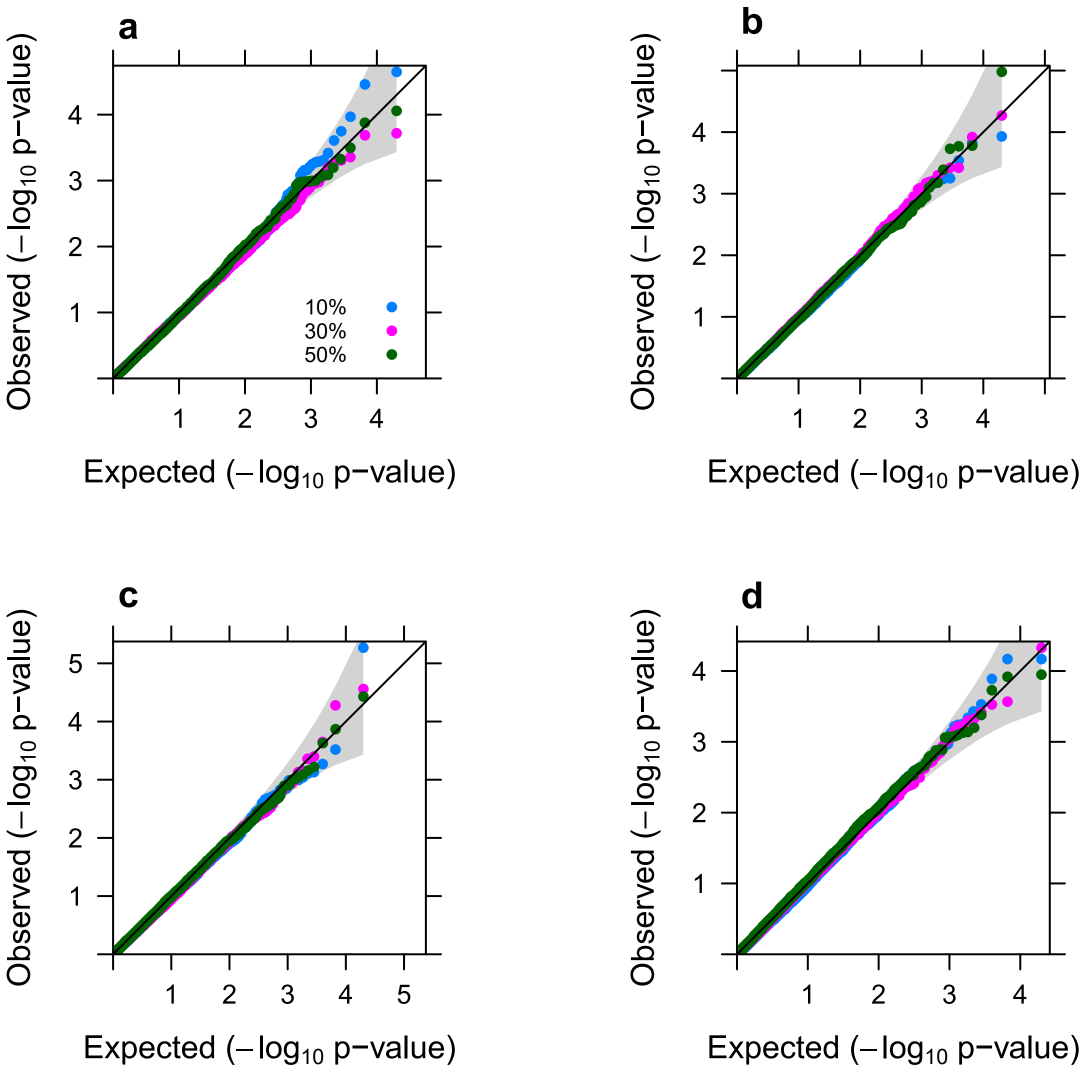


**Fig. 5** Quantile-quantile plot of -log10 p-values from PMR-Egger for testing the causal effect under the null, under various sparse horizontal pleiotropic effect settings. Simulations are performed under different horizontal pleiotropic effect sizes: (a) $\gamma$=0.0001; (b) $\gamma$=0.0005; (c) $\gamma$=0.001; (d) $\gamma$=0.002. In each panel, only a fixed proportion of SNPs (10%, 30%, or 50%) have non-zero horizontal pleiotropic effects. p-values from PMR-Egger behave well across a range of sparse horizontal pleiotropic effect settings.

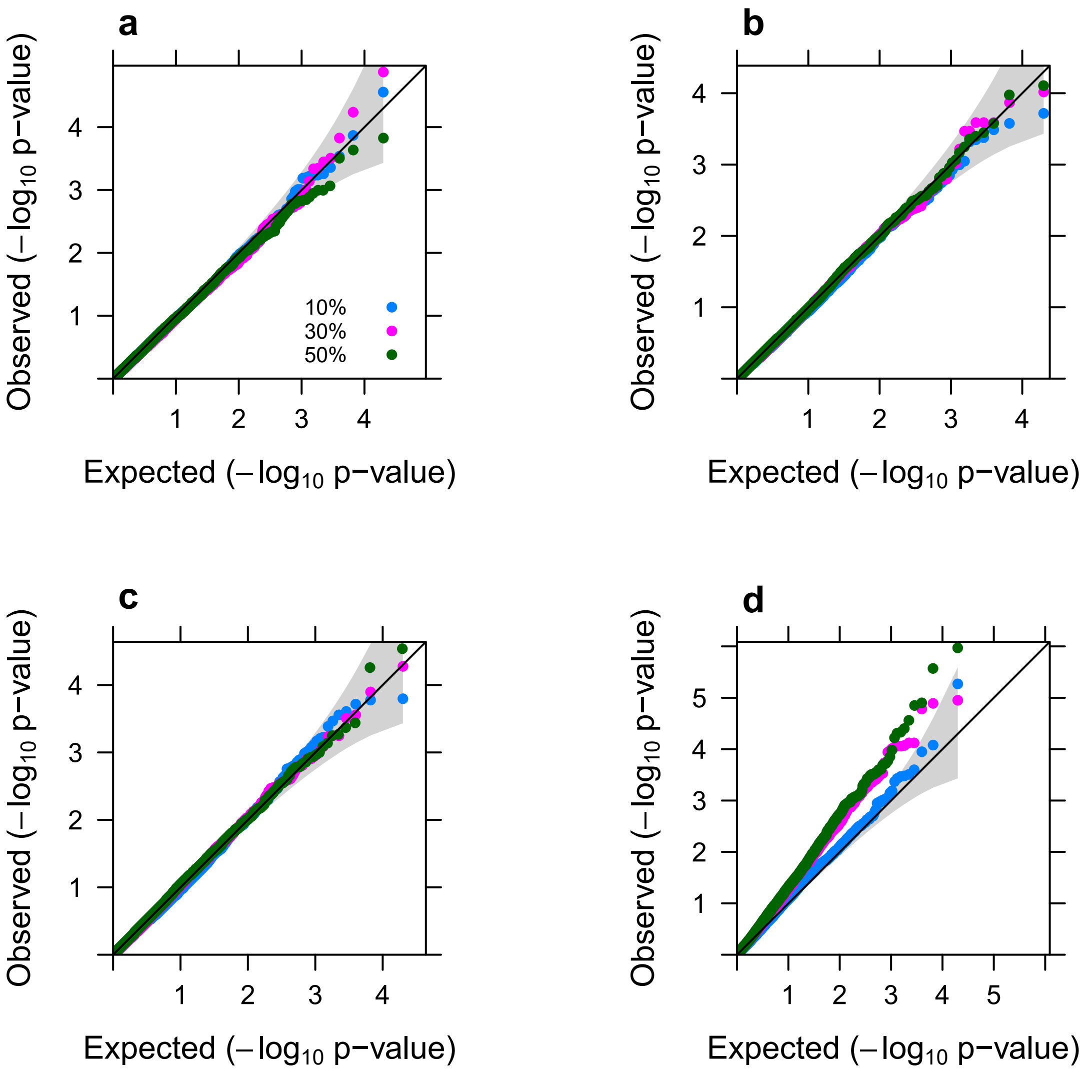


**Fig. 6** Quantile-quantile plot of -log10 p-values from PMR-Egger for testing the causal effect under the null, under various directional horizontal pleiotropic effect assumptions. Simulations are performed under different horizontal pleiotropic effect sizes: (a) $\gamma$=0.0001; (b) $\gamma$=0.0005; (c) $\gamma$=0.001; (d) $\gamma$=0.002. In each panel, a fixed proportion of SNPs (10%, 30%, or 50%) have positive horizontal pleiotropic effects while the remaining proportion of SNPs have negative horizontal pleiotropic effects. p-values from PMR-Egger behave reasonably well across a range of directional or balanced horizontal pleiotropic effect settings, except in the extreme case where horizontal pleiotropic effect size is very large ($\gamma$=0.002) and where the effect size signs across SNPs are approximately balanced.

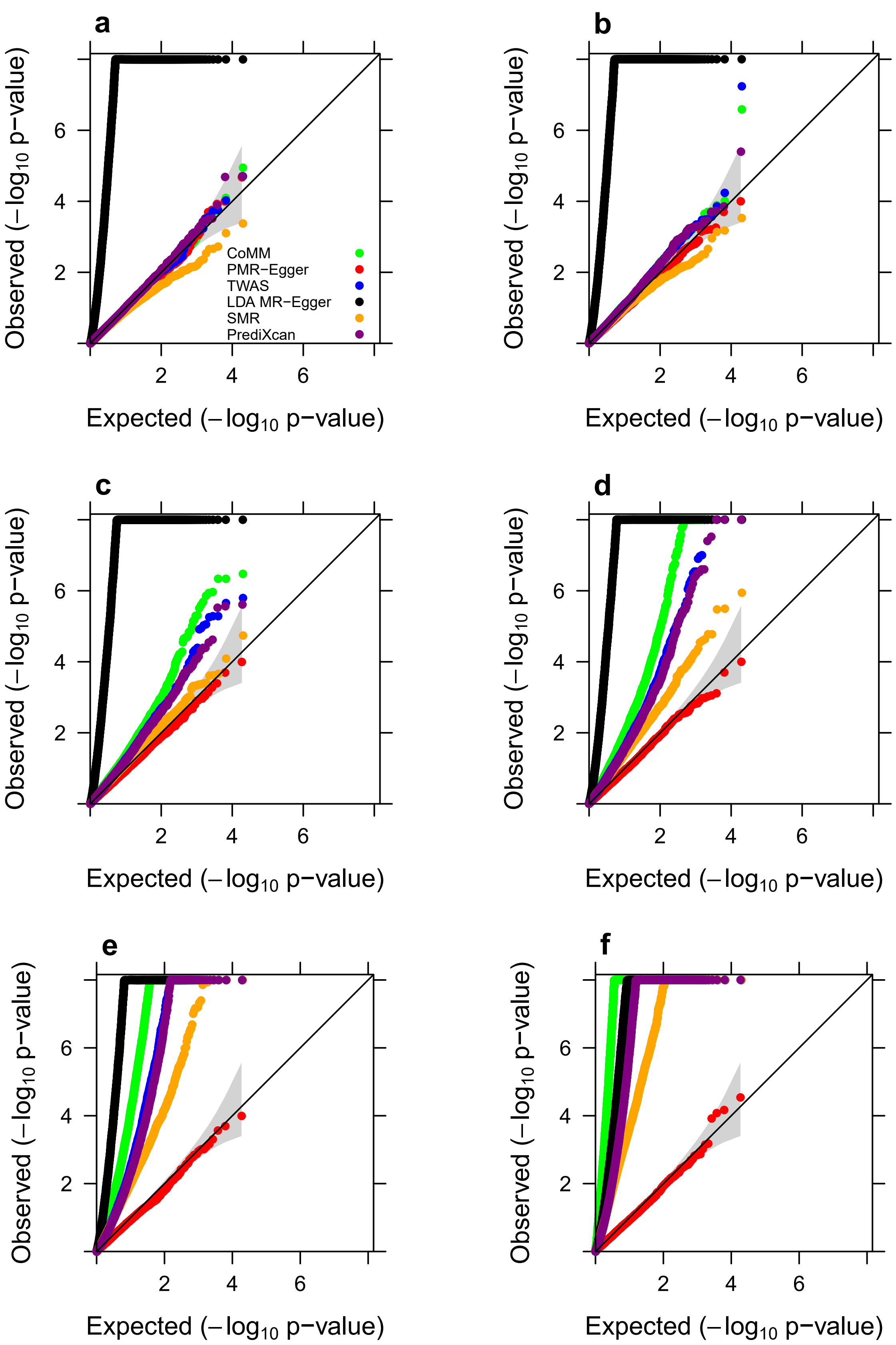


**Fig. 7** Quantile-quantile plot of -log10 p-values from cross-gene simulations for different methods for testing the causal effect either in the absence or in the presence of horizontal pleiotropic effect under null simulations. Compared methods include CoMM (green), PMR-Egger (red), TWAS (blue), LDA MR-Egger (black), SMR (orange), and PrediXcan (purple). Null simulations are performed under different horizontal pleiotropic effect sizes: (a) $\gamma$=0; (b) $\gamma$=0.0001; (c) $\gamma$=0.0002; (d) $\gamma$=0.0003; (e) $\gamma$=0.0005; (f) $\gamma$=0.001. Only p-values from PMR-Egger adhere to the expected diagonal line across a range of horizontal pleiotropic effect sizes.

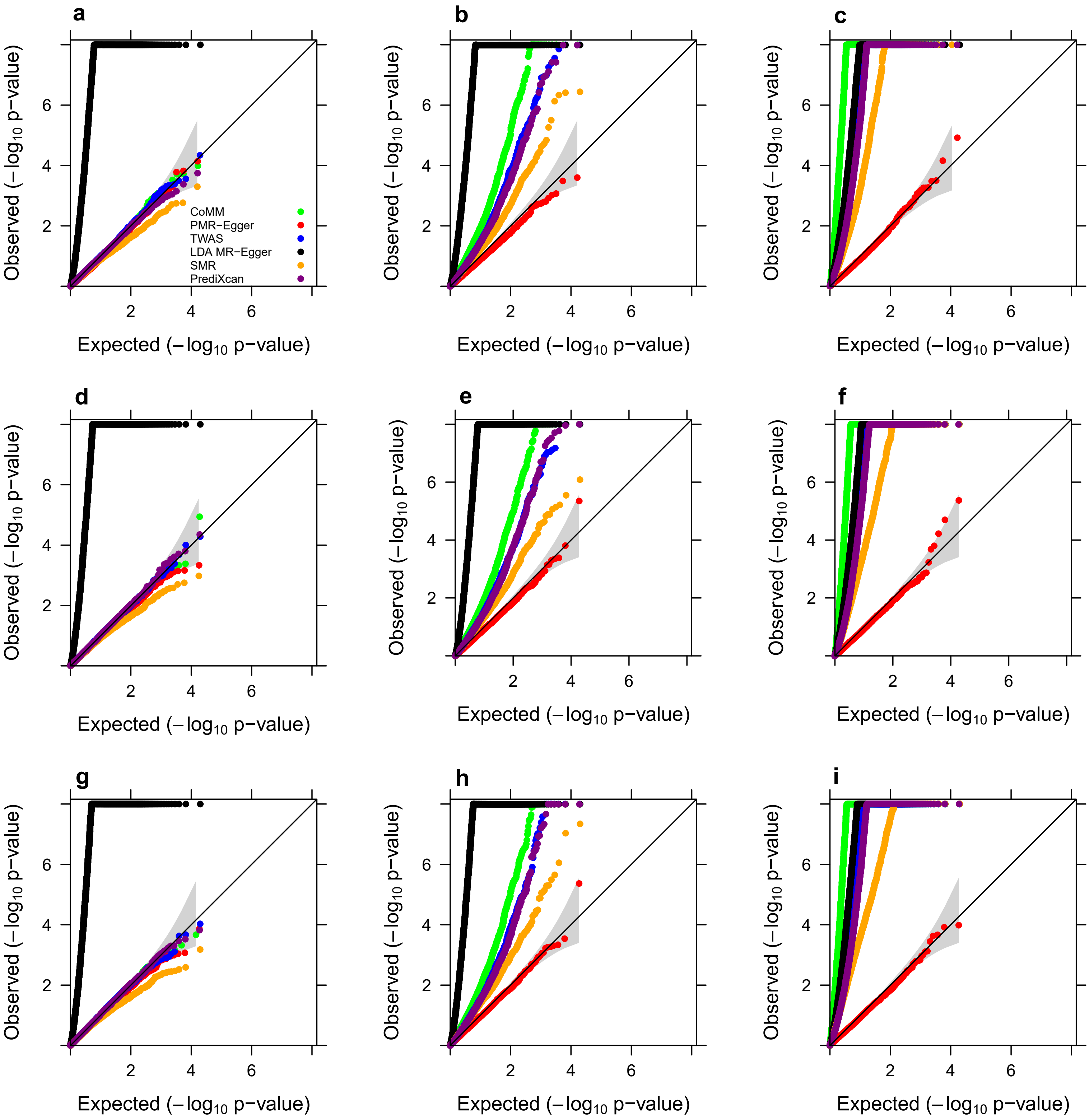


**Fig. 8** Quantile-quantile plot of -log10 p-values from cross-gene simulations of different methods for testing the causal effect under the null simulations, in various sparse settings where only a small proportion of SNPs are associated with the gene expression level. Compared methods include CoMM (green), PMR-Egger (red), TWAS (blue), LDA MR-Egger (black), SMR (orange), and PrediXcan (purple). Simulations are performed either in the absence ($\gamma$=0; a, d, g) or in the presence of horizontal pleiotropic effect ($\gamma$=0.0003; b, e, h; $\gamma$=0.001; c, f, i). Either one SNP (a, b, c), 1% of SNPs (d, e, f), or 10% SNPs (g, h, i) have non-zero effects on gene expression. Only p-values from PMR-Egger adhere to the expected diagonal line across a range of settings.

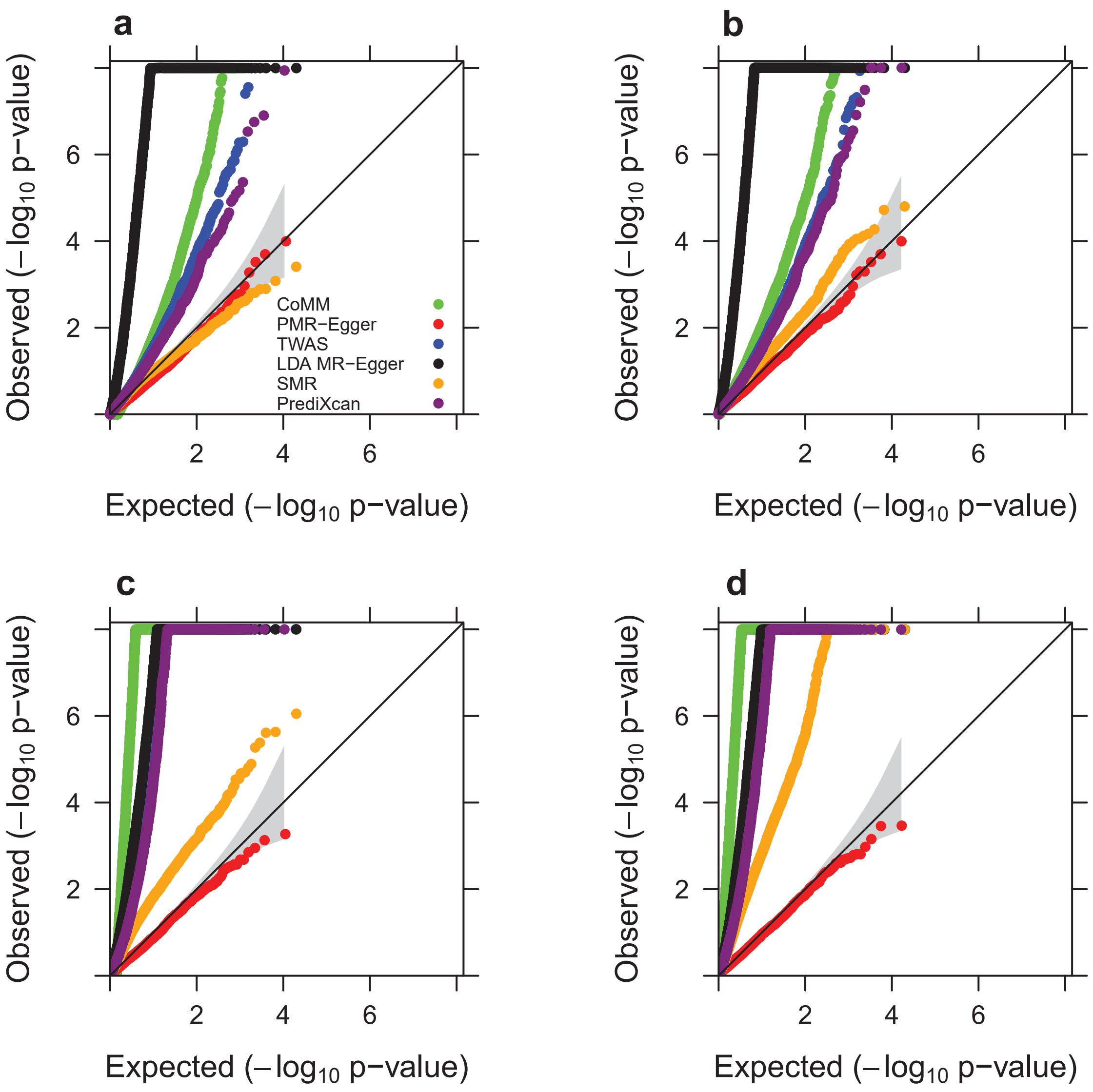


**Fig. 9** Quantile-quantile plot of -log10 p-values from cross-gene simulations of different methods for testing the causal effect under the null, across different gene expression heritability values. Compared methods include CoMM (green), PMR-Egger (red), TWAS (blue), LDA MR-Egger (black), SMR (orange), and PrediXcan (purple). Simulations are performed or in the presence of horizontal pleiotropic effect (γ=0.0003, a, b; γ=0.001, c, d;) with gene expression heritability being either $PVE_{ZX}$=1% (a, c) or $PVE_{ZX}$=5% (b, d). Only p-values from PMR-Egger adhere to the expected diagonal line across a range of settings.

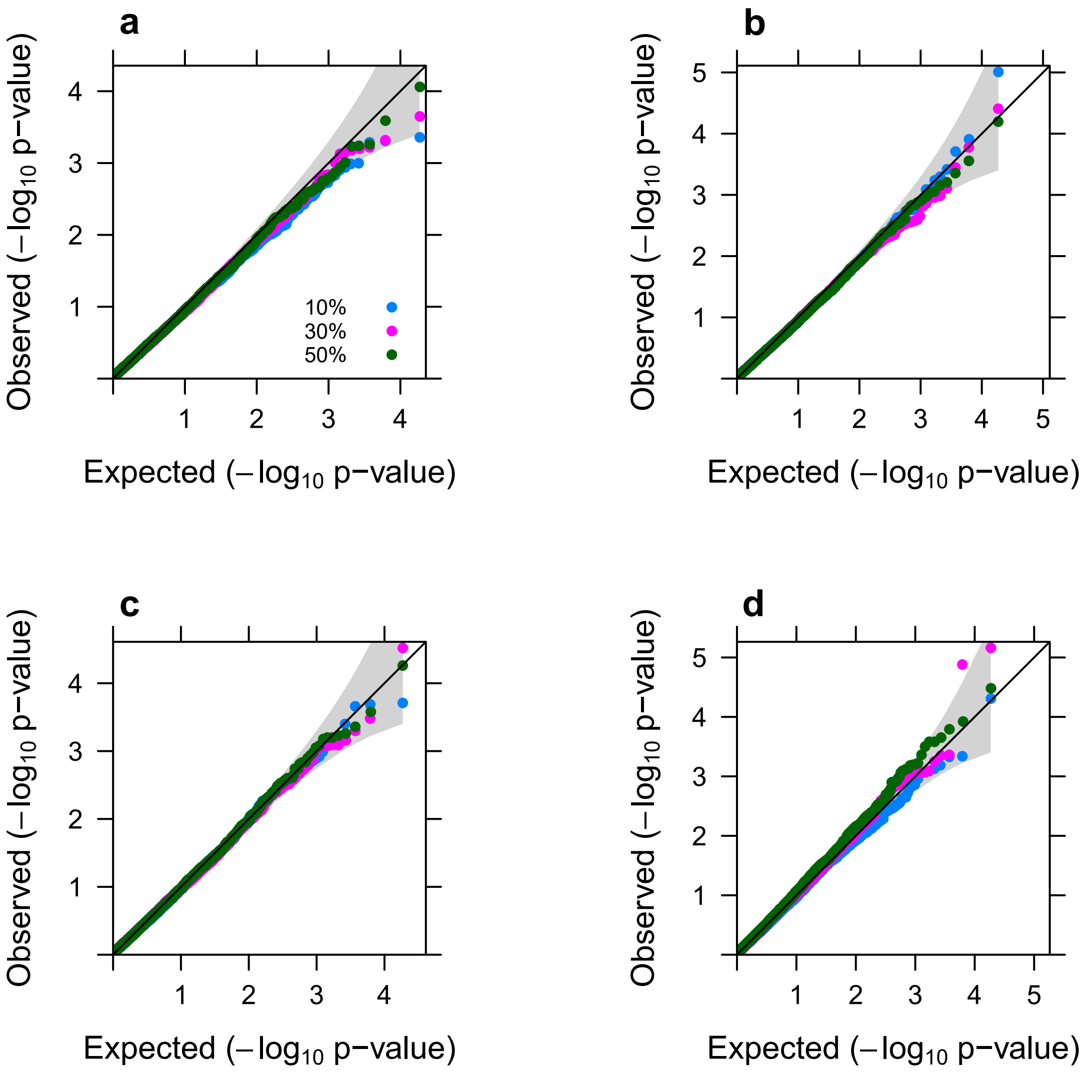


**Fig. 10** Quantile-quantile plot of -log10 p-values from PMR-Egger under cross-gene simulations for testing the causal effect under the null, under various sparse horizontal pleiotropic effect settings. Simulations are performed under different horizontal pleiotropic effect sizes: (a) $\gamma$=0.0001; (b) $\gamma$=0.0005; (c) $\gamma$=0.001; (d) $\gamma$=0.002. In each panel, only a fixed proportion of SNPs (10%, 30%, or 50%) have non-zero horizontal pleiotropic effects. p-values from PMR-Egger behave well across a range of sparse horizontal pleiotropic effect settings.

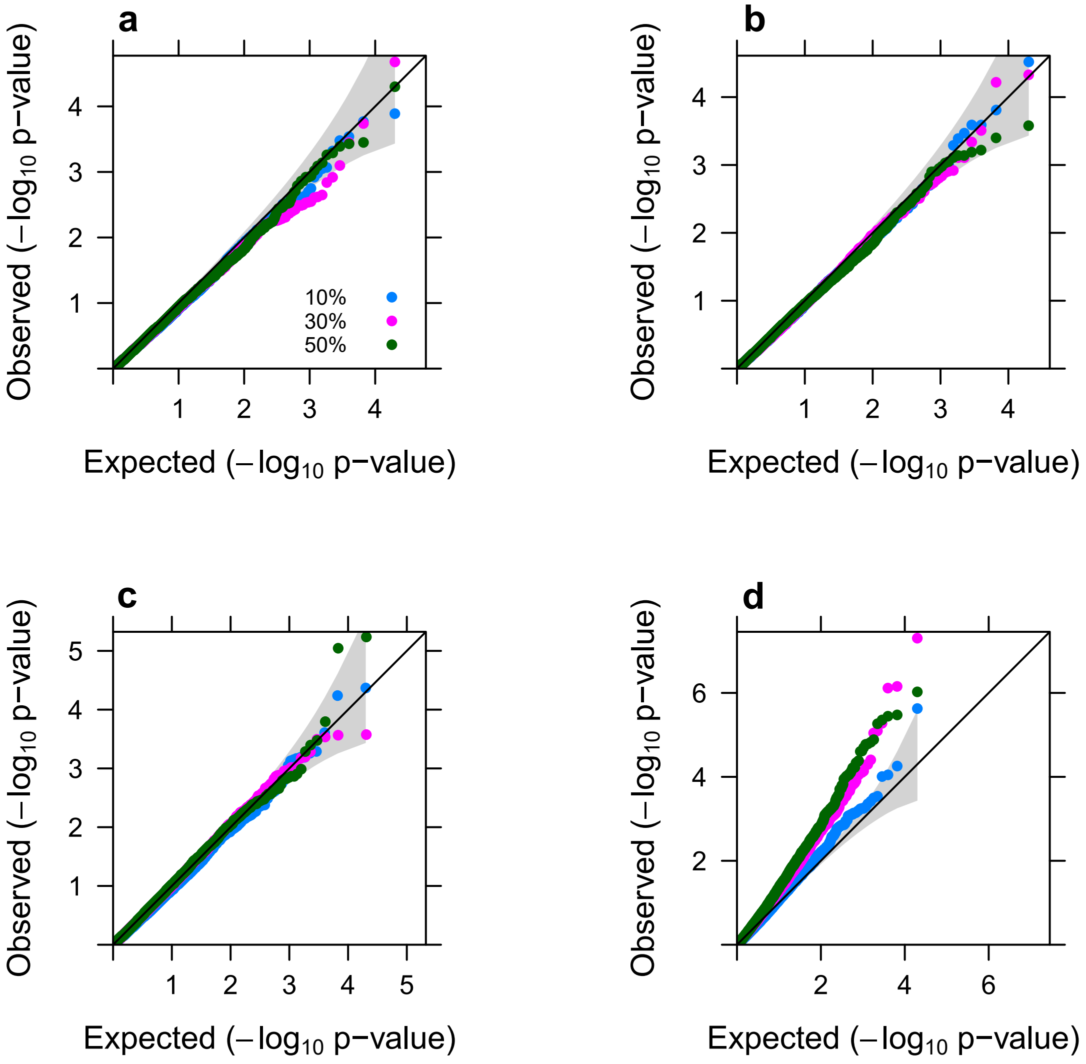


**Fig. 11** Quantile-quantile plot of -log10 p-values from PMR-Egger under cross-gene simulations for testing the causal effect under the null, under various directional horizontal pleiotropic effect assumptions. Simulations are performed under different horizontal pleiotropic effect sizes: (a) $\gamma$=0.0001; (b) $\gamma$=0.0005; (c) $\gamma$=0.001; (d) $\gamma$=0.002. In each panel, a fixed proportion of SNPs (10%, 30%, or 50%) have positive horizontal pleiotropic effects while the remaining proportion of SNPs have negative horizontal pleiotropic effects. p-values from PMR-Egger behave reasonably well across a range of directional or balanced horizontal pleiotropic effect settings, except in the extreme case where horizontal pleiotropic effect size is very large ($\gamma$=0.002) and where the effect size signs across SNPs are approximately balanced.

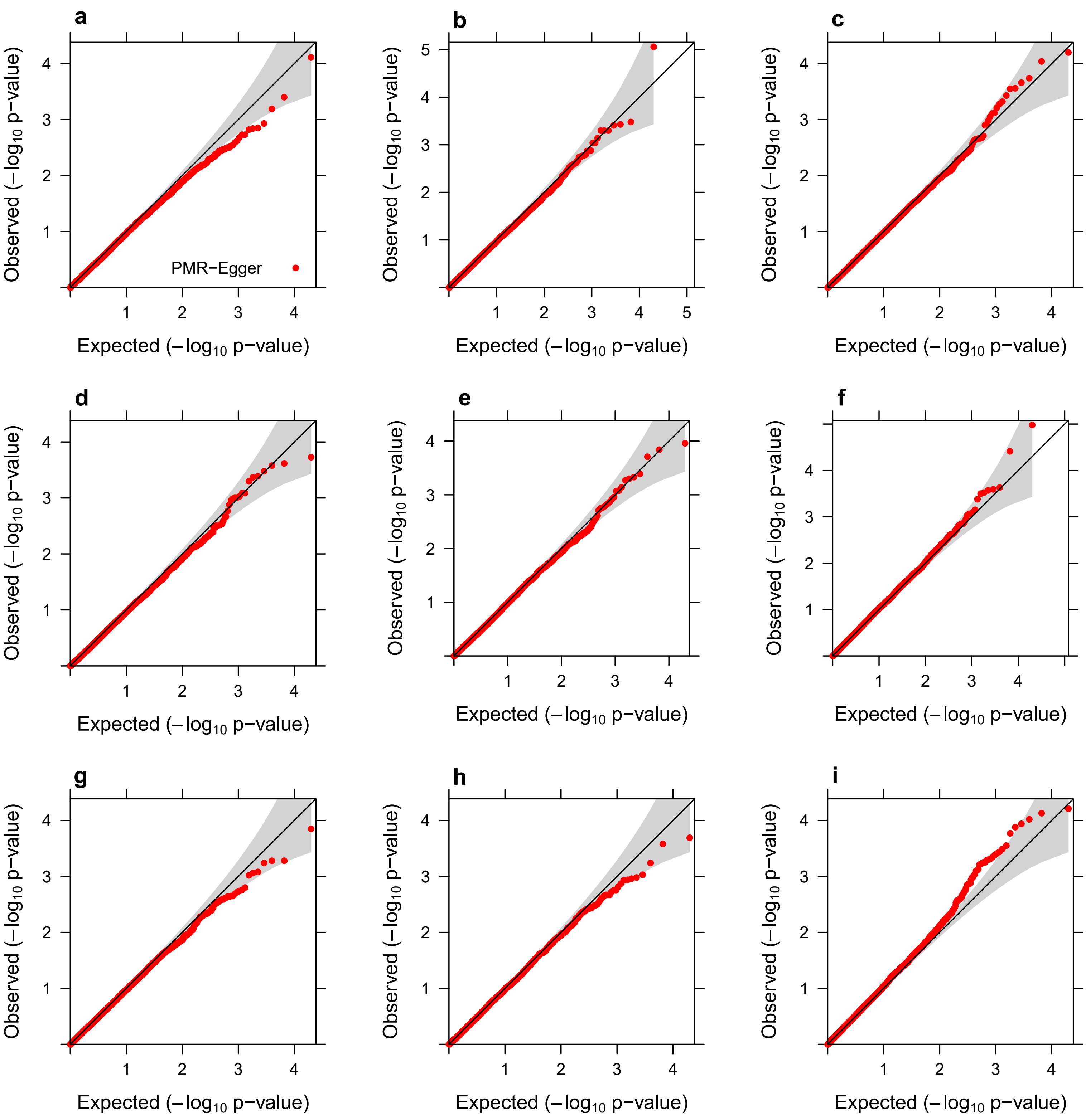


**Fig. 12** Quantile-quantile plot of -log10 p-values from cross-gene simulations of PMR-Egger for testing the causal effect under null simulations, when randomly flipping the encoding at a fixed proportion of cis-SNPs, the proportion equals 10% (a, b, c), 30% (d, e, f) and 50% (g, h, i). Simulations are performed under different pleiotropy effect sizes, $\gamma$=0 (a, d, g), $\gamma$=0.0005 (b, e, h) or $\gamma$=0.001 (c, f, i). P-values from PMR-Egger behave reasonably well across most of the scenarios, and it is a little inflated in the case where the flipping proportion reached 50% and horizontal pleiotropic effect size is large ($\gamma$=0.001).

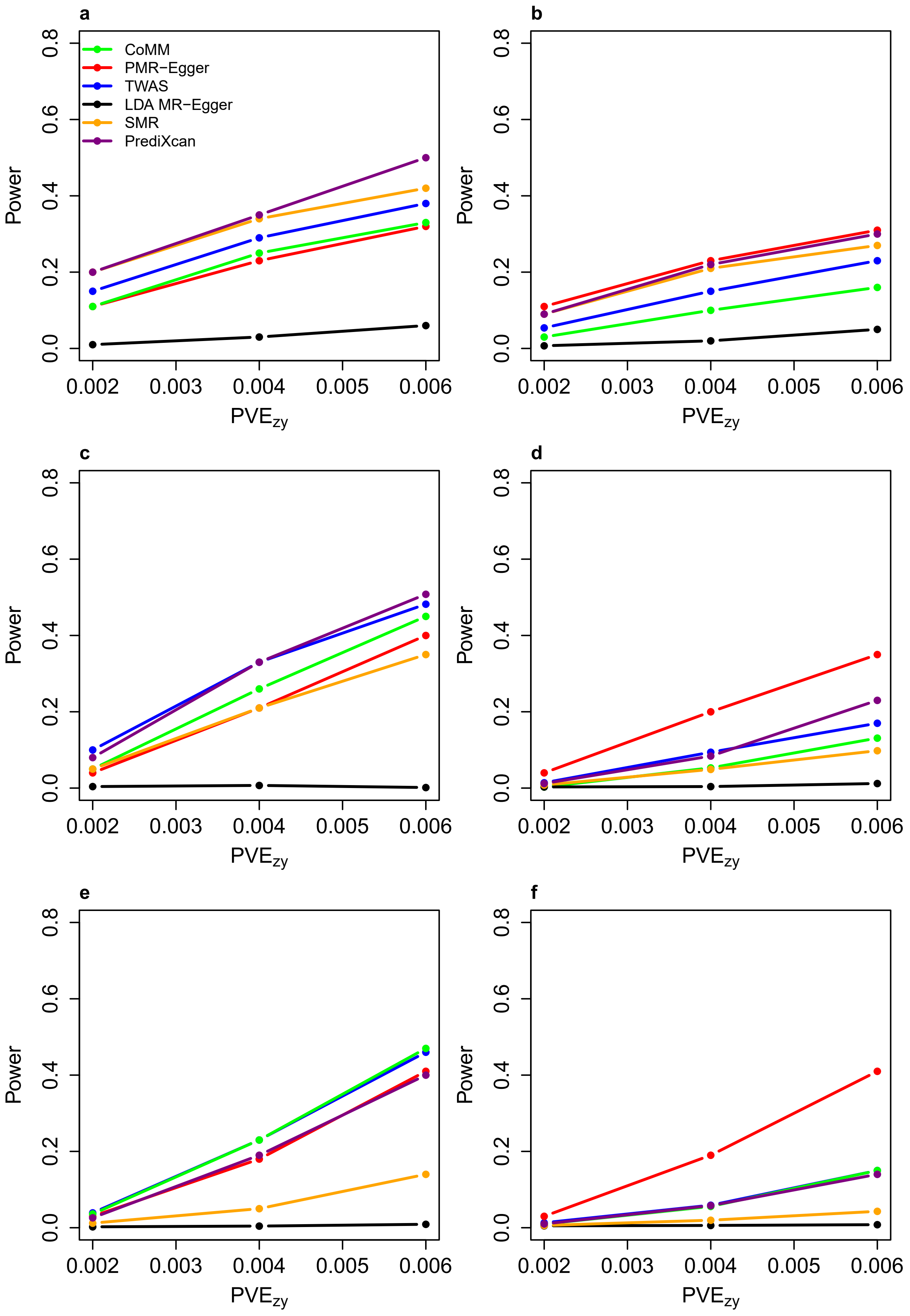


**Fig. 13** Power for testing causal effect by different methods in various sparse settings where only a small proportion of SNPs are associated with the gene expression level. Power (y-axis) at a false discovery rate of 0.1 to detect the causal effect is plotted against different causal effect size characterized by PVE$_{zy}$ (x-axis). Compared methods include CoMM (green), PMR-Egger (red), TWAS (blue), LDA MR-Egger (black), SMR (orange), and PrediXcan (purple). Simulations are performed either in the absence ($\gamma$=0; a, c, e) or in the presence of horizontal pleiotropic effect ($\gamma$=0.001; b, d, f). Either one SNP (a, b), 1% of SNPs (c, d), or 10% SNPs (e, f) have non-zero effects on gene expression.

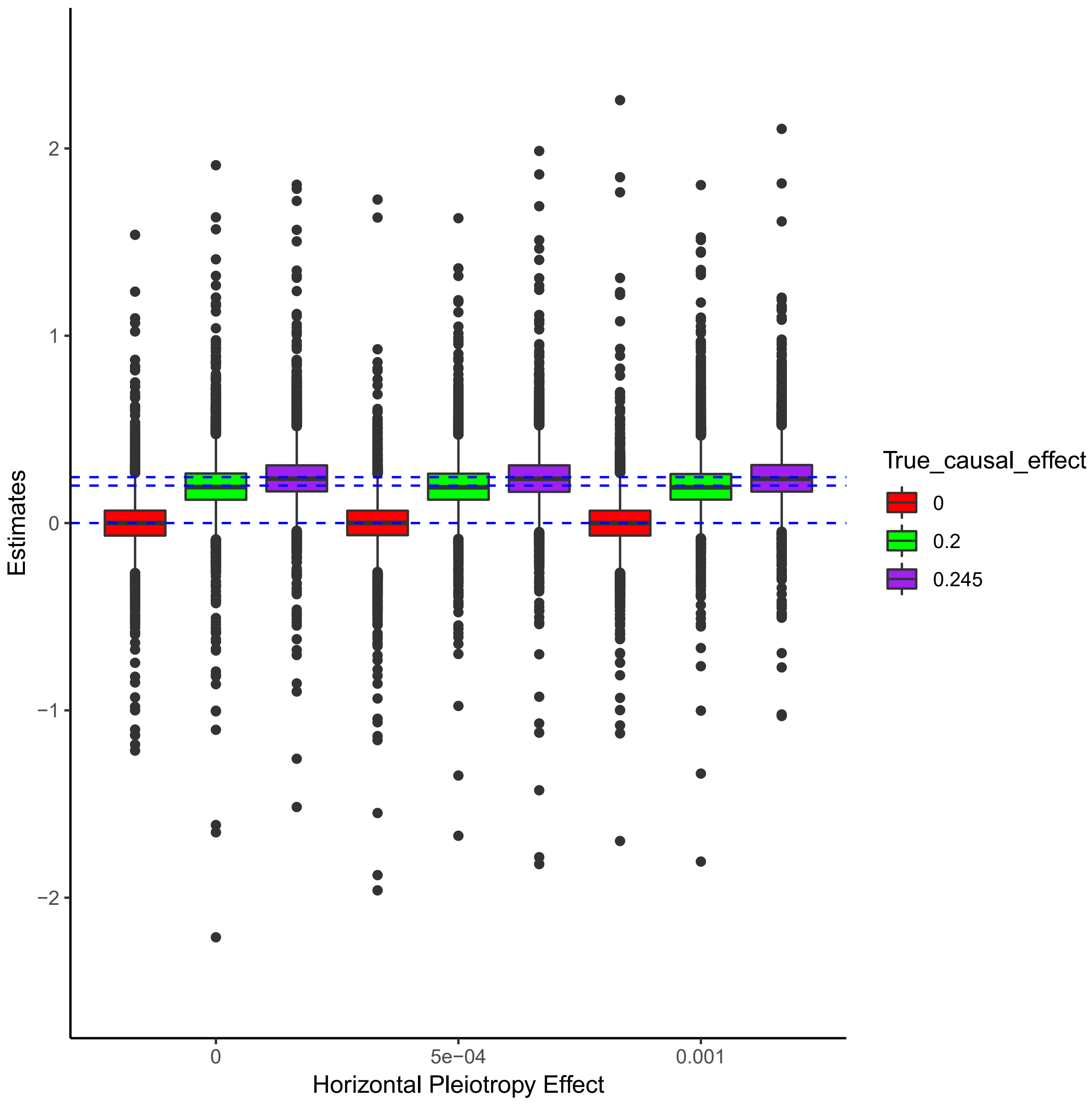


**Fig. 14** Boxplot displays causal effect estimates by PMR-Egger in the absence or presence of horizontal pleiotropic effect. Simulations are performed under different horizontal pleiotropic effect sizes (x-axis: $\gamma$=0, $\gamma$=0.0005 or $\gamma$=0.001). For each horizontal pleiotropic effect size, we examined three true causal effect sizes $\alpha$=0 (red), 0.2 (green), or 0.245 (purple), which corresponds to $PVE_{zy}$=0, 0.4% and 0.6%, respectively. The horizontal red dashed lines represent the three true values of $\alpha$. PMR-Egger produces approximately unbiased causal effect size estimates across different scenarios. 10000 replicates are included for each simulation scenario.

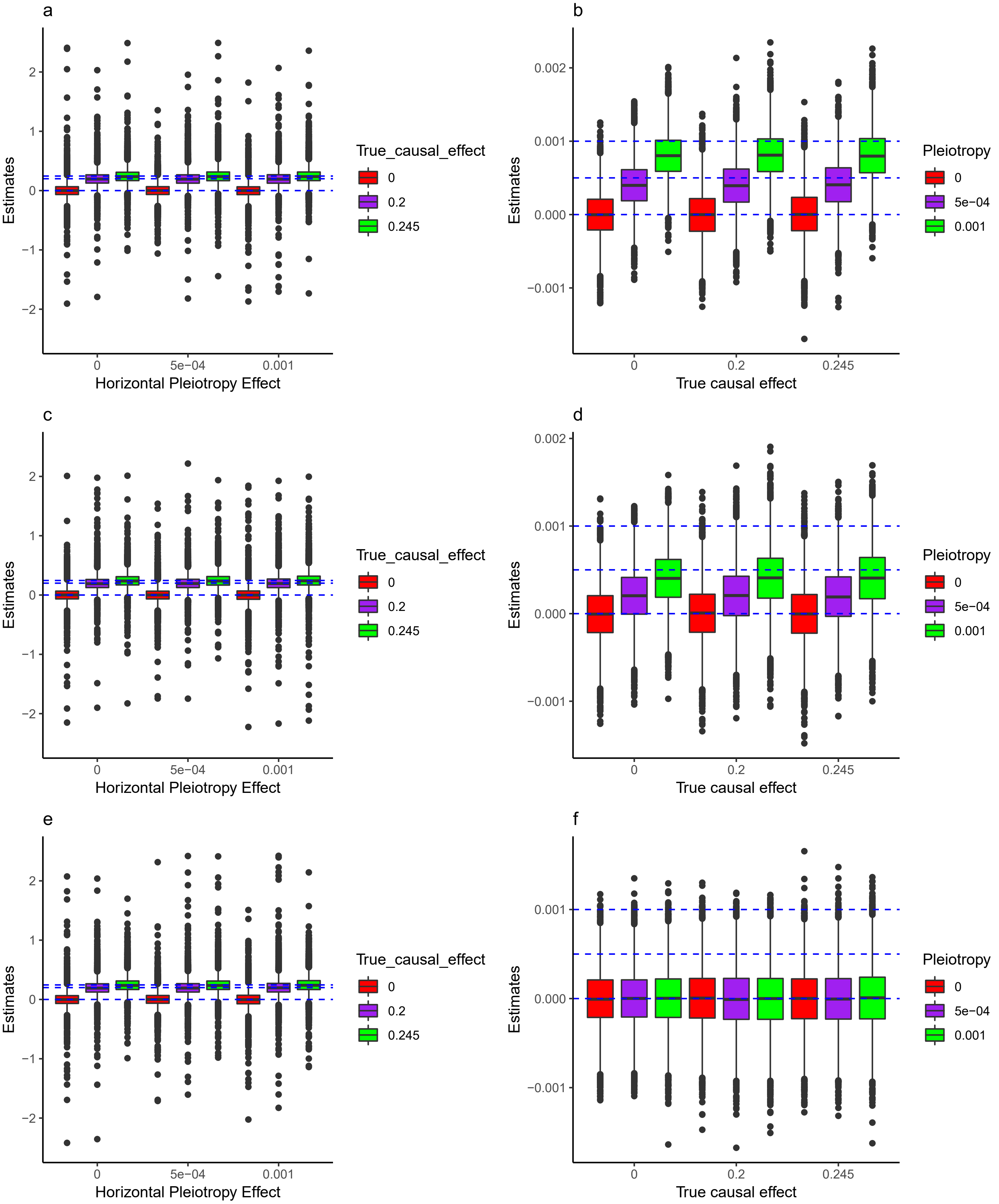


**Fig. 15** Boxplot displays causal and horizontal pleiotropic effect estimates in cross-gene simulations by PMR-Egger when randomly flipping the encoding at a fixed proportion of cis-SNPs, the proportion equals 10% (a, b), 30% (c, d) and 50% (e, f). Simulations are performed under different causal effect sizes (α=0, α=0.2 or α=0.245) and pleiotropy effect sizes (γ=0, γ=0.0005 or γ=0.001). The horizontal blue dashed lines represent the three true values of α (a, c, e) and γ (b, d, f). 10,000 replicates are included for each simulation scenario. The causal effect estimates are always unbiased regardless of the pleiotropy effect size and the proportion of flipping coding. When there is no pleiotropy effect (γ=0), the estimation of pleiotropy effect is always unbiased regardless of the causal effect size and the proportion of flipping encoding. However, in the presence of pleiotropy effect (γ=0.0005,0.001), PMR-Egger seems to underestimate the pleiotropy effect, and more so with the increasing proportion of flipping encoding. As expected, when the proportion of flipping encoding is 50%, the estimation of pleiotropy effect is always close to 0. Furthermore, the strength of underestimation seems unrelated with the causal effect size.

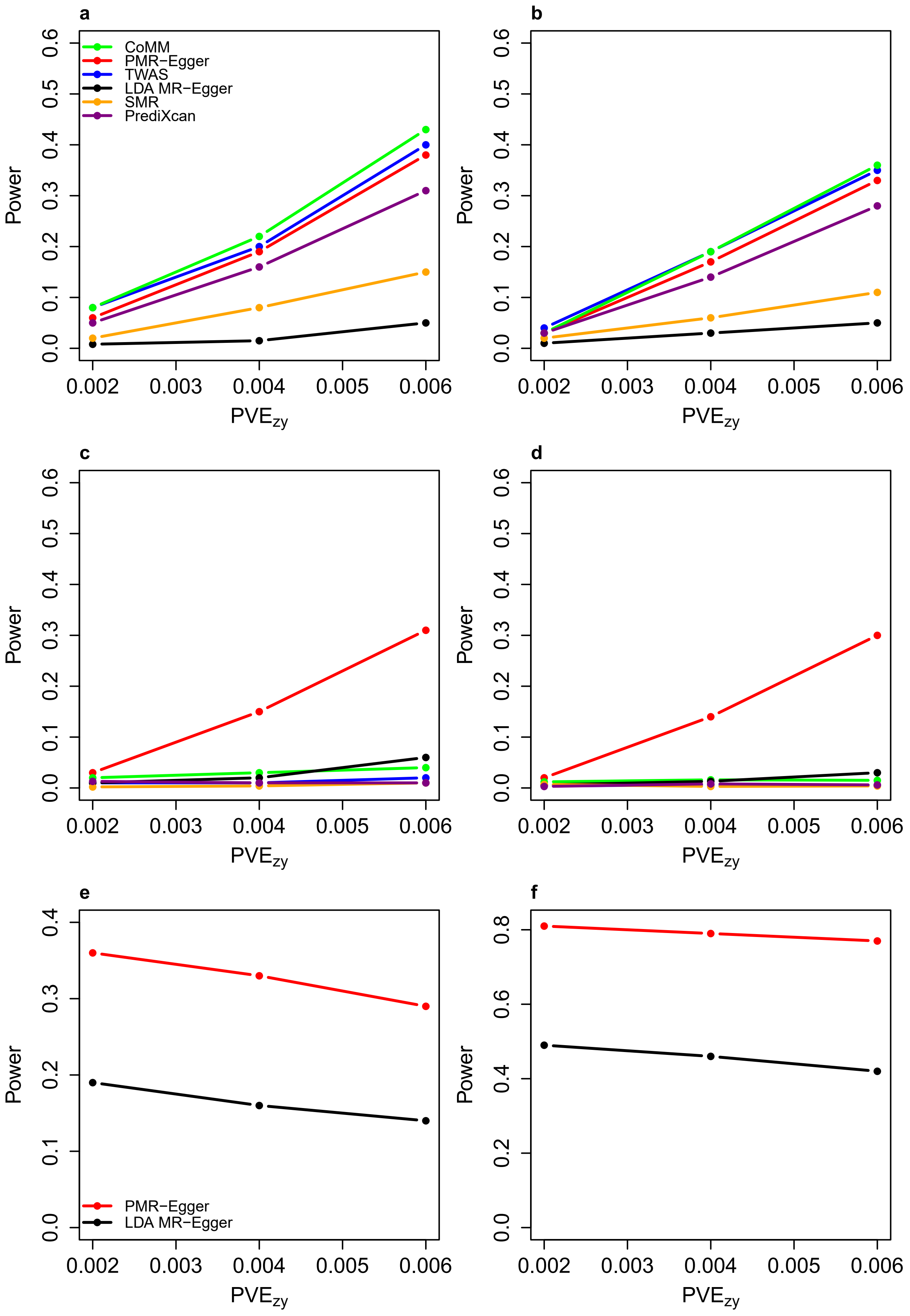


**Fig. 16** Power of different methods under various cross-gene simulation scenarios. Power (y-axis) at a false discovery rate of 0.1 to detect the causal effect (a-d) or the horizontal pleiotropic effect (e-f) is plotted against different causal effect size characterized by $PVE_{zy}$ (x-axis). Compared methods include CoMM (green), PMR-Egger (red), TWAS (blue), LDA MR-Egger (black), SMR (orange), and PrediXcan (purple). Simulations are performed under different horizontal pleiotropic effect sizes: (a) $\gamma$=0; (b) $\gamma$=0.0001; (c, e) $\gamma$=0.0005; (d, f) $\gamma$=0.001.

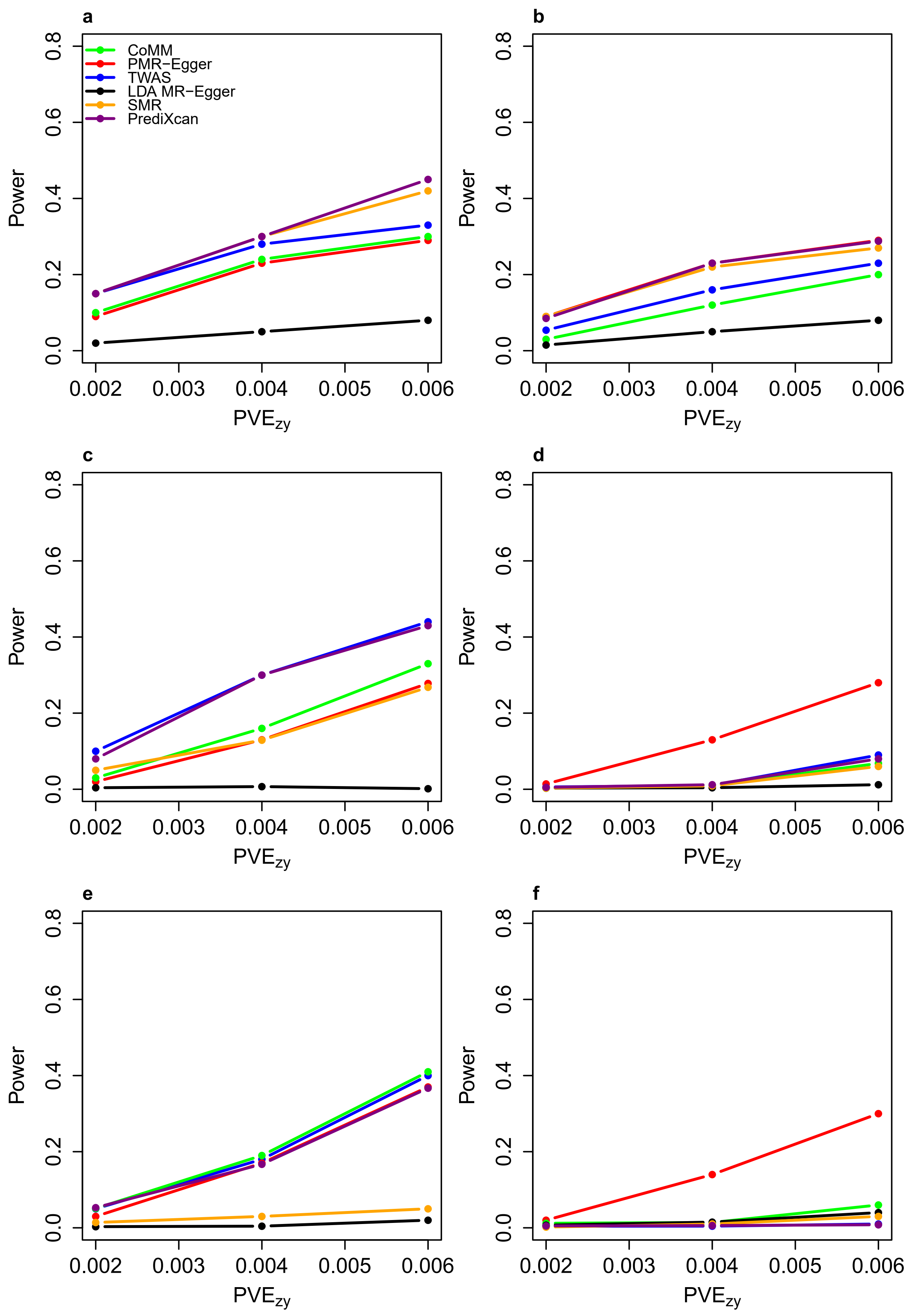


**Fig. 17** Power for testing causal effect by different methods under various cross-gene simulation sparse settings where only a small proportion of SNPs are associated with the gene expression level. Power (y-axis) at a false discovery rate of 0.1 to detect the causal effect is plotted against different causal effect size characterized by $PVE_{zy}$ (x-axis). Compared methods include CoMM (green), PMR-Egger (red), TWAS (blue), LDA MR-Egger (black), SMR (orange), and PrediXcan (purple). Simulations are performed either in the absence ($\gamma$=0; a, c, e) or in the presence of horizontal pleiotropic effect ($\gamma$=0.001; b, d, f). Either one SNP (a, b), 1% of SNPs (c, d), or 10% SNPs (e, f) have non-zero effects on gene expression.

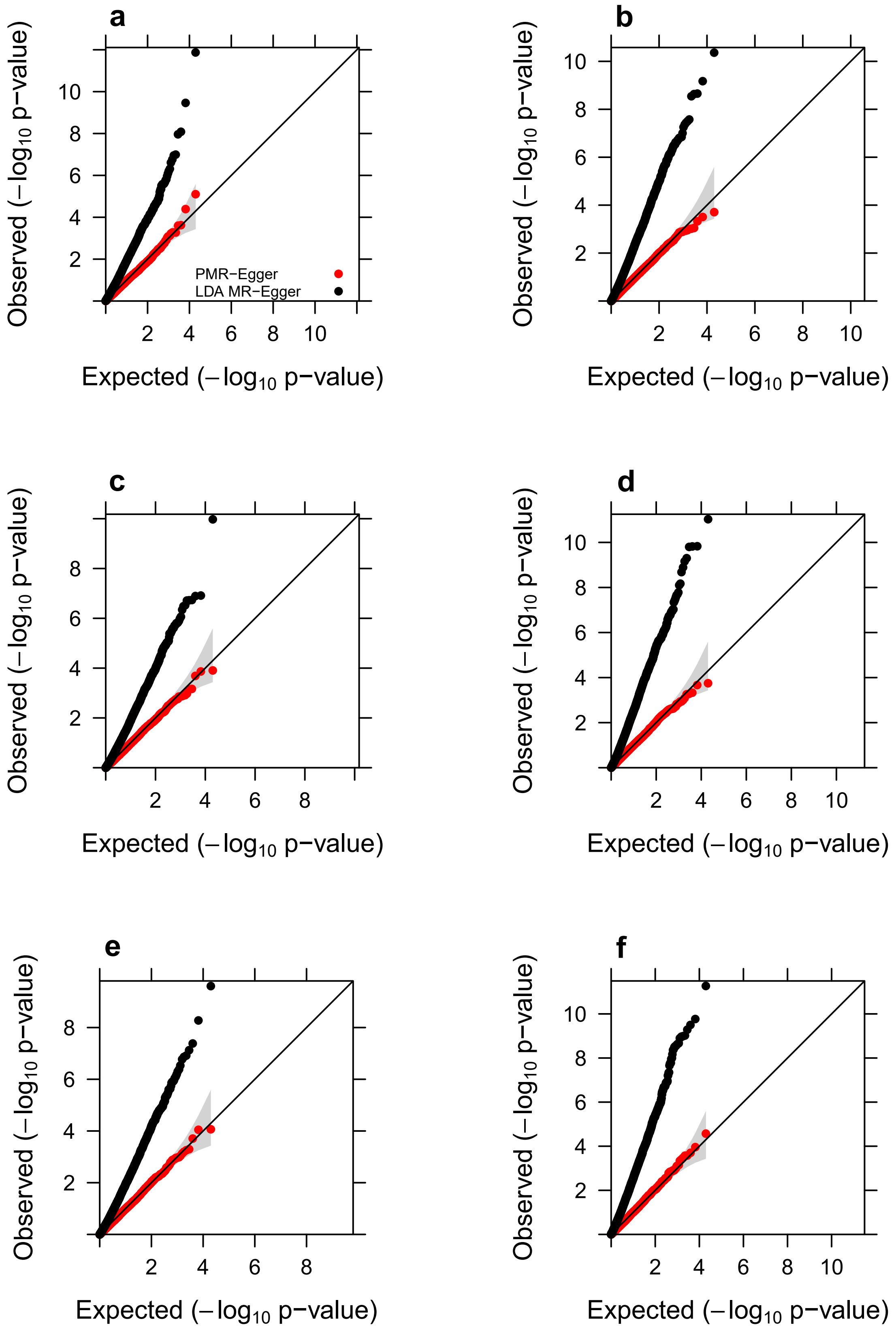


**Fig. 18** Quantile-quantile plot of -log10 p-values from different methods for testing the horizontal pleiotropic effect under null simulations, in various sparse settings where only a small proportion of SNPs are associated with the gene expression level. Compared methods include PMR-Egger (red) and LDA MR-Egger (black). Simulations are performed either in the absence ($PVE_{zy}$=0; a, c, e) or in the presence of causal effect ($PVE_{zy}$=0.6%; b, d, f). Either one SNP (a, b), 1% of SNPs (c, d), or 10% SNPs (e, f) have non-zero effects on gene expression. Only p-values from PMR-Egger adhere to the expected diagonal line across a range of settings. Note that we do not include MR-PRESSO into comparison here due to the relatively higher computation burden and it is difficult in MR-PRESSO to pre-specify the number of simulated expected distribution and obtain the exact p values.

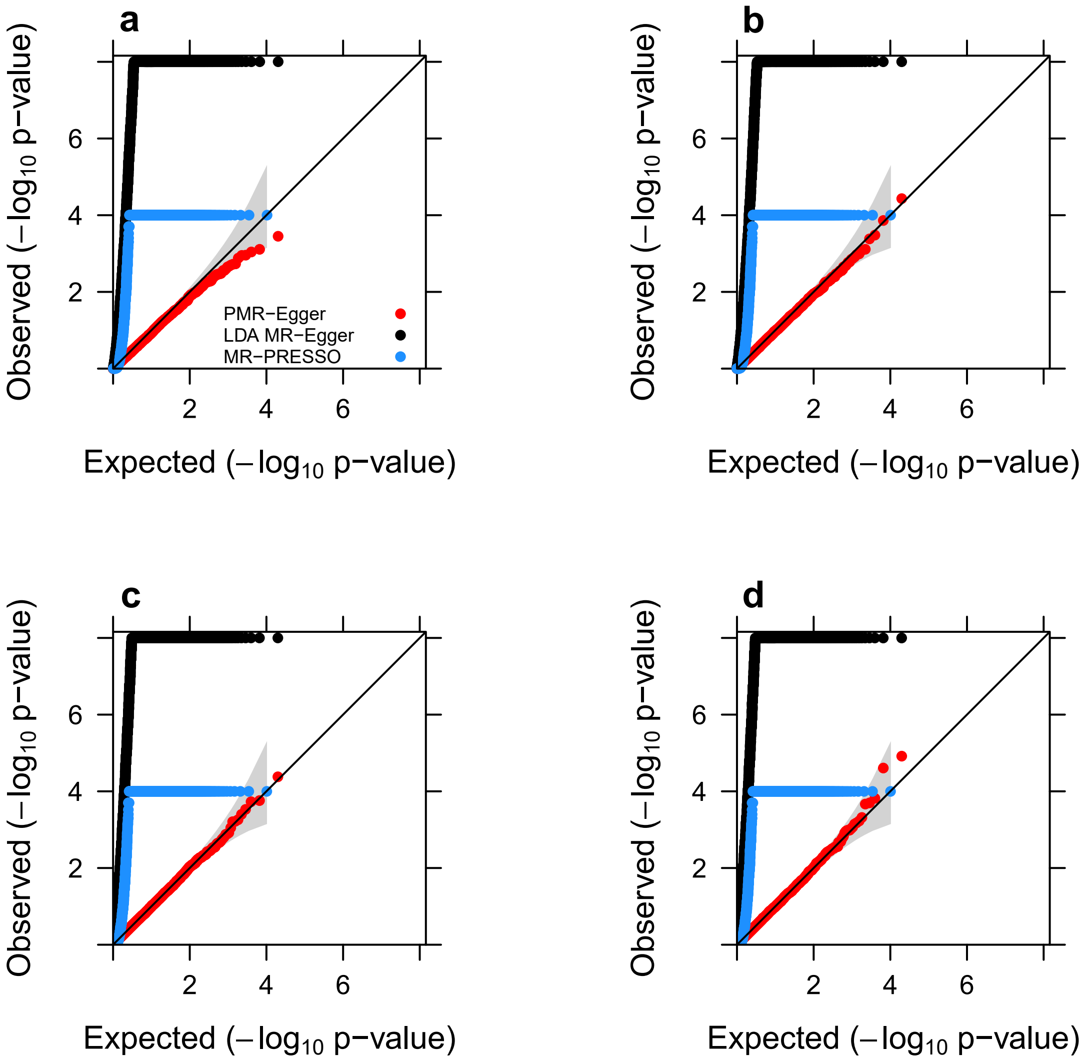


**Fig. 19** Quantile-quantile plot of -log10 p-values from cross-gene simulations of different methods for testing the horizontal pleiotropic effect either in the absence or in the presence of causal effect under null simulations. Compared methods include PMR-Egger (red), LDA MR-Egger (black), and MR-PRESSO (dodger blue). Null simulations are performed under different causal effect sizes characterized by $PVE_{zy}$: (a) $PVE_{zy}$=0; (b) $PVE_{zy}$=0.2%; (c) $PVE_{zy}$=0.4%; and (d) $PVE_{zy}$=0.6%. Only p-values from PMR-Egger adhere to the expected diagonal line across a range of horizontal pleiotropic effect sizes. Due to heavy computational burden, we are only able to run 10,000 permutations for MR-PRESSO. Therefore, the minimal p-value from MR-PRESSO is $10^{-4}$.

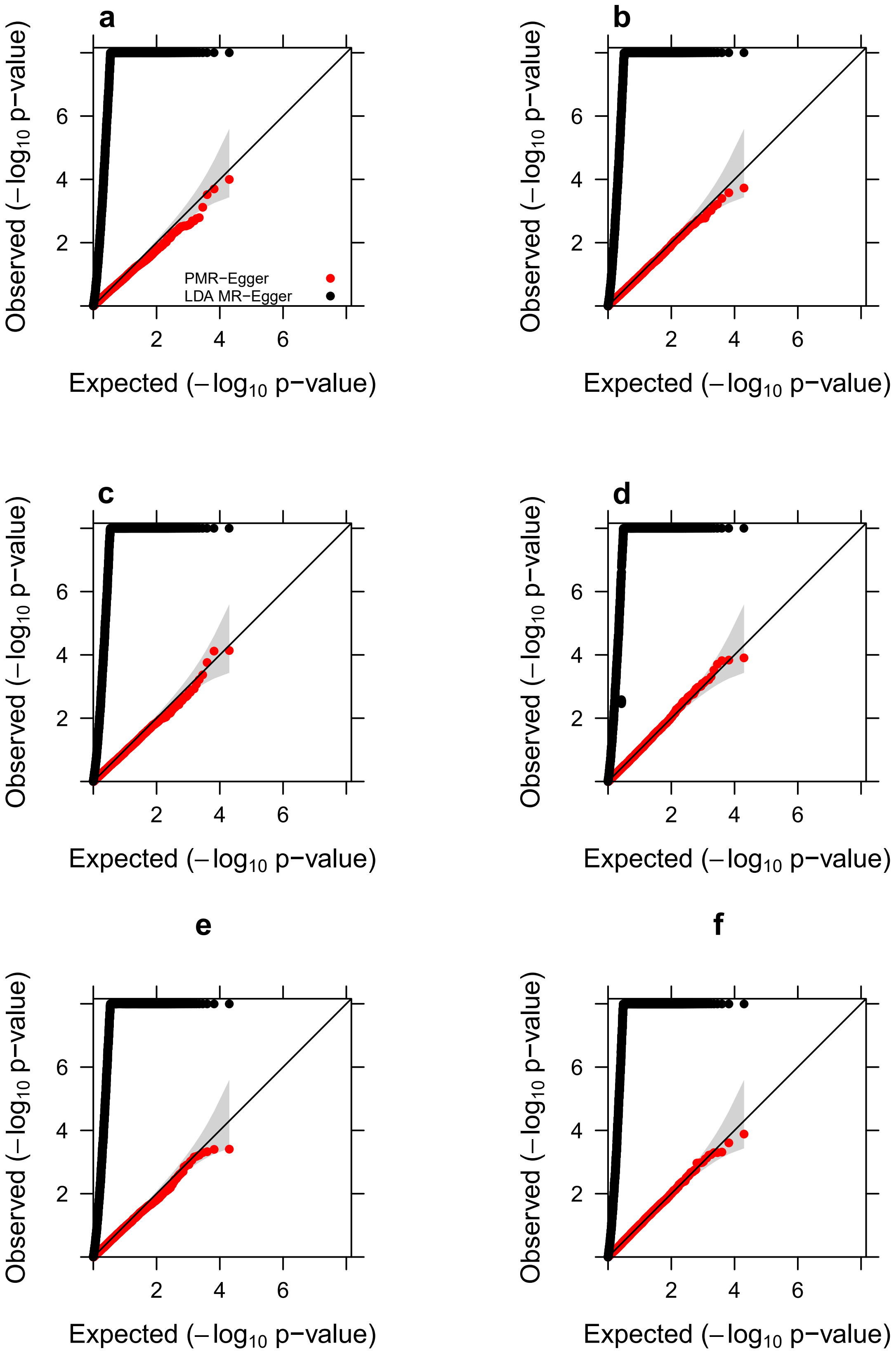


**Fig. 20** Quantile-quantile plot of -log10 p-values from cross-gene simulations of different methods for testing the horizontal pleiotropic effect under null simulations, in various sparse settings where only a small proportion of SNPs are associated with the gene expression level. Compared methods include PMR-Egger (red) and LDA MR-Egger (black). Simulations are performed either in the absence ($PVE_{zy}$=0; a, c, e) or in the presence of causal effect ($PVE_{zy}$=0.6%; b, d, f). Either one SNP (a, b), 1% of SNPs (c, d), or 10% SNPs (e, f) have non-zero effects on gene expression. Only p-values from PMR-Egger adhere to the expected diagonal line across a range of settings. Note that we do not include MR-PRESSO into comparison here due to the relatively higher computation burden and it is difficult in MR-PRESSO to pre-specify the number of simulated expected distribution and obtain the exact p values.

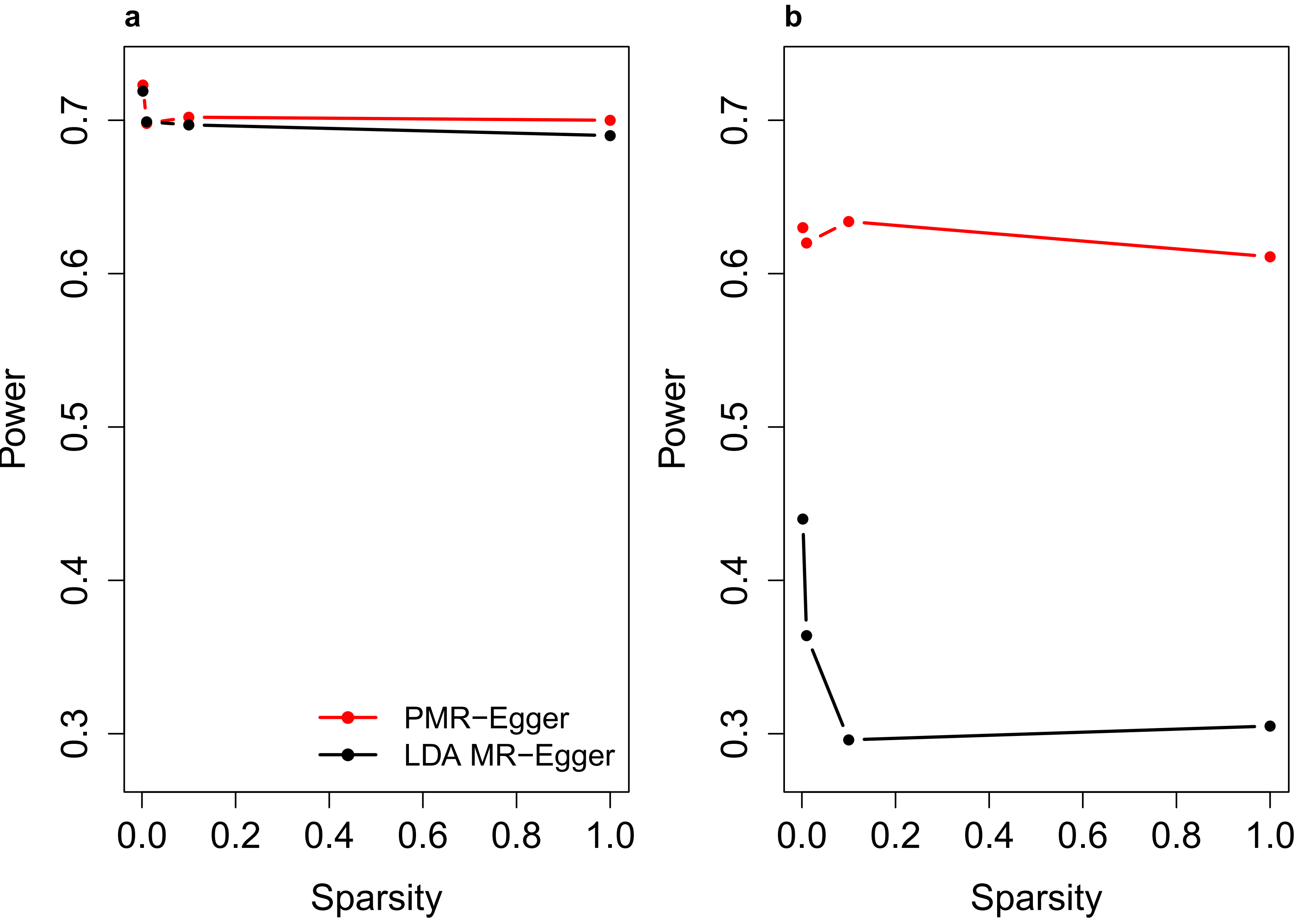


**Fig. 21** Power of different methods for identifying horizontal pleiotropic effect, in various sparse settings where only a small proportion of SNPs are associated with the gene expression level. Compared methods include PMR-Egger (red) and LDA MR-Egger (black). Power (y-axis) at a false discovery rate of 0.1 to detect the horizontal pleiotropic effect is plotted against sparsity levels, either in the absence ($PVE_{zy}$=0; a) or in the presence of causal effect ($PVE_{zy}$=0.6%; b). In terms of sparsity level (x-axis), either one SNP, 1% SNPs, 10% SNPs and 100% SNPs have non-zero effects on gene expression.

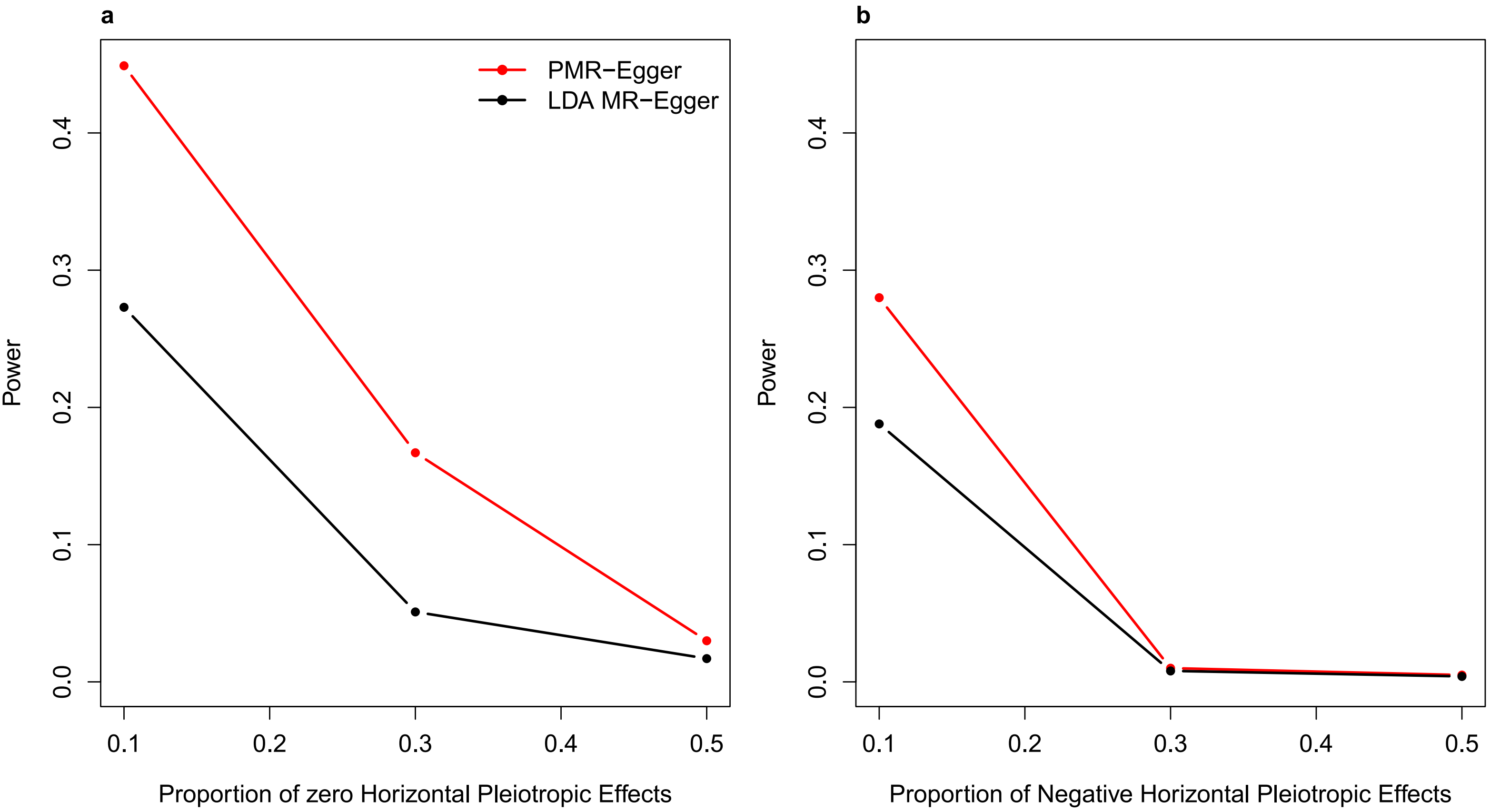


**Fig. 22** Power of different methods for identifying horizontal pleiotropic effect under various model misspecifications of the horizontal pleiotropic effect. Compared methods include PMR-Egger (red) and LDA MR-Egger (black). On both panels, power (y-axis) to detect the horizontal pleiotropic effect is measured at a false discovery rate of 0.1 for a fixed horizontal pleiotropic effect size $\gamma$=0.001. (a) Power is plotted against the proportion of SNPs displaying zero horizontal pleiotropy effects (10%, 30%, or 50%). (b) Power is plotted against the proportion of SNPs exhibiting negative horizontal pleiotropy effects (10%, 30%, or 50%), whereas the remaining proportion of SNPs exhibiting positive effects.

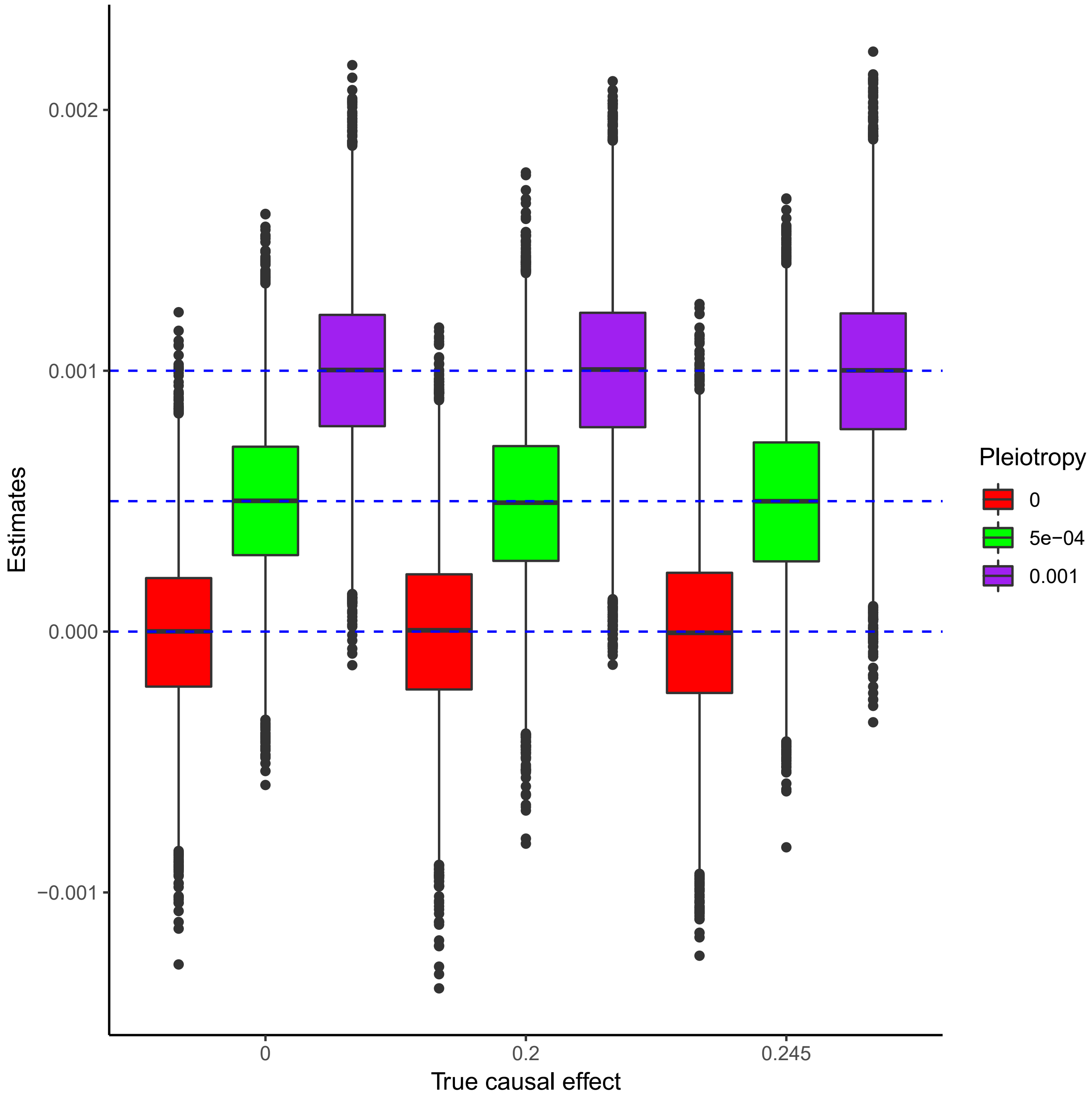


**Fig. 23** Boxplot displays horizontal pleiotropic effect estimates by PMR-Egger in the absence or presence of causal effect. Simulations are performed under different causal effect sizes (x-axis: $\alpha$=0, $\alpha$=0.2 or $\alpha$=0.245). For each causal effect size, we examined three true horizontal pleiotropic effect sizes $\gamma$=0 (red), 0.0005 (green), or 0.001 (purple). The horizontal red dashed lines represent the three true values of $\gamma$. PMR-Egger produces approximately unbiased pleiotropy effect size estimates across different scenarios. 10,000 replicates are included for each simulation scenario.

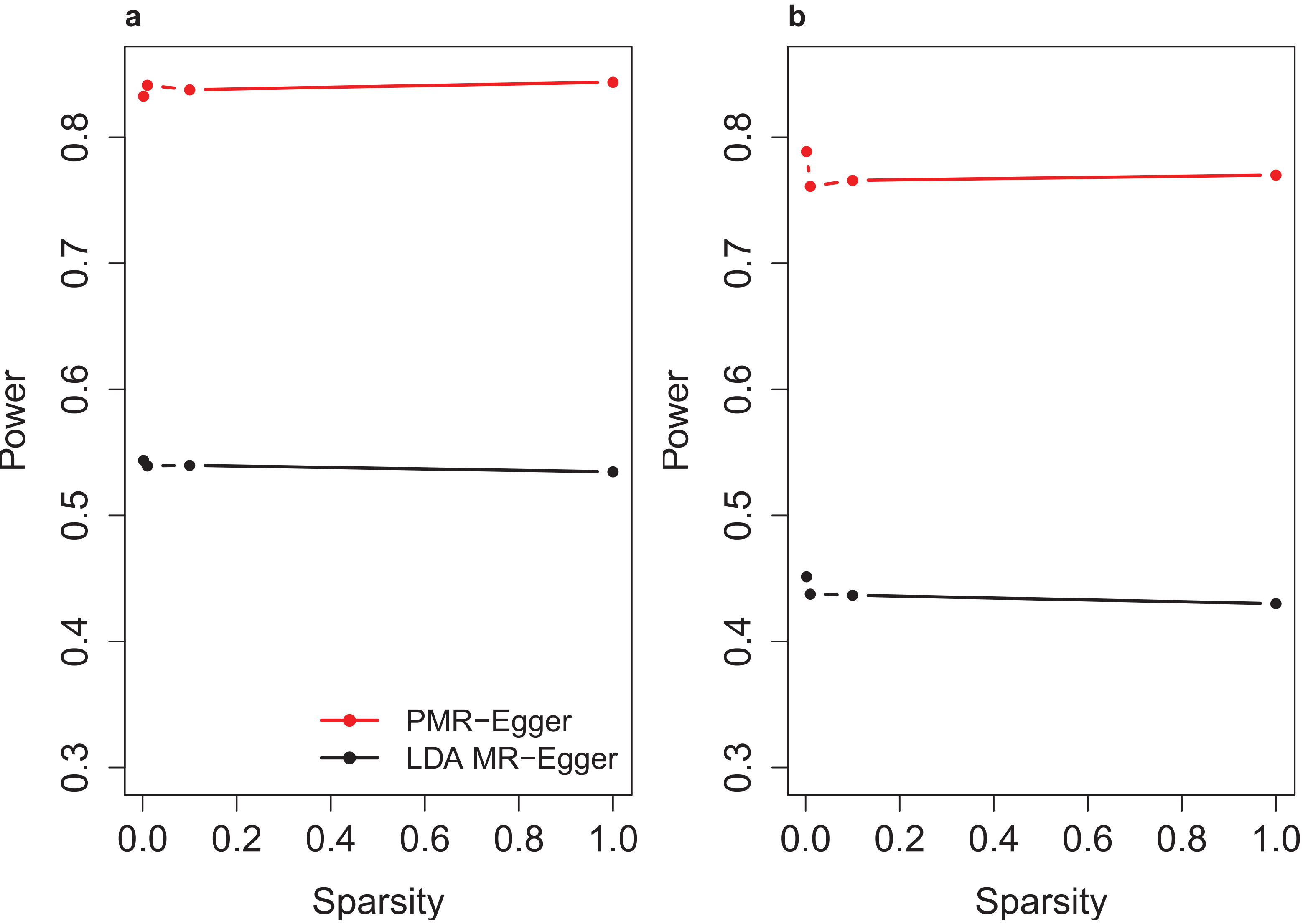


**Fig. 24** Power of different methods for identifying horizontal pleiotropic effect, in various cross-gene simulation sparse settings where only a small proportion of SNPs are associated with the gene expression level. Compared methods include PMR-Egger (red) and LDA MR-Egger (black). Power (y-axis) at a false discovery rate of 0.1 to detect the horizontal pleiotropic effect is plotted against sparsity levels, either in the absence ($PVE_{zy}$=0; a) or in the presence of causal effect ($PVE_{zy}$=0.6%; b). In terms of sparsity level (x-axis), either one SNP, 1% SNPs, 10% SNPs and 100% SNPs have non-zero effects on gene expression.

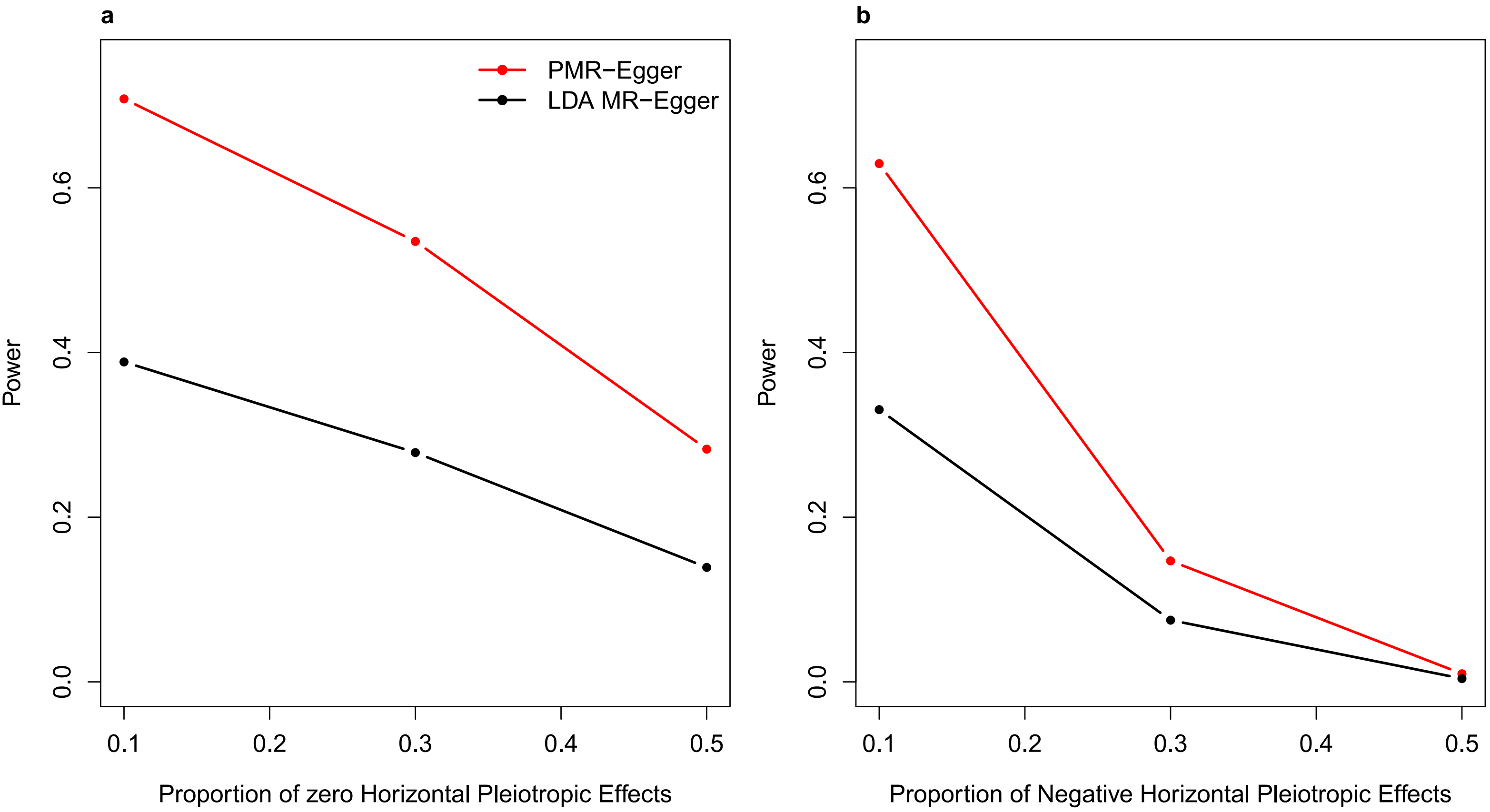


**Fig. 25** Power of different methods for identifying horizontal pleiotropic effect in cross-gene simulations under various model misspecifications of the horizontal pleiotropic effect. Compared methods include PMR-Egger (red) and LDA MR-Egger (black). On both panels, power (y-axis) to detect the horizontal pleiotropic effect is measured at a false discovery rate of 0.1 for a fixed horizontal pleiotropic effect size γ=0.001. (a) Power is plotted against the proportion of SNPs displaying zero horizontal pleiotropy effects (10%, 30%, or 50%). (b) Power is plotted against the proportion of SNPs exhibiting negative horizontal pleiotropy effects (10%, 30%, or 50%), whereas the remaining proportion of SNPs exhibiting positive effects.

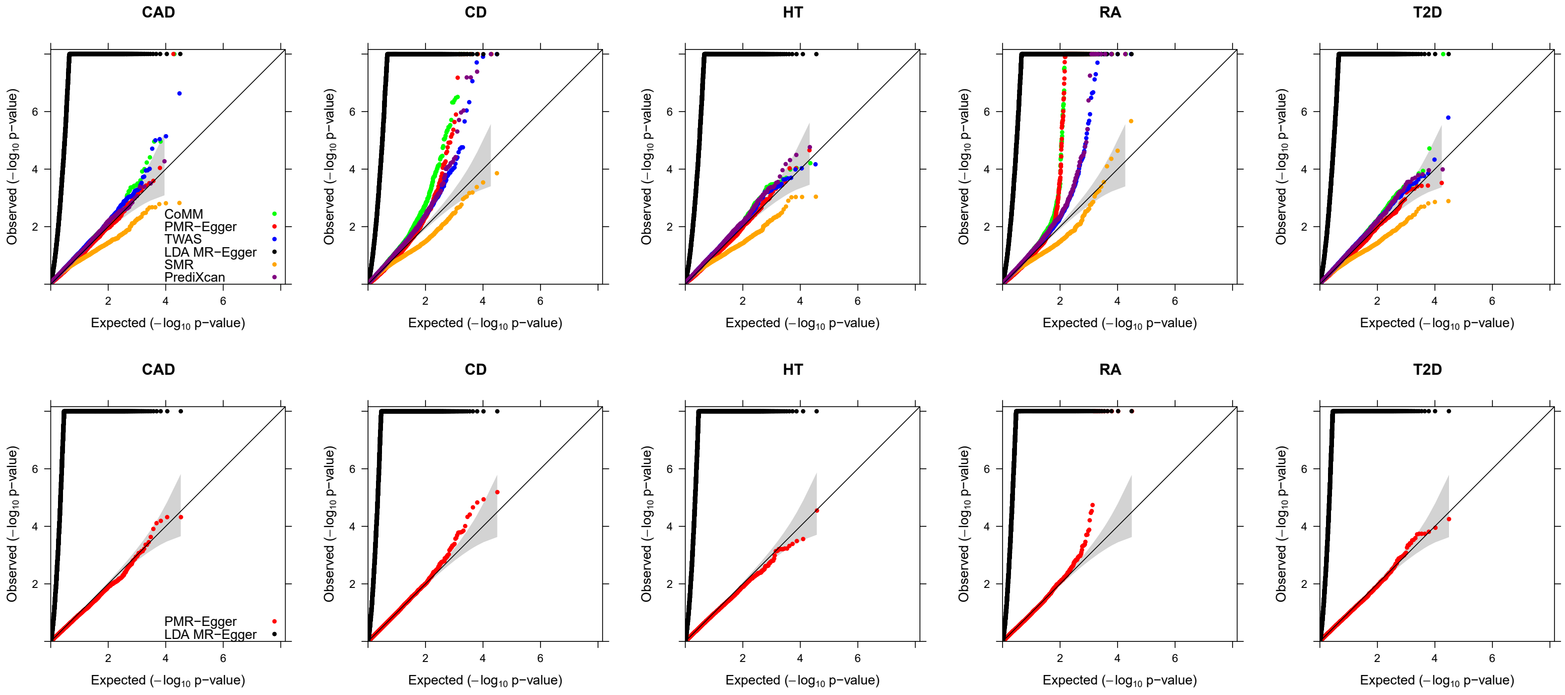


**Fig. 26** Quantile-quantile plot of -log10 p-values from different methods in the TWAS application to WTCCC. Compared methods include CoMM (green), PMR-Egger (red), TWAS (blue), LDA MR-Egger (black), SMR (orange), and PrediXcan (purple). Top panels: Quantile-quantile plot of -log10 p-values from different methods for testing the causal effect for five traits (CAD, CD, HT, RA and T2D). Bottom panels: Quantile-quantile plot of -log10 p-values from different methods for testing the horizontal pleiotropic effect for the same five traits.

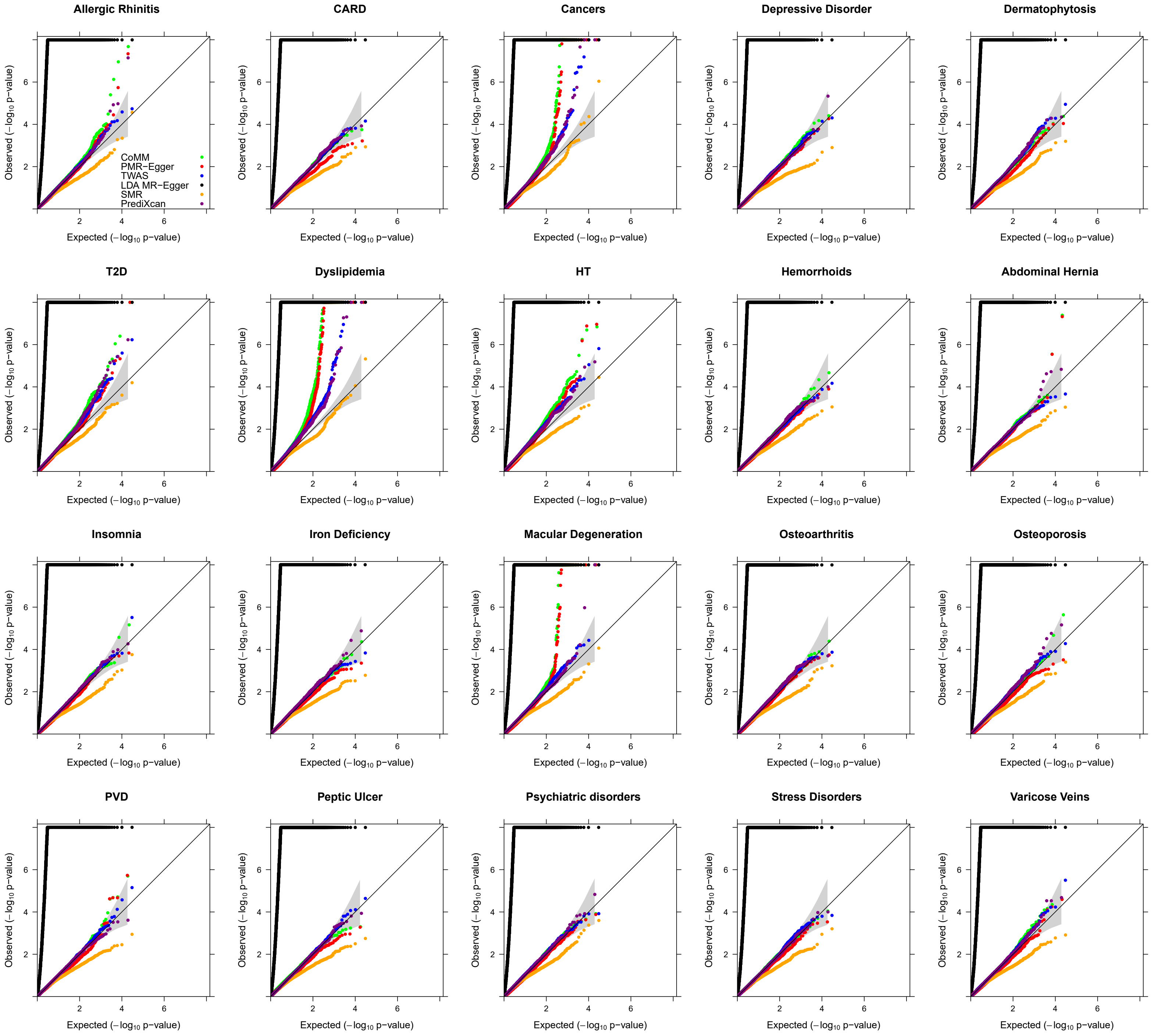


**Fig. 27** Quantile-quantile plot of -log10 p-values from different methods for testing causal effect in the TWAS application to 20 traits in GERA. Compared methods include CoMM (green), PMR-Egger (red), TWAS (blue), LDA MR-Egger (black), SMR (orange), and PrediXcan (purple).

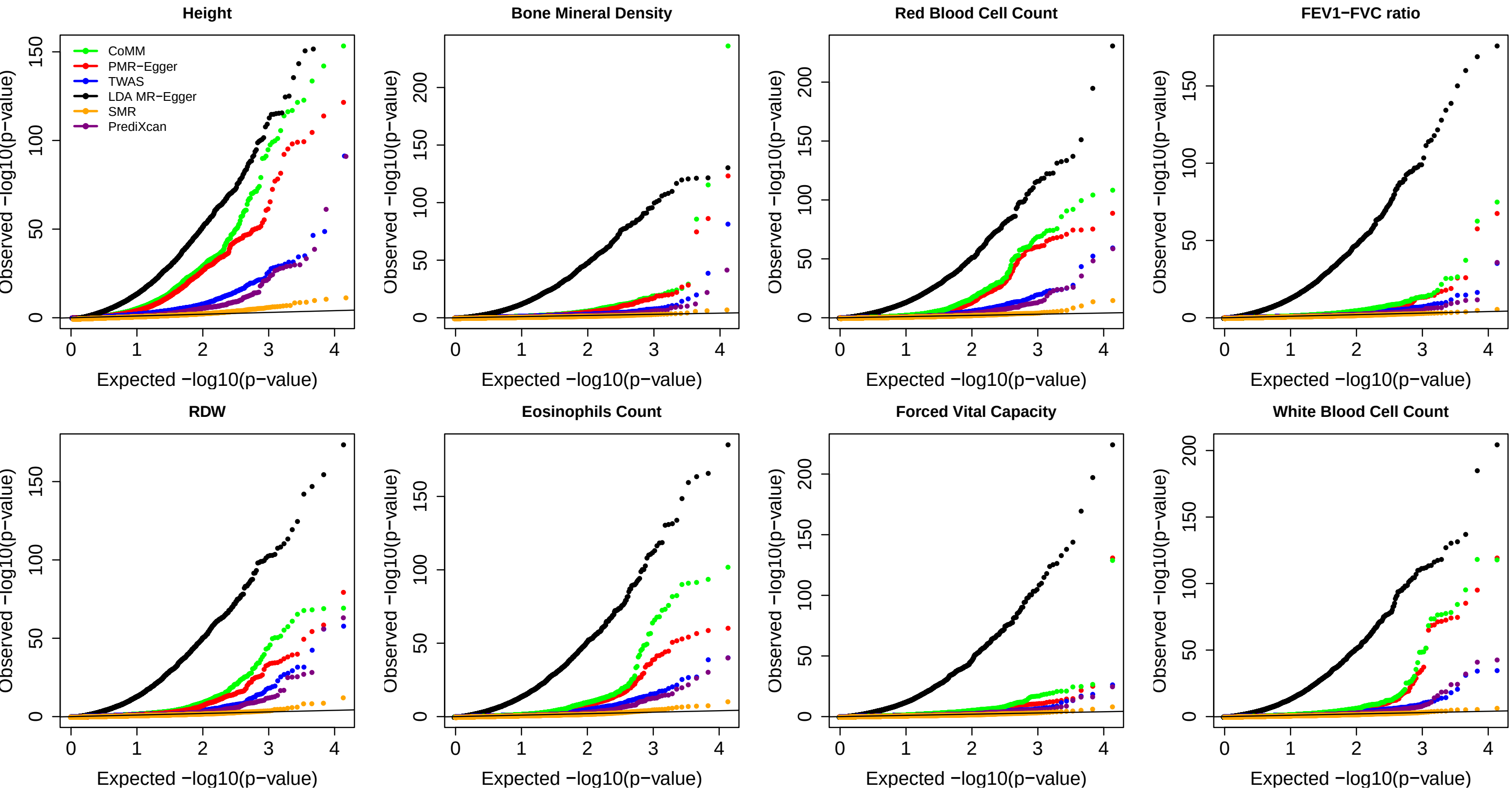


Fig. 28 Quantile-quantile plot of -log10 p-values from different methods for testing causal effect in the TWAS application to 8 traits in UK Biobank. Compared methods include CoMM (green), PMR-Egger (red), TWAS (blue), LDA MR-Egger (black), SMR (orange), and PrediXcan (purple).

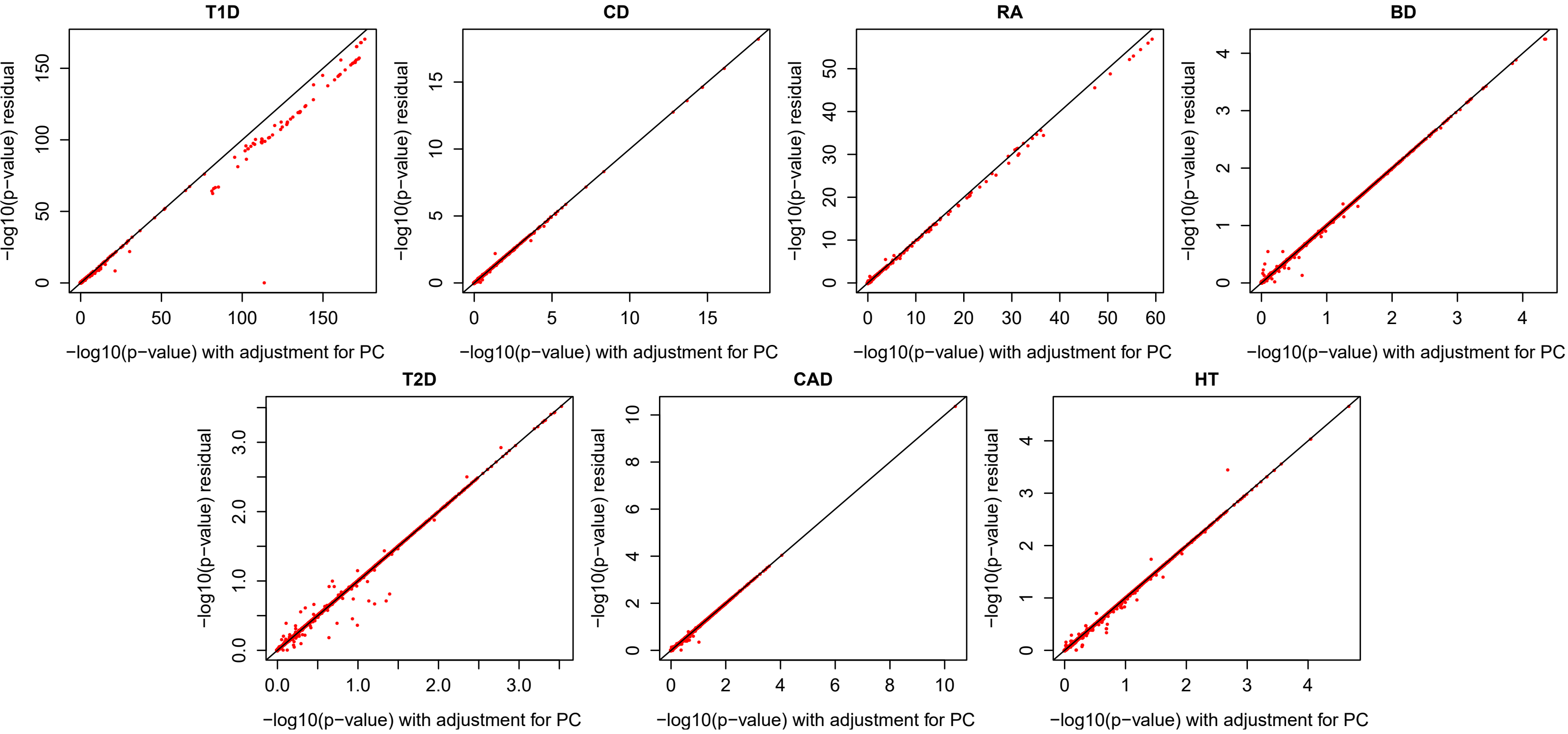


Fig. 29 The scatter plots of -log10 pvalues across genes for the 7 WTCCC traits. The pvalues from phenotype residual analysis (y-axis) is plotted against the pvalues from analysis that treating the top 10 PCs as fixed covariates.

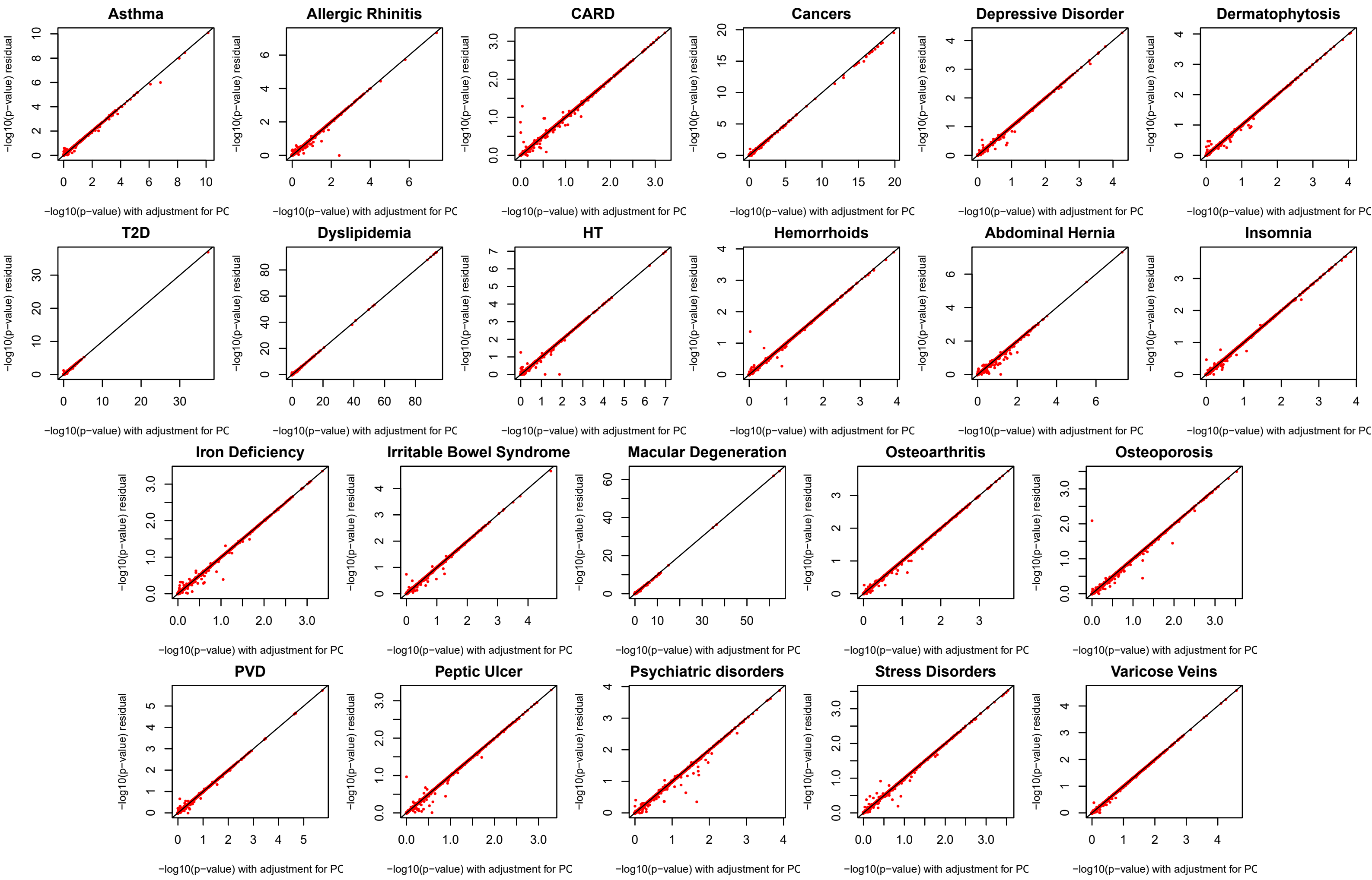


Fig. 30 The scatter plots of -log10 pvalues across genes for the 22 GERA traits. The pvalues from phenotype residual analysis (y-axis) is plotted against the pvalues from analysis that treating the top 10 PCs as fixed covariates.

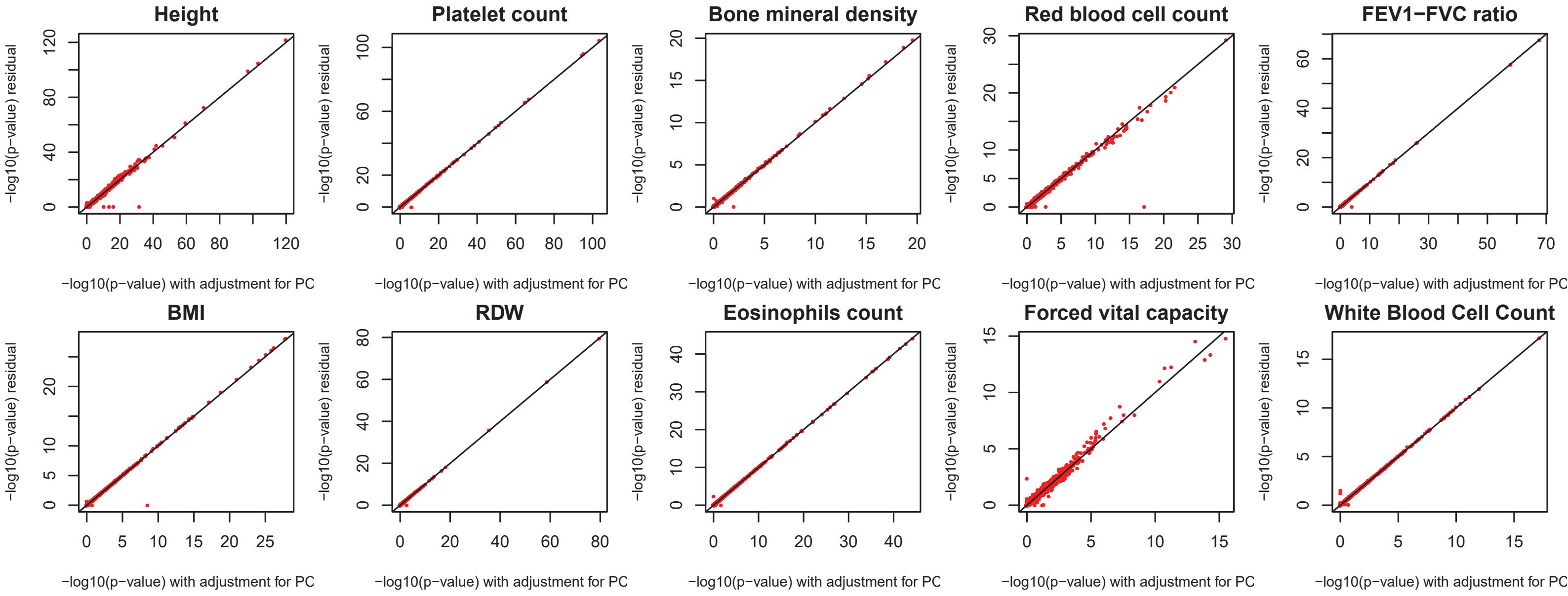


Fig. 31 The scatter plots of -log10 pvalues across genes for the 10 UKBB traits. The pvalues from phenotype residual analysis (y-axis) is plotted against the pvalues from analysis that treating the top 10 PCs as fixed covariates. Note that we only used the genes locating at first 5 chromosomes (4977 total) given the relatively high computation burden for the large sample UKBB data.

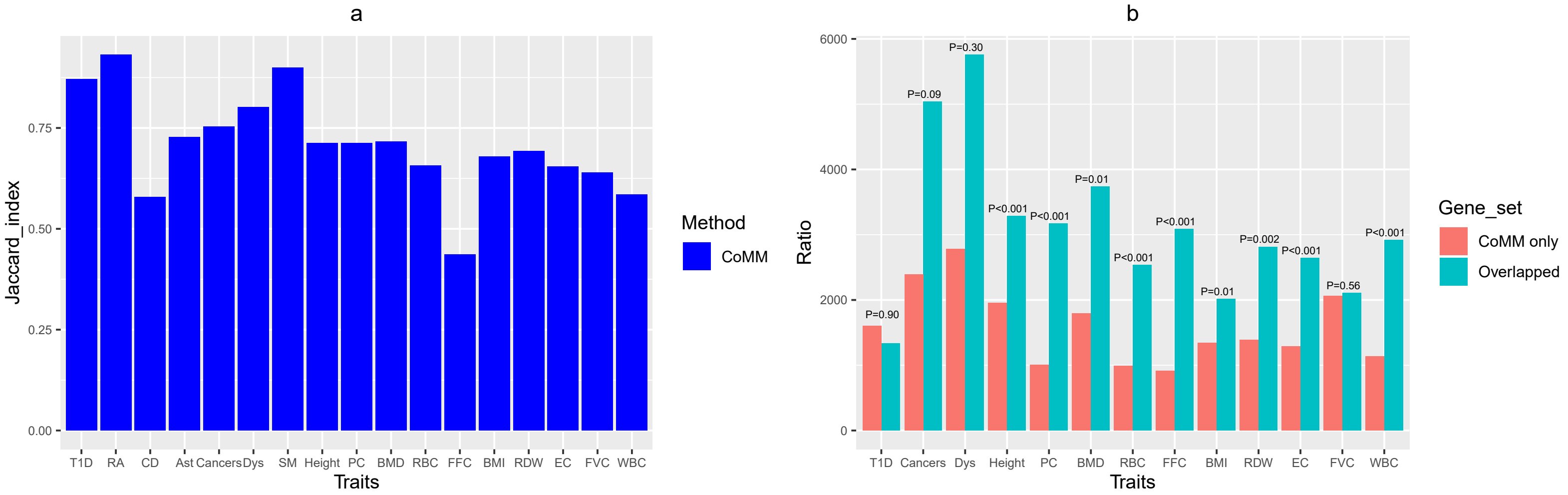


Fig. 32 Overlap of genes detected by PMR-Egger and CoMM. (a) Jaccard index measures relatively high overlap between genes detected by PMR-Egger and genes detected by CoMM. (b) Mean of the estimated $|\alpha/\gamma|$ for the set of genes that are detected by both CoMM and PMR-Egger ("overlapped"), and for the set of genes that only detected by CoMM ("CoMM only"), across 13 traits that have at least 2 detected associations. In (b), p-values are calculated based on Wilcoxon rank sum test and the traits on x-axis are: Ast, Asthma; Dys, Dyslipidemia; MD, Macular Degeneration; PC, Platelet count; BMD, Bone mineral density; RBC, Red blood cell count; FFC, FEV1-FVC ratio; EC, Eosinophils count; FVC, Forced vital capacity; WBC, White blood cell count.

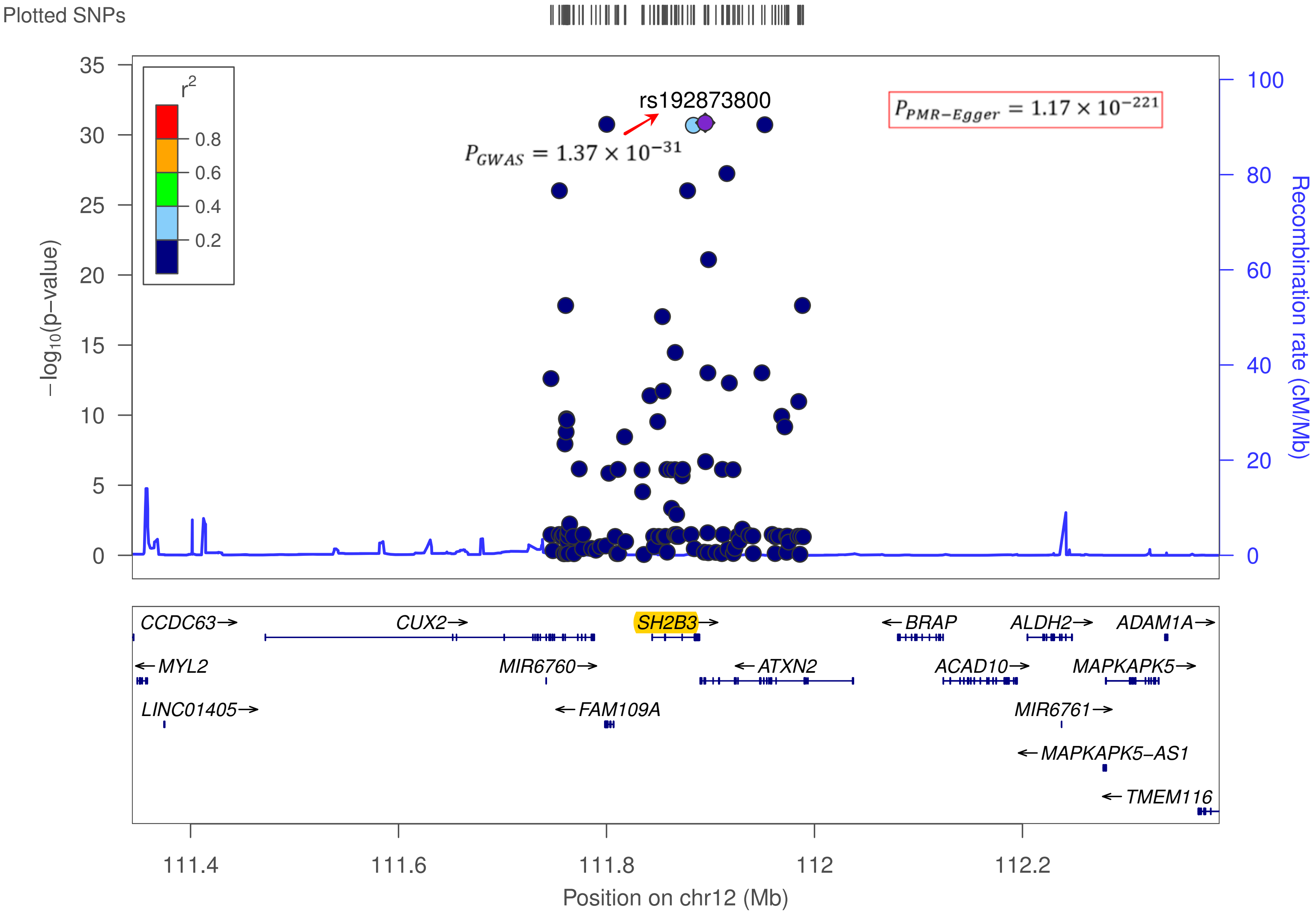


Fig. 33 Regional association plots with lead variants indicated by a purple diamond at 12q24.12. The association of an individual variant is plotted as -log10 (pvalue) against chromosomal position. The y axis shows the recombination rate estimated from 1000 Genomes Project EUR data. The identified gene SH2B3 by PMR-Egger for Platelet count is highlighted in yellow with p value showing in the top right.

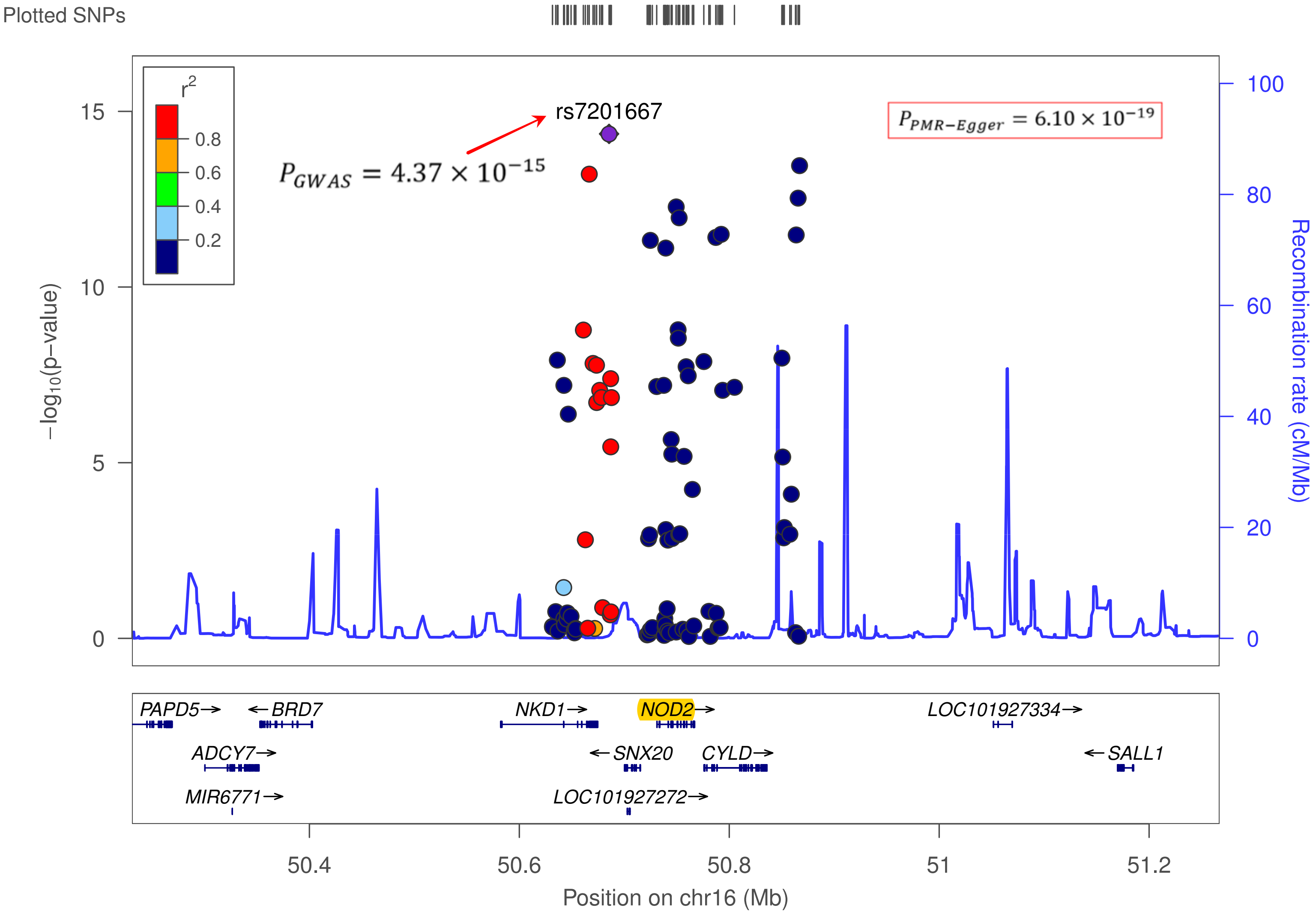


Fig. 34 Regional association plots with lead variants indicated by a purple diamond at 16q12.1. The association of an individual variant is plotted as -log10 (pvalue) against chromosomal position. The y axis shows the recombination rate estimated from 1000 Genomes Project EUR data. The identified gene NOD2 by PMR-Egger for Crohn's disease is highlighted in yellow with p value showing in the top right.

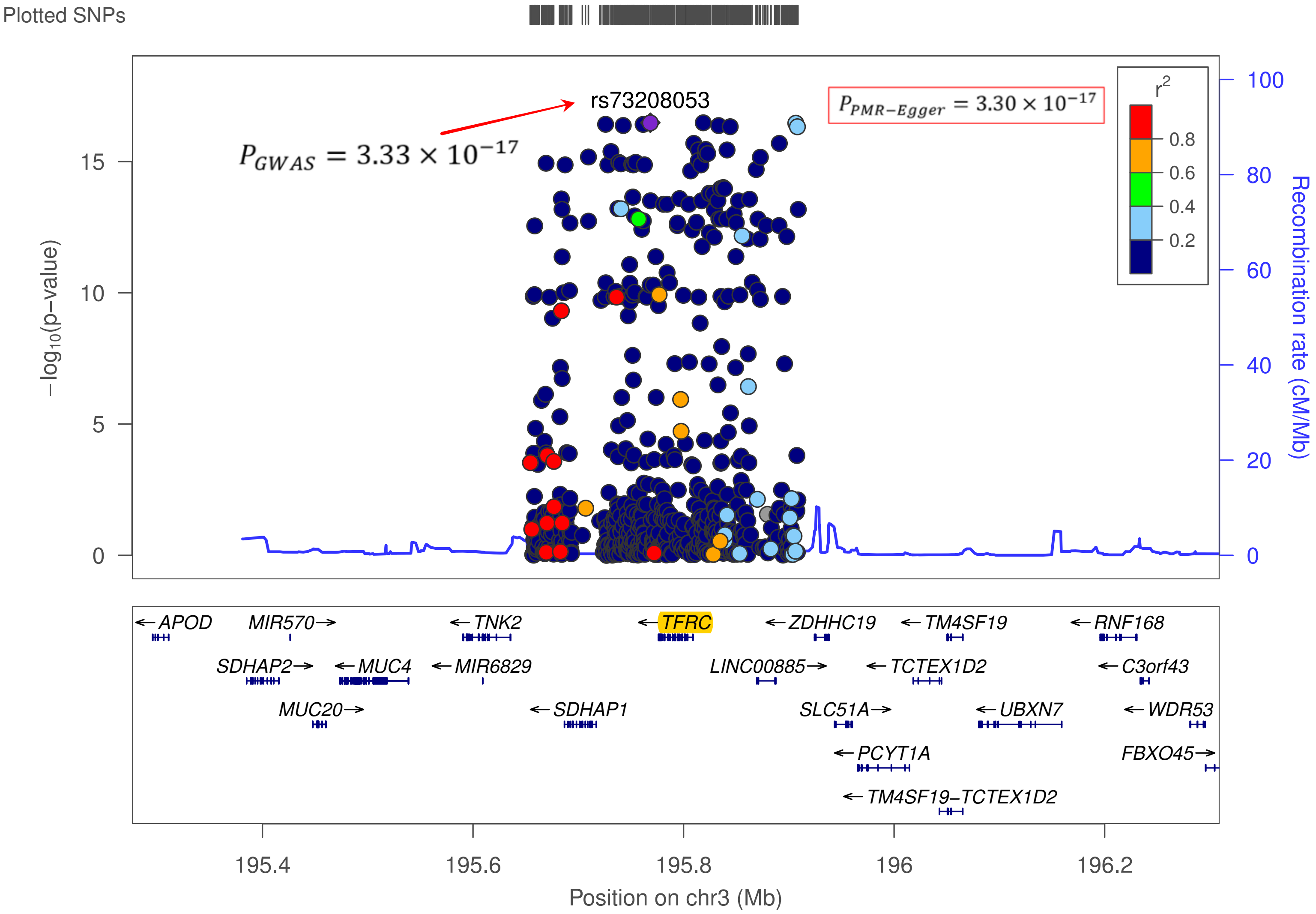


Fig. 35 Regional association plots with lead variants indicated by a purple diamond at 3q29. The association of an individual variant is plotted as -log10 (pvalue) against chromosomal position. The y axis shows the recombination rate estimated from 1000 Genomes Project EUR data. The identified gene TFRC by PMR-Egger for RDW is highlighted in yellow with p value showing in the top right.

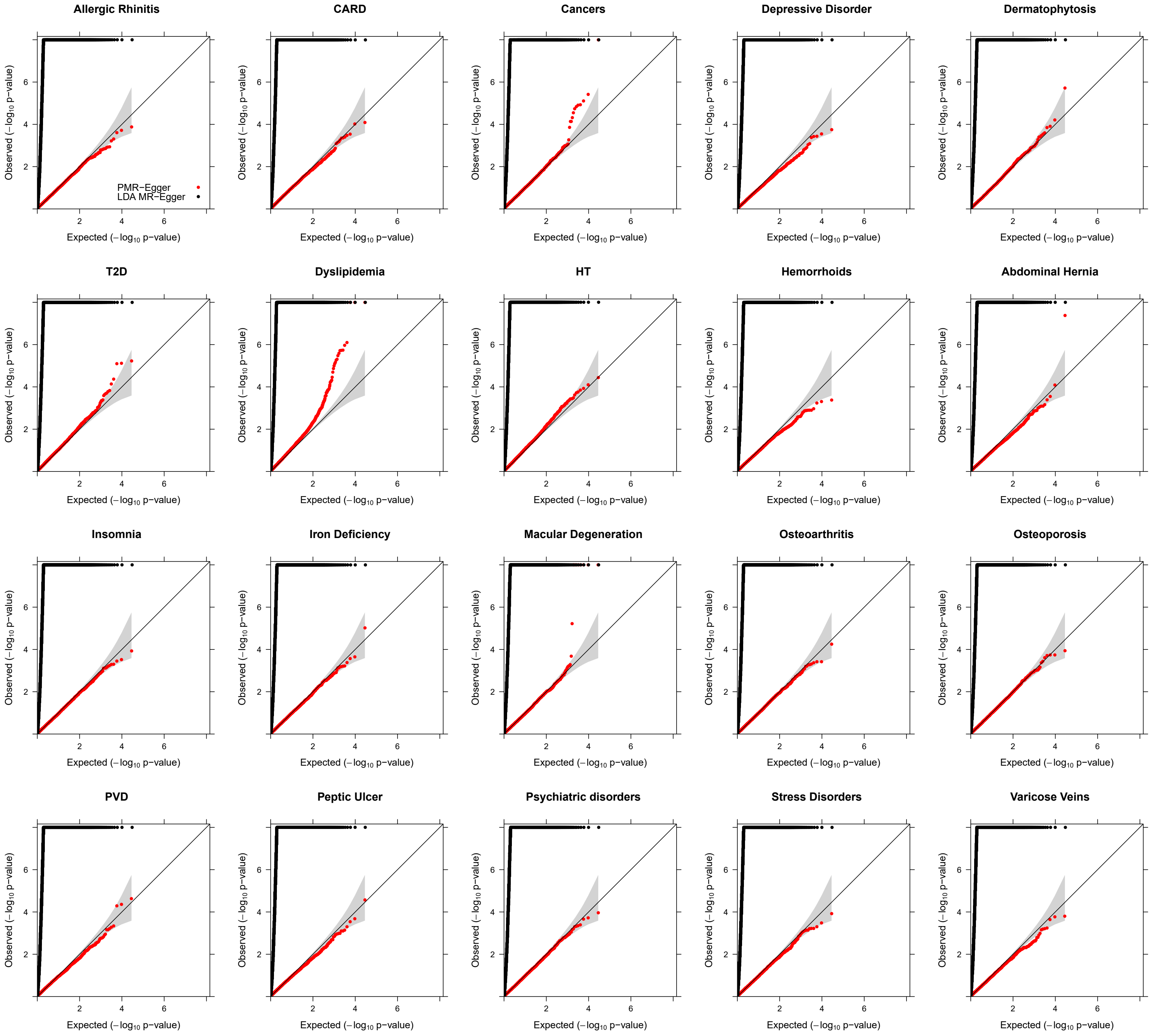


Fig. 36 Quantile-quantile plot of -log10 p-values from different methods for testing horizontal pleiotropic effect in the TWAS application to 20 traits in GERA. Compared methods include CoMM (green), PMR-Egger (red), TWAS (blue), LDA MR-Egger (black), SMR (orange), and PrediXcan (purple).

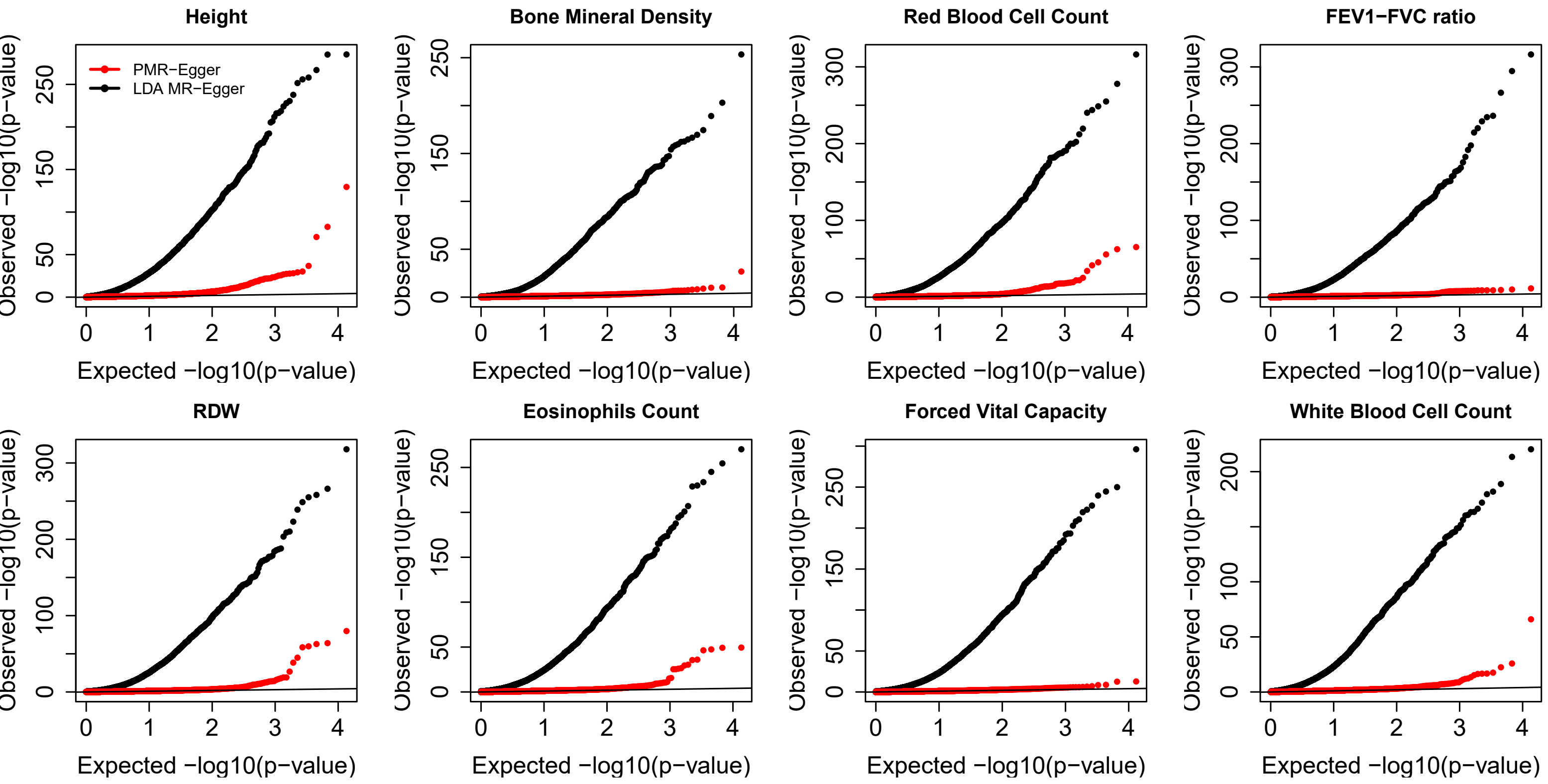


Fig. 37 Quantile-quantile plot of -log10 p-values from different methods for testing horizontal pleiotropic effect in the TWAS application to 8 traits in UK Biobank. Compared methods include CoMM (green), PMR-Egger (red), TWAS (blue), LDA MR-Egger (black), SMR (orange), and PrediXcan (purple).

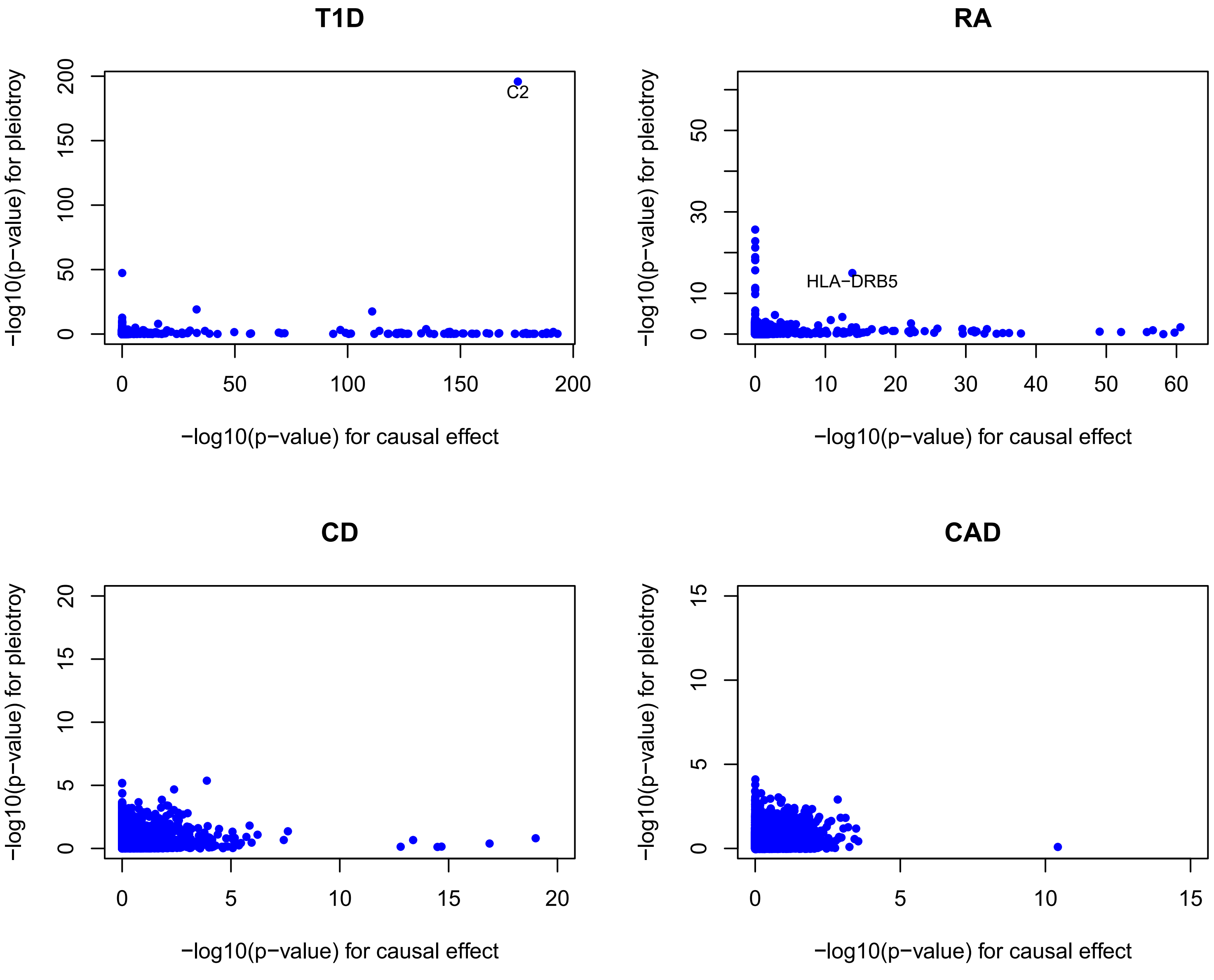


Fig. 38 Scatter plot of -log10 p-values by PMR-Egger for testing causal effect versus -log10 p-values by PMR-Egger for testing horizontal pleiotropic effect across genes for each of four traits in WTCCC. Each dot represents one gene. Only traits with at least one gene that has either a significant causal effect or a significant horizontal pleiotropic effect are displayed.

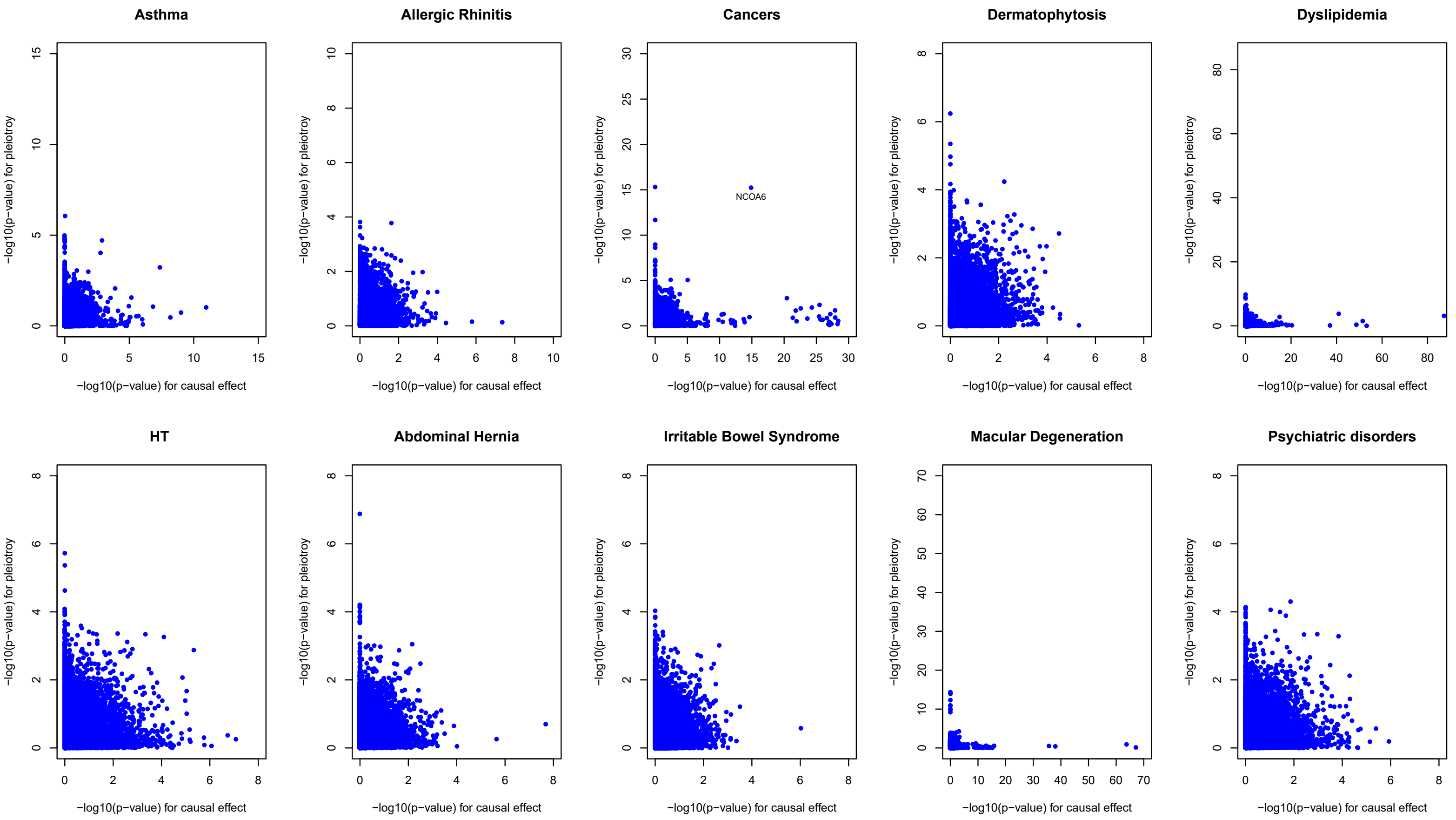


Fig. 39 Scatter plot of -log10 p-values by PMR-Egger for testing causal effect versus -log10 p-values by PMR-Egger for testing horizontal pleiotropic effect across genes for each of 10 traits in GERA. Each dot represents one gene. Only traits with at least one gene that has either a significant causal effect or a significant horizontal pleiotropic effect are displayed.

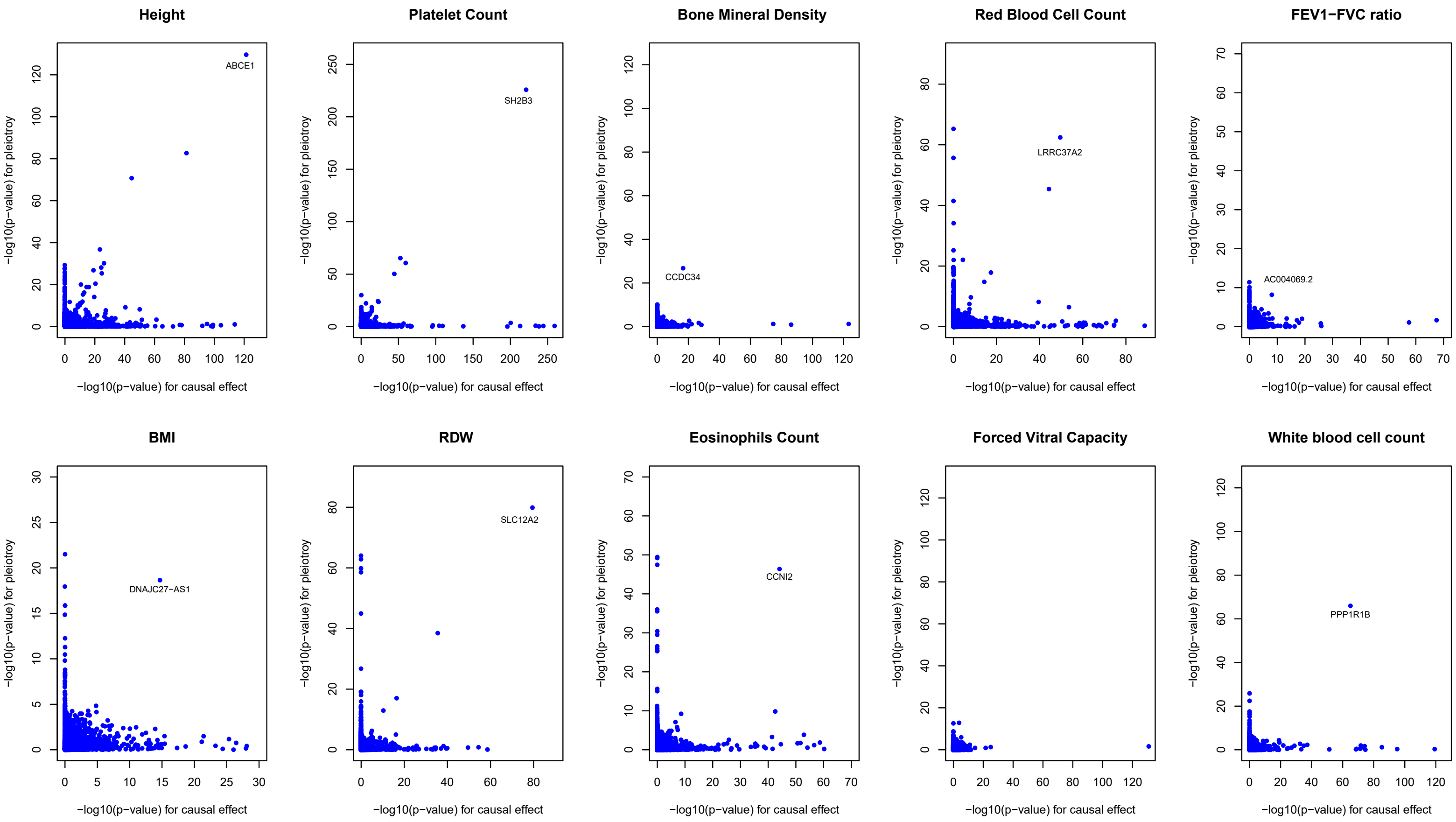


Fig. 40 Scatter plot of -log10 p-values by PMR-Egger for testing causal effect versus -log10 p-values by PMR-Egger for testing horizontal pleiotropic effect across genes for each of 10 traits in UK Biobank. Each dot represents one gene. Only traits with at least one gene that has either a significant causal effect or a significant horizontal pleiotropic effect are displayed.

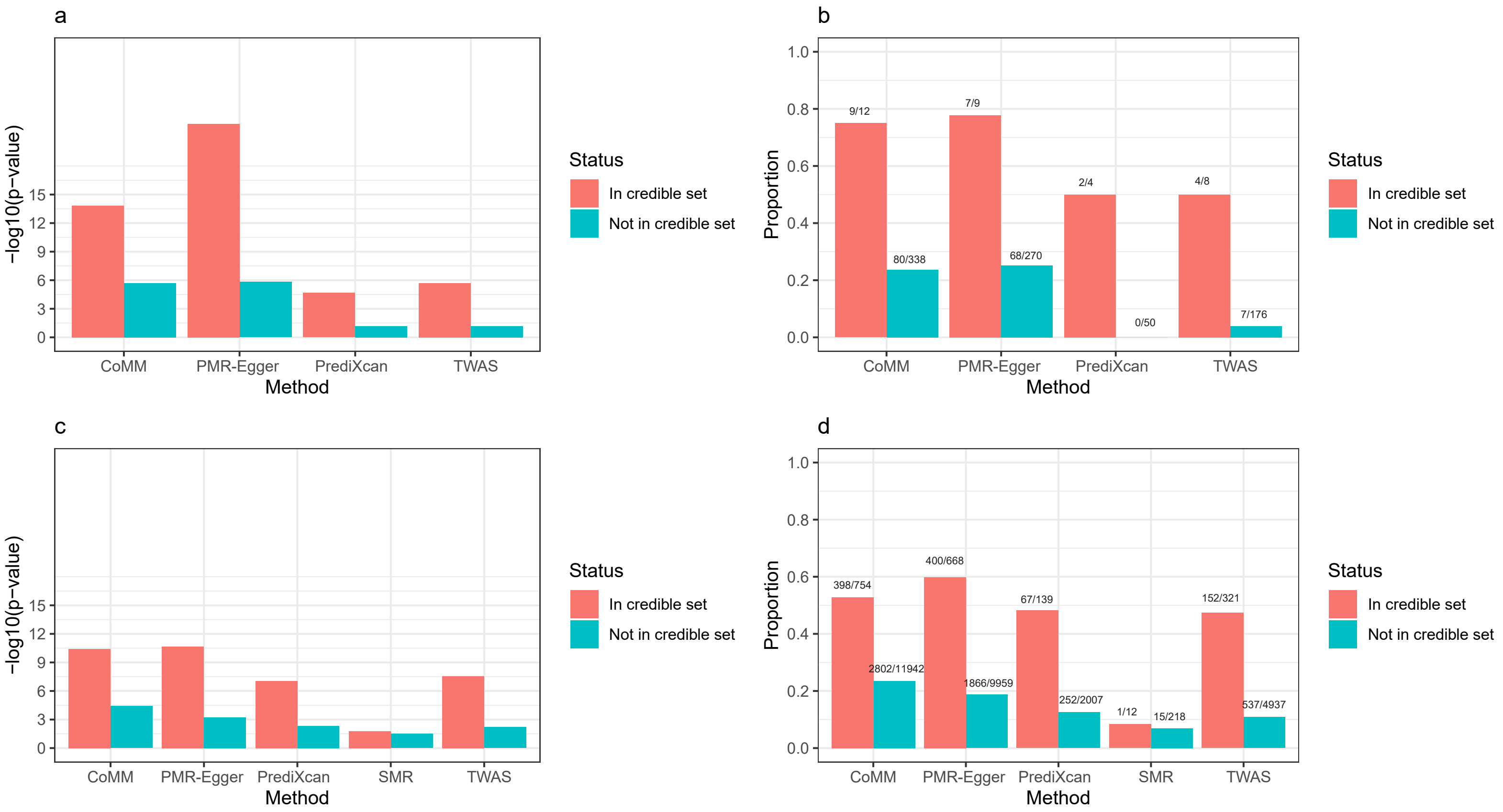


Fig. 41 Comparison of standard TWAS/MR methods with the TWAS fine-mapping method FOCUS. Results are shown for the GERA data (a, b) and the UKBiobank data (c, d). (a, c): For each method (x-axis), we computed the average -log10 p-value for genes in the 90% FOCUS credible gene set (red bars) and the average -log10 p-value for genes outside the FOCUS credible gene set (green bars). (b, d): For each method (x-axis), we also computed the proportion of genes in the 90% FOCUS credible gene set that are detected by the method (red bars) and the proportion of genes outside the FOCUS credible gene set detected by the method (green bars). The significant genes detected by each method is declared based on the Bonferroni threshold (0.05/15810). The number of significant genes/total genes in each category is also shown on top of the bars. Note that the number of total genes inside or outside the credible set is different across different methods, as FOCUS analyzes independent non-overlapping genomic regions that harbor at least one significant TWAS gene declared by the corresponding method. Overall, the PMR-Egger results are highly consistent with that of FOCUS, more so than the other methods.

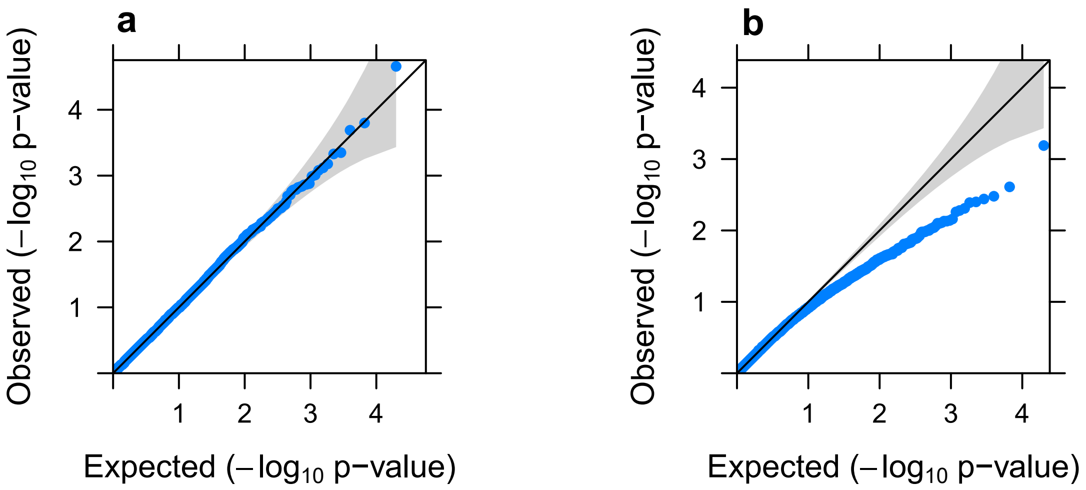


Fig. 42 Quantile-quantile plot of -log10 p-values for testing the causal effect in null simulations, under a variance component modeling assumption for the horizontal pleiotropic effects. In simulations, each $\gamma$ follows the normal distribution with variance equals 0.001/556. The p-values are calibrated if we knew the true hyper-parameters (a). However, due to the uncertainty in hyper-parameter estimates, p-values become are overly conservative when we estimate hyper-parameters (b).

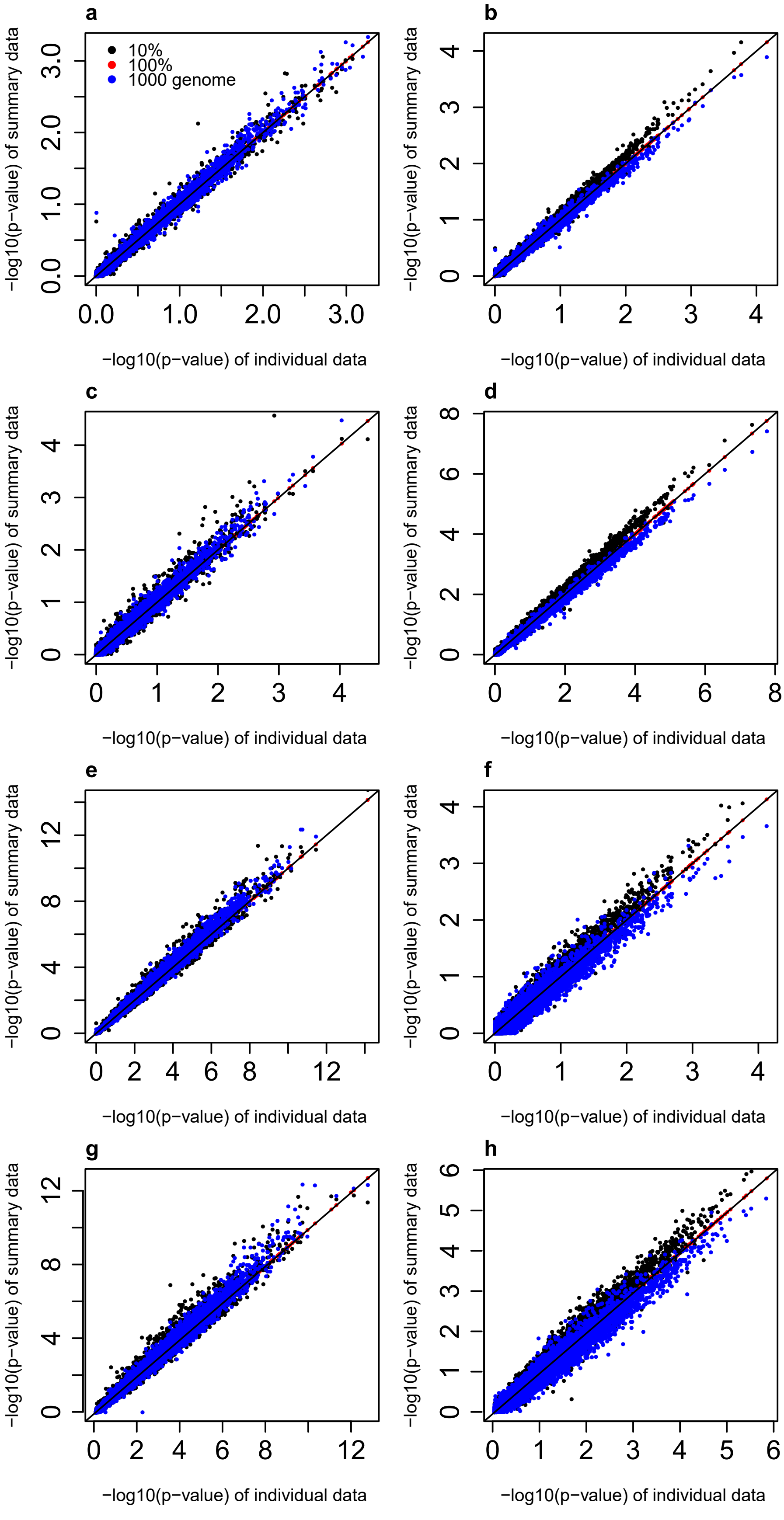


Fig. 43 Comparison of the summary-data version of PMR-Egger with the individual-data version of PMR-Egger. The LD matrix in the eQTL data is directly computed using the individual level from the eQTL study. The LD matrix in the GWAS data is computed through three different ways: by using individual-level genotypes from either all individuals in the GWAS (n=2,000; red), 10% of randomly selected individuals from the GWAS (n=200; black), or the individuals with European ancestry from the 1,000 Genomes project (n=503; blue). We examined both the causal gene association test (a, c, e, g) and the pleiotropy test (b, d, f, h). In each case, we plotted -log10 p-values from the summary-data version of PMR-Egger (y-axis) versus-log10 p-values from the summary-data version of PMR-Egger (x-axis). Simulations are performed under different causal effect and horizontal pleiotropic effect sizes: $\alpha$=0,$\gamma$=0 (a, b); $\alpha$=0,$\gamma$=0.0005 (c, d); $\alpha$=0.245,$\gamma$=0.0005 (e, f); $\alpha$=0.245, $\gamma$=0.0005 (g, h). As expected, the results from the summary-data version of PMR-Egger with that from the individual-data

### Table 1 Genomic inflation factor for three GWAS data.

| Trait | | #Gene | Causal gene effect | | | | | | Pleiotropy_effect | |
|---|---|---|---|---|---|---|---|---|---|---|
| | | | CoMM | PMR-Egger | TWAS | PrediXcan | LDA MR-Egger | SMR | PMR-Egger | LDA MR-Egger |
| WTCCC | T1D | 15584 | 1.14 | 0.95 | 1.20 | 1.21 | 18.26 | 1.06 | 1.01 | 35.53 |
| | CD | 15584 | 1.14 | 1.04 | 1.23 | 1.24 | 17.73 | 1.09 | 0.97 | 34.59 |
| | RA | 15580 | 1.13 | 0.94 | 1.22 | 1.23 | 17.60 | 1.09 | 0.96 | 34.80 |
| | BD | 15582 | 1.22 | 0.94 | 1.25 | 1.26 | 17.76 | 1.02 | 0.96 | 34.22 |
| | T2D | 15583 | 1.14 | 0.95 | 1.22 | 1.21 | 18.09 | 0.98 | 1.01 | 36.00 |
| | CAD | 15584 | 1.20 | 0.93 | 1.28 | 1.26 | 18.56 | 0.94 | 0.93 | 34.00 |
| | HT | 15581 | 1.23 | 0.96 | 1.31 | 1.31 | 18.21 | 0.97 | 0.96 | 34.11 |
| GERA | Asthma | 15162 | 1.25 | 1.05 | 1.10 | 1.06 | 33.60 | 0.98 | 1.09 | 72.19 |
| | Allergic Rhinitis | 15154 | 1.11 | 0.95 | 1.06 | 1.05 | 32.13 | 0.99 | 0.96 | 72.18 |
| | CARD | 15146 | 1.12 | 0.96 | 1.09 | 1.11 | 33.74 | 1.02 | 0.99 | 72.19 |
| | Cancers | 15135 | 1.26 | 1.13 | 1.09 | 1.07 | 34.36 | 1.02 | 1.00 | 72.19 |
| | Depressive Disorder | 15131 | 0.95 | 1.02 | 1.05 | 1.02 | 33.95 | 0.96 | 0.93 | 70.01 |
| | Dermatophytosis | 15128 | 0.95 | 0.93 | 1.01 | 1.02 | 33.64 | 0.95 | 1.05 | 72.18 |
| | T2D | 15123 | 1.26 | 0.98 | 1.15 | 1.10 | 34.23 | 1.05 | 1.02 | 72.19 |
| | Dyslipidemia | 15119 | 1.13 | 1.07 | 1.14 | 1.16 | 32.89 | 1.05 | 1.06 | 72.19 |
| | HT | 15117 | 1.62 | 0.99 | 1.16 | 1.15 | 34.74 | 1.05 | 1.04 | 72.19 |
| | Hemorrhoids | 15110 | 1.02 | 0.95 | 1.05 | 1.03 | 33.40 | 0.97 | 0.95 | 72.19 |
| | Abdominal Hernia | 15104 | 0.99 | 1.01 | 1.03 | 1.03 | 33.65 | 0.97 | 1.03 | 72.19 |
| | Insomnia | 15094 | 0.94 | 0.96 | 1.06 | 1.07 | 33.71 | 0.96 | 1.04 | 72.19 |
| | Iron Deficiency | 15088 | 1.09 | 0.93 | 1.05 | 1.05 | 33.54 | 0.98 | 0.97 | 72.19 |
| | Irritable Bowel Syndrome | 15080 | 0.97 | 0.95 | 1.02 | 1.01 | 33.02 | 0.96 | 1.04 | 72.18 |
| | Macular Degeneration | 15075 | 1.18 | 1.08 | 1.03 | 1.03 | 33.40 | 0.99 | 1.04 | 72.19 |
| | Osteoarthritis | 15072 | 0.94 | 0.93 | 1.08 | 1.06 | 33.51 | 1.01 | 0.96 | 72.19 |
| | Osteoporosis | 15069 | 1.08 | 0.92 | 1.04 | 1.03 | 32.33 | 0.97 | 1.05 | 72.19 |
| | PVD | 15065 | 1.11 | 0.93 | 1.01 | 1.02 | 33.30 | 0.97 | 1.05 | 72.19 |
| | Peptic Ulcer | 15061 | 0.96 | 0.94 | 1.03 | 1.03 | 33.64 | 0.95 | 0.93 | 72.18 |
| | Psychiatric disorders | 15057 | 0.94 | 0.95 | 1.07 | 1.07 | 33.89 | 1.01 | 1.01 | 69.82 |
| | Stress Disorders | 15055 | 1.07 | 0.92 | 0.98 | 1.00 | 32.47 | 0.96 | 0.92 | 72.19 |
| | Varicose Veins | 15051 | 1.01 | 0.96 | 1.06 | 1.02 | 33.34 | 0.94 | 0.93 | 72.19 |
| UKBiobank | Height | 15761 | 1.61 | 1.18 | 2.17 | 1.46 | 16.65 | 1.71 | 1.71 | 29.85 |
| | Platelet Count | 15765 | 1.87 | 1.31 | 1.46 | 1.26 | 15.25 | 1.41 | 1.45 | 26.07 |
| | Bone Mineral Density | 15761 | 1.45 | 1.13 | 1.27 | 1.1 | 10.48 | 1.13 | 1.13 | 17.75 |
| | Red Blood Cell Count | 15751 | 1.71 | 1.18 | 1.64 | 1.22 | 14.12 | 1.36 | 1.38 | 23.45 |
| | FEV1-FVC ratio | 15747 | 1.59 | 1.17 | 1.3 | 1.11 | 12.91 | 1.19 | 1.19 | 19.48 |
| | BMI | 14633 | 1.68 | 1.12 | 1.38 | 1.16 | 14.59 | 1.29 | 1.35 | 25.95 |
| | RDW | 15744 | 1.65 | 1.16 | 1.42 | 1.09 | 13.93 | 1.18 | 1.22 | 22.59 |
| | Eosinophils Count | 15740 | 1.9 | 1.34 | 1.46 | 1.14 | 14.33 | 1.23 | 1.31 | 22.42 |
| | Forced Vital Capacity | 15735 | 1.78 | 1.3 | 1.38 | 1.13 | 13.01 | 1.25 | 1.14 | 22.17 |
| | White blood cell count | 15729 | 1.76 | 1.25 | 1.38 | 1.12 | 13.93 | 1.17 | 1.32 | 22.89 |

**Table 2 Number of significant genes for three GWAS data under Bonferroni correction.**

| Trait | | #Gene | Causal gene effect | | | | | | Pleiotropy_effect | |
|---|---|---|---|---|---|---|---|---|---|---|
| | | | CoMM | PMR-Egger | TWAS | PrediXcan | LDA MR-Egger | SMR | PMR-Egger | LDA MR-Egger |
| **WTCCC** | T1D | 15584 | 151 | 137 | 62 | 42 | 4539 | 4 | 11 | 6860 |
| | CD | 15584 | 14 | 10 | 7 | 7 | 4586 | 0 | 0 | 6797 |
| | RA | 15580 | 81 | 81 | 16 | 10 | 4601 | 1 | 11 | 6773 |
| | BD | 15582 | 0 | 0 | 0 | 0 | 4712 | 0 | 0 | 6890 |
| | T2D | 15583 | 1 | 0 | 1 | 0 | 4565 | 0 | 0 | 6851 |
| | CAD | 15584 | 1 | 1 | 1 | 1 | 4953 | 0 | 0 | 7206 |
| | HT | 15581 | 0 | 0 | 0 | 0 | 5512 | 0 | 0 | 8020 |
| **GERA** | Asthma | 15162 | 6 | 5 | 3 | 4 | 6468 | 4 | 0 | 8928 |
| | Allergic Rhinitis | 15154 | 3 | 2 | 0 | 1 | 6356 | 0 | 0 | 9010 |
| | CARD | 15146 | 0 | 0 | 0 | 0 | 6358 | 0 | 0 | 8943 |
| | Cancers | 15135 | 34 | 28 | 8 | 4 | 6500 | 1 | 1 | 9017 |
| | Depressive Disorder | 15131 | 0 | 0 | 0 | 0 | 6480 | 0 | 0 | 8752 |
| | Dermatophytosis | 15128 | 0 | 0 | 0 | 0 | 6465 | 0 | 1 | 8876 |
| | T2D | 15123 | 3 | 1 | 2 | 1 | 6529 | 0 | 0 | 8979 |
| | Dyslipidemia | 15119 | 63 | 57 | 11 | 8 | 6392 | 0 | 10 | 9139 |
| | HT | 15117 | 4 | 3 | 1 | 0 | 6564 | 0 | 0 | 9042 |
| | Hemorrhoids | 15110 | 0 | 0 | 0 | 0 | 6422 | 0 | 0 | 8801 |
| | Abdominal Hernia | 15104 | 2 | 2 | 0 | 0 | 6508 | 0 | 1 | 8894 |
| | Insomnia | 15094 | 0 | 0 | 1 | 0 | 6483 | 0 | 0 | 8884 |
| | Iron Deficiency | 15088 | 0 | 0 | 0 | 0 | 6451 | 0 | 0 | 8855 |
| | Irritable Bowel Syndrome | 15080 | 0 | 0 | 0 | 0 | 6384 | 0 | 0 | 8918 |
| | Macular Degeneration | 15075 | 32 | 28 | 1 | 2 | 6458 | 0 | 8 | 8900 |
| | Osteoarthritis | 15072 | 0 | 0 | 0 | 0 | 6460 | 0 | 0 | 8895 |
| | Osteoporosis | 15069 | 1 | 0 | 0 | 0 | 6404 | 0 | 0 | 8797 |
| | PVD | 15065 | 1 | 1 | 0 | 0 | 6386 | 0 | 0 | 8909 |
| | Peptic Ulcer | 15061 | 0 | 0 | 0 | 0 | 6449 | 0 | 0 | 8888 |
| | Psychiatric disorders | 15057 | 0 | 0 | 0 | 0 | 6443 | 0 | 0 | 8693 |
| | Stress Disorders | 15055 | 0 | 0 | 0 | 0 | 6318 | 0 | 0 | 8834 |
| | Varicose Veins | 15051 | 0 | 0 | 1 | 0 | 6464 | 0 | 0 | 8945 |
| **UKBiobank** | Height | 15761 | 1295 | 1008 | 285 | 172 | 3802 | 26 | 170 | 5509 |
| | Platelet Count | 15765 | 645 | 499 | 155 | 101 | 3606 | 13 | 123 | 5171 |
| | Bone Mineral Density | 15761 | 146 | 108 | 29 | 14 | 2933 | 3 | 15 | 4036 |
| | Red Blood Cell Count | 15751 | 497 | 353 | 145 | 81 | 3417 | 8 | 92 | 4928 |
| | FEV1-FVC ratio | 15747 | 97 | 51 | 28 | 12 | 3258 | 1 | 27 | 4520 |
| | BMI | 14633 | 240 | 173 | 47 | 31 | 3533 | 2 | 28 | 4971 |
| | RDW | 15744 | 247 | 183 | 78 | 51 | 3413 | 7 | 57 | 4835 |
| | Eosinophils Count | 15740 | 295 | 223 | 111 | 48 | 3548 | 8 | 57 | 4800 |
| | Forced Vital Capacity | 15735 | 118 | 82 | 26 | 16 | 3243 | 3 | 11 | 4574 |
| | White blood cell count | 15729 | 182 | 116 | 54 | 34 | 3363 | 4 | 46 | 4793 |

**Table 3 Summary of gene-based fine mapping results in three GWAS data by FOCUS**

| Trait | | #Analyzed regions | #Analyzed regions with multivariate TWAS- significant gene | #TWAS-significant gene in analyzed regions | #genes in analyzed regions | #genes in 90% credible gene sets |
|---|---|---|---|---|---|---|
| **WTCCC** | T1D | 7 | 6 | 28 | 210 | 4 |
| | CD | 3 | 2 | 7 | 24 | 1 |
| | RA | 2 | 2 | 11 | 42 | 1 |
| | CAD | 1 | 1 | 1 | 12 | 0 |
| **GERA** | Allergic Rhinitis | 1 | 1 | 2 | 5 | 0 |
| | Cancers | 4 | 4 | 27 | 81 | 4 |
| | Dyslipidemia | 9 | 9 | 42 | 167 | 8 |
| | Abdominal Hernia | 1 | 1 | 2 | 9 | 1 |
| | Macular Degeneration | 2 | 2 | 4 | 15 | 0 |
| **UKBiobank** | Height | 169 | 150 | 946 | 3559 | 247 |
| | Platelet Count | 86 | 75 | 462 | 1991 | 140 |
| | Bone Mineral Density | 28 | 24 | 97 | 461 | 32 |
| | Red Blood Cell Count | 56 | 50 | 343 | 1426 | 89 |
| | FEV1-FVC ratio | 15 | 13 | 47 | 309 | 20 |
| | BMI | 33 | 30 | 171 | 923 | 62 |
| | RDW | 45 | 33 | 199 | 1196 | 76 |
| | Eosinophils Count | 48 | 38 | 210 | 1223 | 54 |
| | Forced Vital Capacity | 15 | 14 | 83 | 422 | 30 |
| | White blood cell count | 20 | 19 | 127 | 495 | 29 |

Summary of fine mapping results from FOCUS in three GWAS data sets. For each trait in turn (listed in rows), we performed FOCUS analysis on independent non-overlapping genomic regions that harbor on least one genome-wide-significant SNP ($p < 5 \times 10^{-8}$) and at least a significant TWAS gene under Bonferroni correction. In this table, the significant TWAS genes are declared based on PMR-Egger results; the other parallel FOCUS analyses based on the other corresponding MR methods are not shown in this table. The table lists the number of analyzed regions (3$^{rd}$ column), number of analyzed regions that contain multiple significant genes detected by PMR-Egger (4$^{th}$ column), number of significant TWAS genes in the analyzed regions (5$^{th}$ column), the number of total genes analyzed in these regions (6$^{th}$ column) and the number of genes in the 90% credible set by FOCUS (7$^{th}$ column).
